# A fast and efficient colocalization algorithm for identifying shared genetic risk factors across multiple traits

**DOI:** 10.1101/592238

**Authors:** Christopher N Foley, James R Staley, Philip G Breen, Benjamin B Sun, Paul D W Kirk, Stephen Burgess, Joanna M M Howson

## Abstract

Genome-wide association studies (GWAS) have identified thousands of genomic regions affecting complex diseases. The next challenge is to elucidate the causal genes and mechanisms involved. One approach is to use statistical colocalization to assess shared genetic aetiology across multiple related traits (e.g. molecular traits, metabolic pathways and complex diseases) to identify causal pathways, prioritize causal variants and evaluate pleiotropy. We propose HyPrColoc (Hypothesis Prioritisation in multi-trait Colocalization), an efficient deterministic Bayesian algorithm using GWAS summary statistics that can detect colocalization across vast numbers of traits simultaneously (e.g. 100 traits can be jointly analysed in around 1 second). We performed a genome-wide multi-trait colocalization analysis of coronary heart disease (CHD) and fourteen related traits. HyPrColoc identified 43 regions in which CHD colocalized with ≥1 trait, including 5 potentially new CHD loci. Across the 43 loci, we further integrated gene and protein expression quantitative trait loci to identify candidate causal genes.

## Introduction

Genome wide association studies (GWAS) have identified thousands of genomic loci associated with complex traits and diseases (https://www.ebi.ac.uk/gwas/). However, identification of the causal mechanisms underlying these associations and subsequent biological insights have not been as forthcoming, due to issues such as linkage disequilibrium (LD) and incomplete genomic coverage. One approach to aid biological insight following GWAS is to make use of functional data. For example, candidate causal genes can be proposed when the overlap in association signals between a complex trait and functional data (*e.g*. gene expression) is a consequence of both traits sharing a causal variant, *i.e*. the association signals for both traits colocalize. The abundance of significant associations identified by GWAS means that chance overlap between association signals for different traits is likely^1^. Consequently, overlap does not by itself allow us to identify causal variants^1,2^. Statistical colocalization methodologies seek to resolve this. By constructing a formal statistical model, colocalization approaches have been successful in identifying whether a molecular trait (e.g. gene expression) and a disease trait share a causal variant in a genomic region^3–7^, and potentially prioritise a candidate causal gene. Recently it has been proposed that colocalization methodologies can be further enhanced by integrating additional information from *multiple* intermediate traits linked to disease, e.g. protein expression, metabolite levels^8^. The underlying hypothesis of *multi-trait colocalization* is that if a variant is associated with multiple related traits then this provides stronger evidence that the variant may be causal^8^. Thus, multi-trait colocalization aims to increase power to identify causal variants. We show that by using multi-level functional datasets in this way can reveal candidate causal genes and pathways underpinning complex disease.

A number of statistical methods have been developed to assess whether association signals across a *pair* of traits colocalize^3–7^. These methods predominantly assess colocalization between a pair of traits using individual participant data^9,10^, limiting their applicability. In contrast, the COLOC algorithm uses GWAS summary statistics^2^. This approach works by systematically exploring putative “causal configurations”, where each configuration locates a causal variant for one or both traits, under the assumption that there is at most one causal variant per trait. COLOC was recently extended to the multi-trait framework, MOLOC^8^. The authors achieved a 1.5-fold increase in candidate causal gene identification when a third relevant trait was included in colocalization analyses relative to results from two traits. However, the approach is computationally impractical beyond 4 traits due to prohibitive computational complexity arising from the exponential growth in the number of causal configurations that must be explored with each additional trait analysed.

Here we present a computationally efficient method, *Hypothesis Prioritisation in multi-trait Colocalization* (HyPrColoc), to identify colocalized association signals using summary statistics on large numbers of traits. The approach extends the underlying methodology of COLOC and MOLOC. Our major result is that the posterior probability of colocalization at a single causal variant can be accurately approximated by enumerating only a small number of putative causal configurations. Moreover, HyPrColoc is able to identify *subsets* (which we refer to as *clusters*) of traits which colocalize at *distinct* causal variants in the genomic locus by employing a novel branch and bound divisive clustering algorithm. We applied HyPrColoc genome-wide to coronary heart disease (CHD) and many related traits^11,12^, to identify genetic risk loci shared across these traits.

## Results

### Overview

HyPrColoc is a Bayesian method for identifying shared genetic associations between complex traits in a particular gene region using summary GWAS results. HyPrColoc provides two principal novelties: (i) Efficient computation of the posterior probability that all *m* traits share a causal variant (which we refer to as the posterior probability of full colocalization, PPFC); and (ii) partitioning of traits into clusters, such that each cluster comprises traits sharing a causal variant. HyPrColoc only requires regression coefficients and their corresponding standard errors from summary GWAS (for binary traits these can be on the log-odds scale, **Methods**). The approach makes three key assumptions: (i) for non-independent studies, that the GWAS results are from the same underlying population, i.e. that the LD pattern is the same across studies, (ii) that there is at most one causal variant in the genomic region for each trait (we assess limitations of this assumption when there are multiple underlying variants in the **Discussion**/**Supplementary Material**), and (iii) that these causal variants are either directly typed or well imputed in all of the GWAS datasets^2,8^.

#### Description of the HyPrColoc method

We define a putative *causal configuration* matrix *S* to be a binary *m* × *Q* matrix, where *m* is the number of traits and *Q* is the number of variants. *S*_*ij*_ is 1 if the *j*^*th*^ variant is causal for the *i*^*th*^ trait and 0 otherwise (**Supplementary Material**). A *hypothesis* uniquely identifies traits which share a causal variant, traits which have distinct causal variants and traits which do not have a causal variant. Except for the null hypothesis (*H*_0_) of no causal variant for any trait, hypotheses such as “ *H*_*m*_ : all *m* traits share a causal variant” correspond to multiple configuration matrices, *S* (**Figure 1**). By considering the set of configurations to which a hypothesis corresponds, the posterior odds of the hypothesis against the null hypothesis can be computed. For example, let 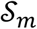 denote the set of configurations for hypothesis *H*_*m*_ and *S*_0_ denote the single configuration for *H*_0_, then the posterior odds for the hypothesis that all traits colocalize to a single causal variant is given by,

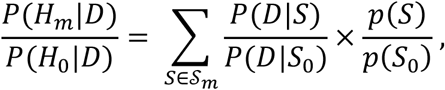

where *D* represents the combined trait data, the first term in the summation is a Bayes factor and the second term is a prior odds^2,8^. To identify a candidate causal variant across the *m* traits, i.e. to perform *multi-trait fine-mapping*, we locate the configuration *S*\* satisfying 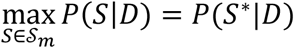. If the summary data for the genetic associations between traits are independent, then the Bayes factor for each configuration *S* can be computed by combining Wakefield’s approximate Bayes factors^13^ for each trait in the configuration (**Methods**). If the summary data between traits are correlated because a subset of the participant data was used in at least two of the GWAS analyses, then an extension to Wakefield’s approximate Bayes factors, which jointly models the trait associations, can be employed (**Methods**). For a given hypothesis *H* and set of corresponding configurations 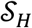, the prior probability of configuration *S*, *p*(*S*), can either be equal for all 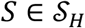, or can be defined as a product of variant-level priors (**Methods**). Our variant-level prior extends that of COLOC^2^ and MOLOC^8^ to a framework that is suitable for the analysis of large numbers of traits. This approach requires specification of only two interpretable parameters: *p*, the probability that a variant is causal for one trait, and *γ*, where 1 − *γ* is the conditional probability that a variant is causal for a second trait given it is causal for one other trait (**Methods**).

**Figure 1:**
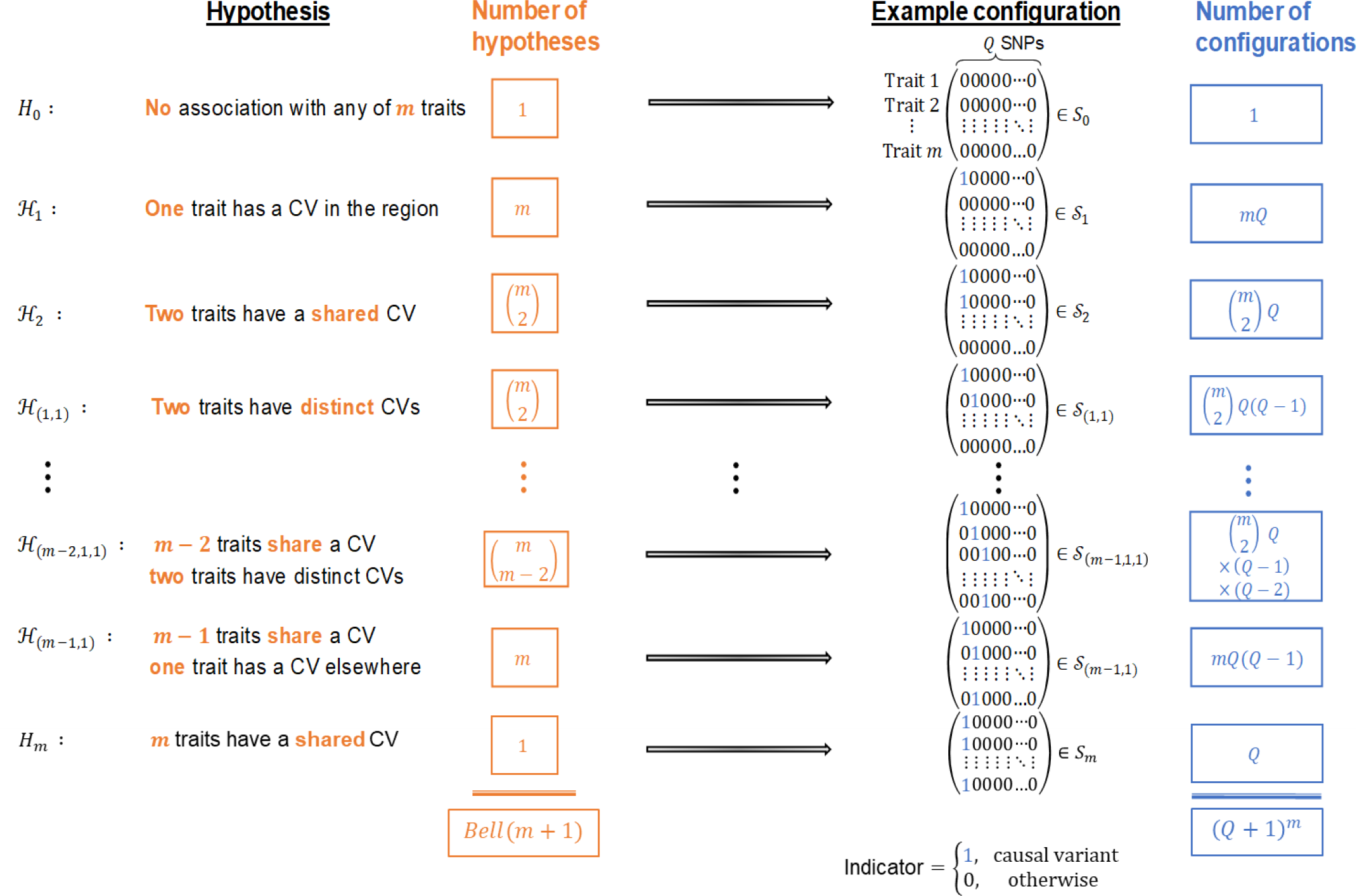
Colocalization hypotheses and causal configurations. Statistical colocalization hypotheses and examples of their associated SNP configurations that allow for at most one causal variant for each of *m* traits in a region containing *Q* genetic variants. For clarity, the hypotheses and a single configuration associated with each hypothesis are shown for *m* ≥ 4 traits, but the column totals *Bell*(*m* + 1) and (*Q* + 1)^*m*^ are correct for *m* ≥ 2.

#### Efficient computation of PPFC

For a pre-specified genomic region comprising *Q* variants, the aim is to evaluate the *PPFC*, *P*(*H*_*m*_|*D*), that all *m* traits share a causal variant within that region, given the summarized data *D*. According to Bayes’ rule, this is given by:

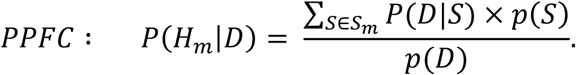

*Brute-force* computation of the denominator, *p*(*D*), requires the exhaustive enumeration of (*Q* + 1)^*m*^ causal configurations, which is computationally prohibitive for *m* > 4, e.g. MOLOC^8^. HyPrColoc overcomes this challenge by approximating *p*(*D*) in a way that is both computationally efficient and tightly bounds the approximation error.

As we show in the **Methods**, the PPFC can be approximated as

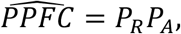

where *P*_*R*_, *P*_*A*_ > 0 are rapidly computable values that quantify the probability that two criteria necessary for colocalization are satisfied (**Figure 2**). The first of these criteria is that all the traits must share an association with one or more variants within the region. *P*_*R*_, which we refer to as the *regional association probability*, is the probability that this criterion is satisfied. By itself, this criterion does not guarantee that there is a single causal variant shared by all traits, because it could be the case that two or more traits have distinct causal variants in strong LD with one another. To safeguard against this, we have a second criterion that ensures the shared associations between all traits are owing to a single shared putative causal variant. *P*_*A*_ is the probability that this second criterion is satisfied. We refer to *P*_*A*_ as the *alignment probability* as it quantifies the probability of alignment at a single causal variant between the shared associations. Both *P*_*R*_ and *P*_*A*_ have *linear* computational cost in the number of traits *m*, making a calculation of 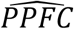 possible when analysing vast numbers of traits. If the first criterion is satisfied, but the second is not, this may be because it is possible to partition the traits into clusters, such that each cluster has a distinct causal variant. HyPrColoc additionally seeks to identify these clusters.

**Figure 2:**
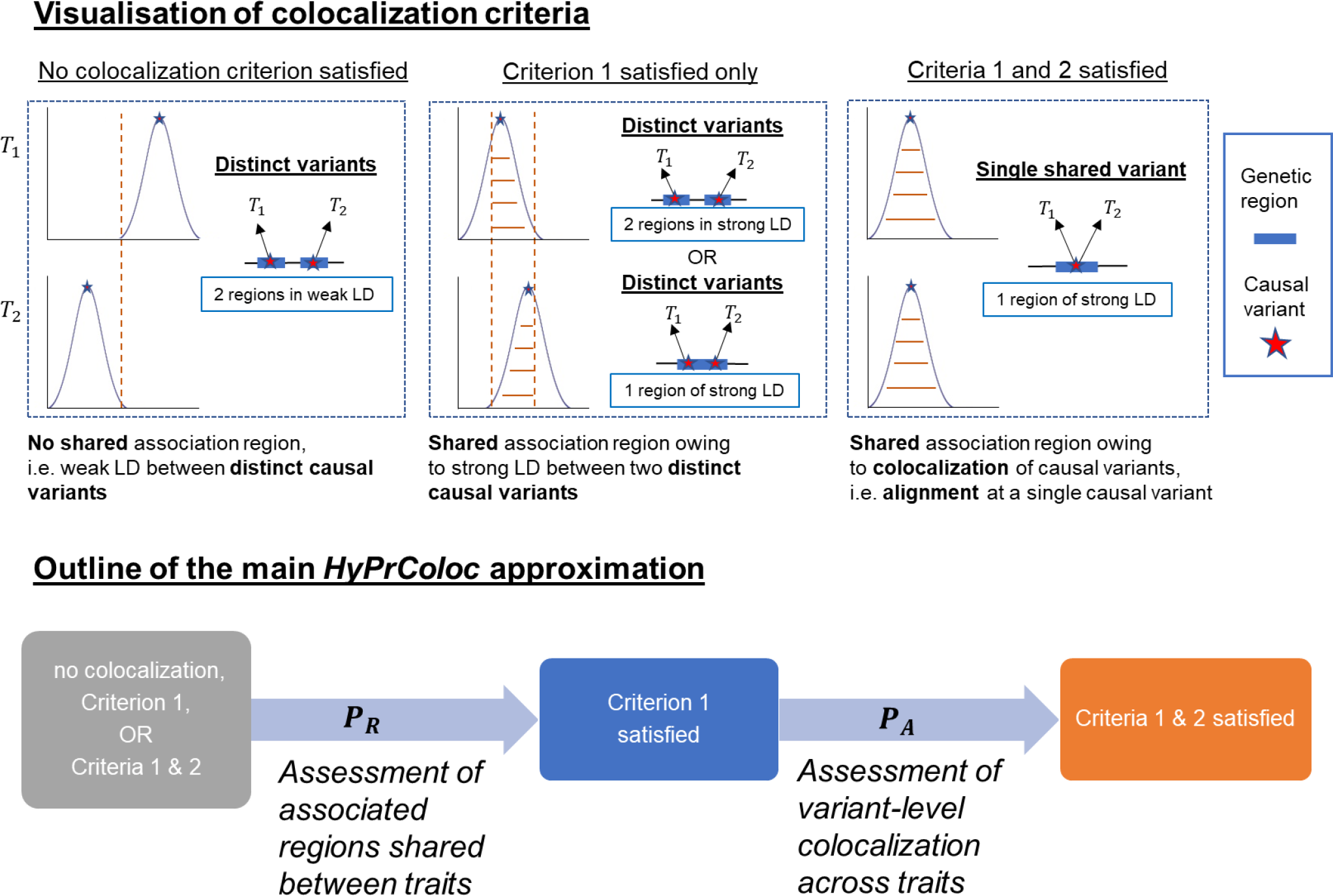
Illustration of the HyPrColoc approximation. We illustrate the HyPrColoc approach with *m* = 2 traits. Statistical colocalization between traits which do not share an association *region*, i.e. do not have shared genetic predictors, is not possible (no colocalization criteria satisfied). However, traits which do (satisfying criterion 1) possess the possibility. HyPrColoc first assesses evidence supporting all *m* traits sharing an association region, which quickly identifies utility in a colocalization mechanism. HyPrColoc then assesses whether any shared association region is due to colocalization between the traits (criteria 1 and 2) or due to a region of strong LD between two distinct causal variants, one for each trait (criterion 1 only). Results from these two calculations are combined to accurately approximate the *PPFC*.

#### Identification of clusters of colocalized traits

If 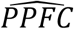 falls below a threshold value, *τ*, we reject the hypothesis *H*_*m*_ that all *m* traits colocalize to a shared causal variant. In practice, this threshold is specified by defining separate thresholds, 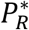 and 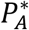, for *P*_*R*_ and *P*_*A*_, such that 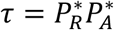 (**Methods**). If *H*_*m*_ is rejected, HyPrColoc seeks to determine if there are values ℓ < *m* such that *H*_ℓ_ cannot be rejected; i.e. if there exist subsets of the traits such that all traits within the same subset colocalize to a shared causal variant. Starting with a single cluster containing all *m* traits, our *branch and bound* divisive clustering algorithm (**Figure 3**) iteratively partitions the traits into larger numbers of clusters, stopping the process of partitioning a cluster of two or more traits when all traits in a cluster satisfy both 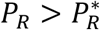 and 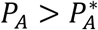. The process of partitioning a cluster into two smaller clusters is performed using one of two criteria: (i) regional (*P*_*R*_) or (ii) alignment (*P*_*A*_) selection (**Methods** and **Supplementary Note**). For *k* ≤ *m* traits in a cluster, the regional selection criterion has 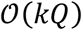 computational cost and is computed from a collection of hypotheses that assume not all traits in a cluster colocalize because one of the traits does not have a causal variant in the region. The alignment selection criterion has 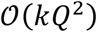 computational cost and is computed from hypotheses that assume not all traits in a cluster colocalize because one of the traits has a causal variant elsewhere in the region (**Supplementary Note**). By default, the HyPrColoc software uses the more computationally efficient regional selection criterion to partition a cluster.

**Figure 3:**
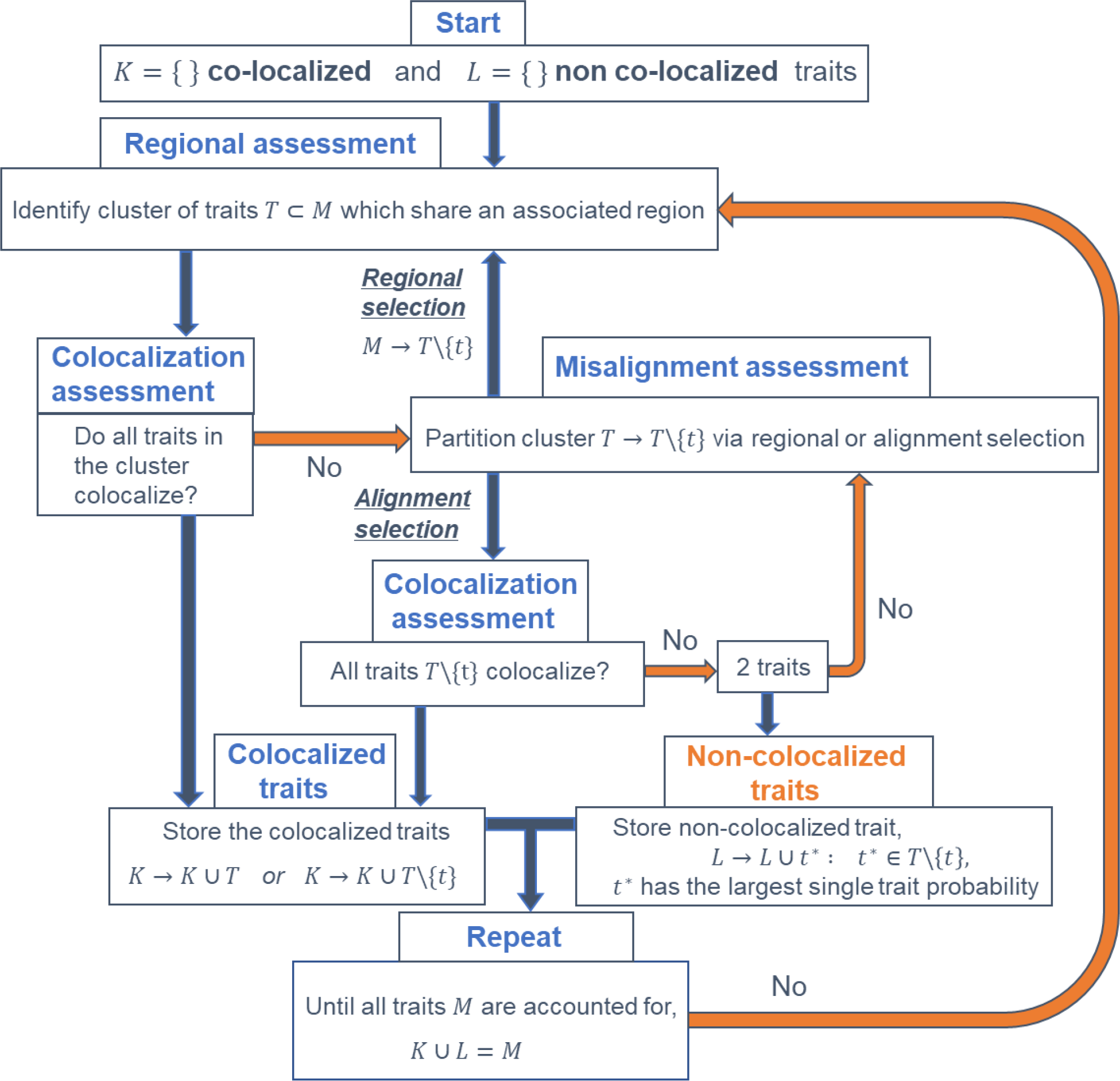
Branch and bound divisive clustering algorithm. Illustration of the pipeline used to detect complex patterns of colocalization. The set of all *m* traits is denoted *M*, *T* denotes a subset (i.e. *cluster*) of traits in *M* and *t* a single trait. The algorithm aims to identify one or more clusters of colocalized traits and stores these clusters in the set *K*. The remaining traits *L*, where *L* = *K*\*M*, are identified as not having or sharing a causal variant with any other trait. The traits in the sample are partitioned into multiple clusters via a *regional* or an *alignment* selection criterion. Regional selection (software default) has 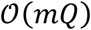 time cost and identifies the trait least likely to share an associated region with the other *m* − 1 traits. Alignment selection identifies the trait whose causal variant is least likely to be shared with the other *m* − 1 traits and has 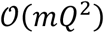 time cost (**Supplementary Note**).

### Model validation using simulations

We created simulated datasets by resampling phased haplotypes from the European samples in 1000 Genomes^14^ and for each dataset we randomly selected one of the first 50 regions confirmed to be associated with CHD^15^ (**Methods**). For each simulation scenario, 1,000 replicates were performed.

#### Computational efficiency

The posterior probability of colocalization, across *m* traits and in a region of *Q* variants, can be accurately approximated by computing 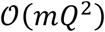 causal configurations. **Figure 4** illustrates this for varying numbers of independent studies and variants, demonstrating a close linear relationship between computation time and the number of traits. Consequently, HyPrColoc is able to assess 100 traits, in a region of 1,000 SNPs, in under 1 second compared to MOLOC which takes approximately one hour to analyse five traits. For *m* ≤ 4, traits the median absolute relative difference between the HyPrColoc and MOLOC^8^ posterior probabilities was found to be ≲ 0.5% (**Figure 4**).

**Figure 4:**
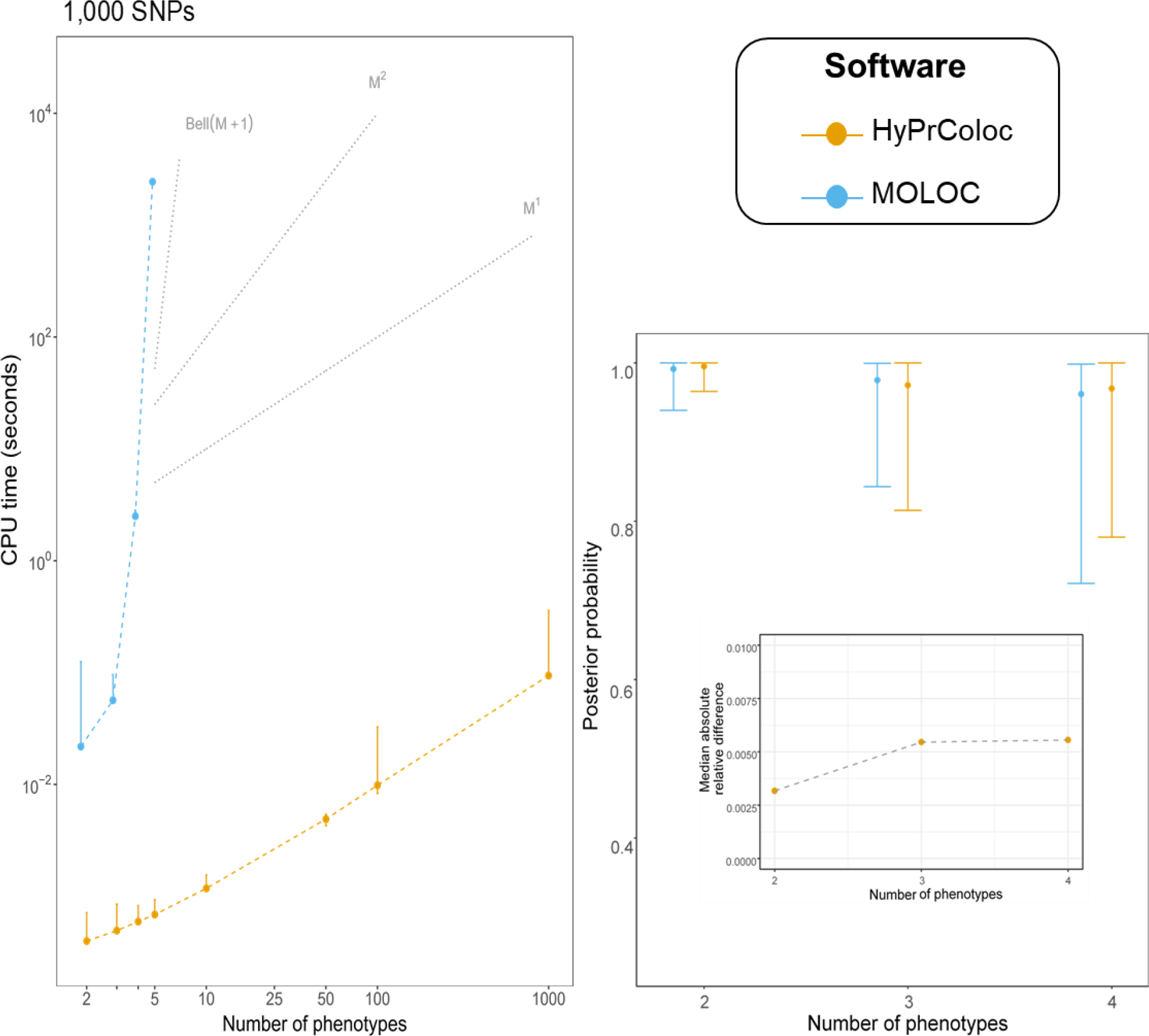
Comparison of HyPrColoc and MOLOC computation time and posterior probability of colocalization. (Left panel) Computation time (seconds) for HyPrColoc (yellow) and MOLOC (blue) to assess full colocalization across *M* ≤ 1000 traits in a region containing *Q* = 1000 SNPs (middle panel). MOLOC was restricted to *M* ≤ 5 traits owing to the computational and memory burden of the MOLOC algorithm when *M* > 5. Three reference lines are plotted: (i) *Bell*(*M* + 1), which denotes the theoretical cost of exhaustively enumerating all hypotheses; (ii) *M*^2^, denoting quadratic cost and; (ii) *M*^1^, denoting the linear complexity of the HyPrColoc algorithm. (Right panel) Distribution of the posterior probability of colocalization using HyPrColoc (yellow) and MOLOC (blue) across *M* ∈ {2,3,4} traits. Where error bars are present, plotted are the 1^st^, 5^th^ (median), and 9^th^ deciles. Despite differences in the prior set-up between the methods, the median absolute relative difference between the two posterior probabilities was ≾ 0.005.

#### Performance of HyPrColoc to detect multi-trait colocalization

We used simulated datasets in which all traits colocalize to assess the accuracy of HyPrColoc in detecting colocalization across varying numbers of traits and study sample sizes. We simulated independent datasets with sample sizes of 5,000, 10,000, and 20,000 individuals for up to 100 quantitative traits and for which all traits share a single causal variant explaining either 0.5%, 1% or 2% of trait variance. For each simulated dataset, we used HyPrColoc to approximate the PPFC. The distribution of PPFC across the simulated datasets was narrower in the analysis of two traits relative to a larger number of traits, as the probability of random misalignment of the lead variant between traits increases as the number of traits increases (top **Figure 5**). However, the estimated PPFC is always close to 1 for 5, 10 and 20 traits illustrating that the distribution of the estimate is stable across a broad number of traits and sample sizes. For 100 traits there is a small decrease in power due to the growth in the number of hypotheses in which only a subset of the traits colocalize. This is expected when sample size is fixed and the shared causal variant explains only a small fraction of trait variation for each trait, as combined evidence supporting hypotheses in which a subset of the traits colocalize are eventually greater than evidence supporting full colocalization.

**Figure 5:**
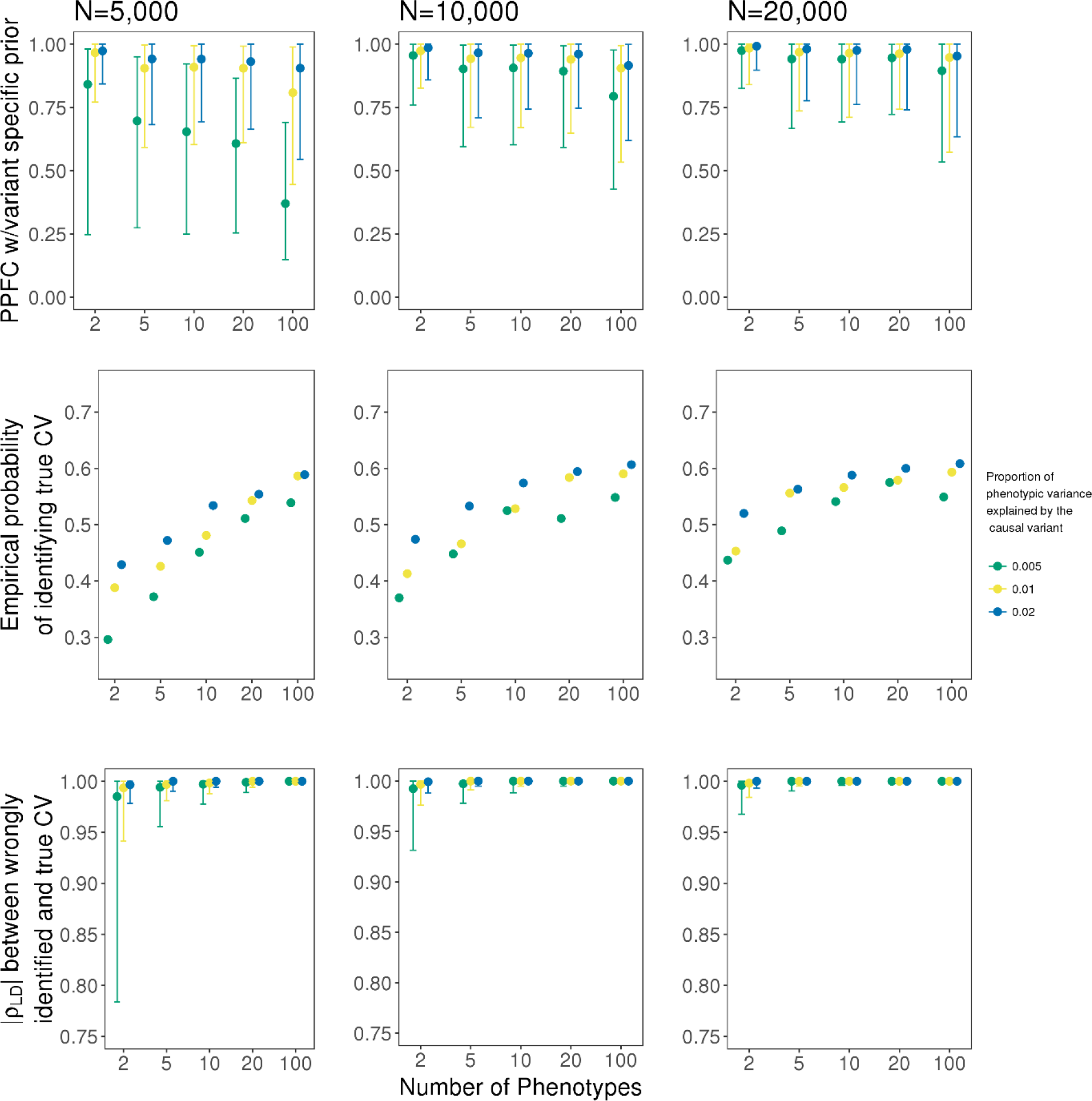
Assessment of the HyPrColoc posterior probability. Simulation results for a sample size *N* ∈ {5000, 10000, 20000} and a causal variant explaining {0.5%, 1%, 2%} of variation across *m* ∈ {2, 5, 10, 20, 100} traits. Presented is the distribution of the HyPrColoc posterior for variant-level priors only (top); the probability of correctly identifying the causal variant (middle) and; linkage disequilibrium between an incorrectly identified causal variant and the true causal variant (bottom). Where error bars are present, plotted are the first, fifth (median), and ninth deciles.

When at least one trait did not have a causal variant in the region the false detection rate was negligible. For example, we generated 100 quantitative traits, each from a study with sample size 10,000, in which 99 traits share a causal variant and the remaining trait had either: (i) a distinct causal variant or (ii) no causal variant in the region. In each scenario a causal variant explained 1% of trait variation. The 1^st^, 5^th^ (median) and 9^th^ deciles of the PPFC were (4 × 10^−24^, 1 × 10^−17^, 5 × 10^−8^) in scenario (i) and (0.02, 0.05, 0.10) in scenario (ii). There is a considerable difference between the results from each scenario, but the PPFC is small in both situations.

#### Fine mapping the causal variant with HyPrColoc

If HyPrColoc identified a variant that was not the true causal variant, we computed the LD between the true causal variant and the identified variant. The proportion which HyPrColoc correctly identified the true causal variant increased as the number of colocalized traits included in the analyses increased up to 2-fold, irrespective of sample size and variance explained by the causal variant (middle **Figure 5**), highlighting a major benefit of performing multi-trait fine-mapping. In cases where the identified variant was not the causal variant, the variant was typically in very strong LD (median *r*^2^ ≥ 0.99) with the true causal variant and for large numbers of traits, i.e. *m* ≥ 20, with sample size 20,000, the two variants were in perfect LD, *i.e. r*^2^ = 1 (bottom **Figure 5**).

#### Branch and bound divisive clustering algorithm

Here we assess the performance of the branch and bound (BB) divisive clustering algorithm to identify clusters of colocalized traits over a range of scenarios. We simulated 100 traits from non-overlapping datasets with 10,000 individuals under three situations: in all scenarios there exists a cluster of 10 traits sharing a single causal variant, 80 traits do not have a causal variant (reflecting “hypothesis free” colocalization searches) and the remaining traits either (i) do not have a causal variant (**Figure 6a**); (ii) form a separate cluster of 10 traits sharing a distinct causal variant (**Figure 6b**) or; (iii) separately have distinct causal variants (**Figure 6c**). In all scenarios, the causal variant for each trait explained 1% of trait variance and the probability parameters were set to 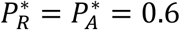(**Methods**). HyPrColoc correctly identified the cluster or clusters of colocalized traits with probability ≈ 0.95 in all simulation scenarios. However, owing to the large number of traits analysed and strong LD between distinct causal variants these clusters occasionally wrongly included one additional trait. To provide insight into when this happens, in each scenario we stratified results into two categories: (a) *P*_*R*_ *P*_*A*_ > 0.6 and (b) *P*_*R*_*P*_*A*_ > 0.7, where *P*_*R*_*P*_*A*_ denotes the posterior probability that a cluster of traits are identified as colocalizing. In scenario (iii) we additionally stratified according to LD between causal variants: (a) *r*^2^ ≤ 1 and (b) *r*^2^ < 0.95. Across all scenarios, the probability of identifying the true cluster(s) of colocalized traits was higher for larger *P*_*R*_*P*_*A*_. For example, in scenarios (i) and (ii) when *P*_*R*_*P*_*A*_ > 0.7 the BB algorithm identifies the true cluster(s) of colocalized traits with probability ≳ 0.9, whereas for *P*_*R*_*P*_*A*_ > 0.6 the true detection probability was lower but still > 0.8. When many traits have a distinct causal variant, scenario (iii), the probability of detecting the true cluster of colocalized traits dropped markedly (≈ 0.7). This was due to the increased chance that the causal variant from a non-colocalized trait is in strong LD with the colocalized causal variant, i.e. *r*^2^ ≥ 0.95, a scenario in which no algorithm is likely to perform well. In scenarios where *r*^2^ < 0.95, for all causal variants, the true detection probability was ≳ 0.9. We found an increase in the true detection probabilities of the BB algorithm when analysing 20 traits under a similar simulation framework (**Supplementary Material, Figure S2**), indicating that the performance of the algorithm is somewhat dependent upon the number of traits under consideration. Overall, across the range of scenarios considered the selection algorithm performs well in terms of sensitivity and specificity.

**Figure 6:**
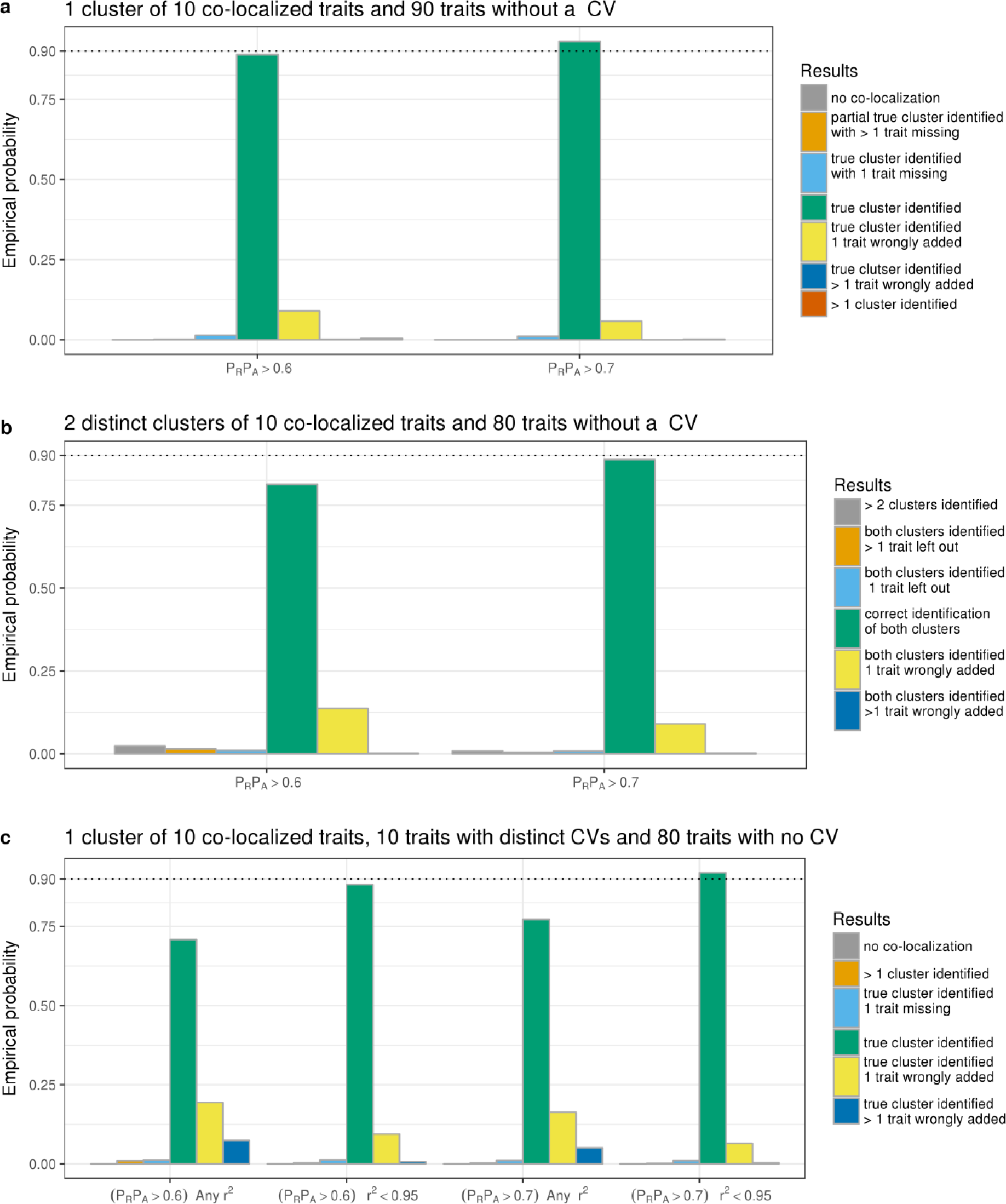
Assessing the performance of the BB clustering algorithm. In each of the three scenarios presented, *m* = 100 traits with non-overlapping samples were generated, all traits had a study sample size of *N* = 10000 and variant-level causal configuration priors were used. In all scenarios there exists at least one cluster of 10 traits which share a causal variant, 80 traits which do not have a causal variant and either: (a) the remaining traits do not have a causal variant in the region; (b) there exists another cluster of 10 traits which share a distinct causal variant or; (c) all remaining traits have a causal variant and these variants are ‘distinct’ from one another (a distinct variant can be in perfect LD, i.e. *r*^2^ = 1, with another distinct variant and/or the shared causal variant). In all scenarios the detection probability is presented by posterior probability of colocalization, i.e. *P*_*R*_*P*_*A*_ ≥ (0.6, 0.7). Where indicated, detection probabilities are presented by LD (*r*^2^) between the causal variant, shared across the 10 (default) colocalized traits, and any other distinct causal variant, i.e. when *r*^2^ ≤ (1, 0.95).

We further tested the algorithm using a variety of thresholds 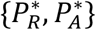, two different prior frameworks and accounting for overlapping samples in analyses (**Figures S4-5**). We demonstrated that treating studies as independent, even when there is complete sample overlap (i.e. participants are the same in all studies) gives reasonable results (**Figure S3**). We discuss the theoretical reasons for this in **Supplementary Material**. We also assessed the reliability of the BB algorithm when a secondary causal variant was added to one or more traits in the region. Our results indicate continued good performance when a secondary causal variant explains less trait variation than the shared causal variant (**Supplementary Material and Table S5**).

### Map of genetic risk shared across CHD and related traits

We used HyPrColoc to investigate genetic associations shared across CHD^16^ and 14 related traits: 12 CHD risk factors^17–21^, a comorbidity^22^ and a social factor^23^ (Supplementary Table S1 for details). We performed colocalization analyses in pre-defined disjoint LD blocks spanning the entire genome^24^. To highlight that multi-trait colocalization analyses can aid discovery of new disease-associated loci, we used the CARDIoGRAMplusC4D 2015 data for CHD^16^, which brought the total number of CHD associated regions to 58, and contrasted our findings with the current total of ~160 CHD associated regions^25^. For each region in which CHD and at least one related trait colocalized, we integrated whole blood gene expression^26^ quantitative trait loci (eQTL) and protein expression^27^ quantitative trait loci (pQTL) information into our analyses to prioritise candidate causal genes (**Methods**).

#### Multi-trait colocalization

Our genome-wide analysis identified 43 regions in which CHD colocalized with one or more related traits (**Figure 7** and **Table 1**). Twenty-three of the 43 colocalizations involved blood pressure, consistent with blood pressure being an important risk factor for CHD^28^. Other traits colocalizing with CHD across multiple genomic regions were cholesterol measures (16 regions); adiposity measures (9 regions); type 2 diabetes (T2D; 4 regions) and; rheumatoid arthritis (2 regions). Moreover, by colocalizing CHD and related traits, our analyses suggest these traits share some biological pathways.

**Table 1.** Forty-three regions with colocalized associations across CHD and 14 related traits. Loci are sorted into three categories: (i) those *known* at the time of the release of CARDIoGRAMplusC4D 2015 data for CHD^16^; (ii) those *later identified* in a subsequent study (or studies) or; (iii) those that have not been previously reported and are considered *future candidate* CHD loci.

| Known CHD loci identified by HyPrColoc that share associations with CHD related traits |  |  |  |  |  |  |  |  |
| --- | --- | --- | --- | --- | --- | --- | --- | --- |
| Chr | Locus | Traits | Colocalized SNP (consequence) | Gene | Known CHD locus (known CHD SNP) | PPFC (PPE) | Expressed gene (eQTL) | Protein (pQTL) |
| 2 | *ABCG8, ABCG5* | CHD, LDL | rs4299376 (Intron) | *ABCG8* | Yes<sup>31</sup> (Yes<sup>31</sup>) | 0.9176 (0.9486) | - | - |
| 4 | *GUCY1A1* | CHD, DBP | rs72689147 (Intron) | *GUCY1A1* | Yes<sup>15</sup> (Yes<sup>16</sup>) | 0.931 (0.2409) | *GUCY1A1* (rs12643599) | - |
| 6 | *PHACTR1, EDN1* | CHD, SBP | rs9349379 (Intron) | *PHACTR1* | Yes<sup>32,34</sup> (Yes<sup>32</sup>) | 0.9994 (1) | - | - |
| 6 | *LPA* | CHD, LDL | rs10455872 (Intron) | *LPA* | Yes<sup>31,34</sup> (Yes<sup>31,34</sup>) | 0.998 (0.5383) | - | - |
| 7 | *HDAC9* | CHD, SBP | rs2107595 (Intergenic) | *HDAC9* | Yes<sup>15</sup> (Yes<sup>16</sup>) | 0.9961 (0.7294) | - | - |
| 7 | *ZC3HC1, KLHDC10* | CHD, DBP | rs11556924 (Missense) | *ZC3HC1* | Yes<sup>15,31,34</sup> (Yes<sup>15,31,34</sup>) | 0.9998 (0.9936) | - | - |
| 8 | *TRIB1* | CHD, HDL, LDL, TG, eGFR | rs2954029 (Intron) | *RP11-136O12.2* | Yes<sup>15</sup> (Yes<sup>15</sup>) | 0.925 (0.8724) | - | - |
| 9 | *ANRIL, CDKN2B-AS1* | CHD, DBP | rs2891168 (Intron) | *CDKN2B-AS1* | Yes<sup>16</sup> (Yes<sup>16</sup>) | 0.8696 (0.7552) | - | - |
| 9 | *ABO* | CHD, LDL, DBP, T2D | rs507666 (Intron) | *ABO* | Yes<sup>15,34</sup> (Yes<sup>16</sup>) | 0.9835 (0.5825) | - | - |
| 10 | *KIAA1462* | CHD, DBP | rs1887318<br>(Intron) | *KIAA1462* | Yes<sup>15,32</sup><br>(Yes<sup>16</sup>) | 0.9369 (0.4331) | - | - |
| 11 | *APOA1-C3-A4-A5* | CHD,<br>HDL,<br>LDL, TG | rs964184<br>(3 prime UTR) | *ZPR1*,<br>*BUD13* | Yes<sup>34</sup> (Yes<sup>34</sup>) | 0.9572 (1) | - | Apolipoprotein<br>A-V<br>(rs964184) |
| 12 | *ATP2B1* | CHD, SBP | rs2681492<br>(Intron) | *ATP2B1* | Yes<sup>16</sup> (Yes<sup>16</sup>) | 0.9803 (0.3027) | - | - |
| 12 | *SH2B3* | CHD,<br>HDL,<br>LDL, SBP,<br>DBP, RA | rs7137828<br>(Intron) | *ATXN2* | Yes<sup>34</sup> (Yes<sup>16</sup>) | 0.9094 (0.7684) | *TRAFD1*<br>(rs7137828) | - |
| 15 | *FES, FURIN* | CHD, SBP,<br>DBP | rs35346340<br>(Splice region) | *FES* | Yes<sup>15</sup> (Yes<sup>16</sup>) | 0.9597 (0.5789) | *FES*<br>(rs8027450) | - |
| 18 | *MC4R, PMAIP1* | CHD,<br>HDL, TG,<br>BMI, WC | rs12967135<br>(Intergenic) | - | Yes<sup>16</sup>(Yes<sup>16</sup>) | 0.8585 (0.4337) | - | - |
| 19 | *LDLR, SMARCA4* | CHD, LDL | rs112374545<br>(Intergenic) | *LDLR* | Yes<sup>15,34</sup><br>(Yes<sup>16</sup>) | 0.9374 (0.5563) | - | - |
| 19 | *APOC1, APOE,*<br>*PVRL2, COTL1* | CHD,<br>HDL, WC | rs4420638<br>(Downstream) | *APOC1* | Yes<sup>16</sup> (Yes<sup>16</sup>) | 0.9596 (0.9997) | - | Apolipoprotein E<br>(rs4420638) |
| 21 | *KCNE2* | CHD, DBP | rs28451064<br>(Intron) | *AP000318.2* | Yes<sup>16</sup> (Yes<sup>16</sup>) | 0.9982 (0.9735) | - | - |
| **CHD loci reported after time of data release (2015) identified by HyPrColoc to share associations with CHD related traits** |  |  |  |  |  |  |  |  |
| 1 | *PRDM16* | CHD, SBP,<br>DBP | rs2493288<br>(Intron) | *PRDM16* | Yes<sup>25</sup> (Yes<sup>25</sup>) | 0.8009 (0.3471) | - | - |
| 1 | *FHL3* | CHD, SBP | rs34655914<br>(Missense) | *INPP5B* | Yes<sup>25</sup> (Yes<sup>25</sup>) | 0.9468 (0.0832) | *SF3A3*<br>(rs28428561);<br>*UTP11L*<br>(rs4360494);<br>*RNU6-510P*<br>(rs61776719) | - |
| 1 | *SORT1* | CHD,<br>HDL | rs12740374<br>(3 prime UTR) | *CELSR2* | Yes<sup>25</sup> (Yes<sup>25</sup>) | 0.9898 (0.9997) | - | - |
| 1 | *LMOD1* | CHD,<br>BMI, WC | rs2678204<br>(Intron) | *IPO9* | Yes<sup>29</sup> (Yes<sup>29</sup>) | 0.8273 (0.1627) | *IPO9*<br>(rs2494115) | - |
| 2 | *FIGN* | CHD, SBP | rs268263<br>(Intron) | *AC092684.1* | Yes<sup>25</sup> (Yes<sup>25</sup>) | 0.789 (0.995) | - | - |
| 2 | *IRS1* | CHD, HDL, TG | rs62188784<br>(Intergenic) | *AC068138.1* | Yes<sup>25</sup> (Yes<sup>25</sup>) | 0.8234 (0.4852) | - | - |
| 3 | *RHOA* | CHD, BMI, EDU | rs73078367<br>(Downstream) | *NCKIPSD* | Yes<sup>25</sup> (Yes<sup>25</sup>) | 0.9541 (0.5656) | - | - |
| 3 | *RHOA* | CHD, SBP | rs7623687<br>(Intron) | *RHOA* | Yes<sup>33</sup> (Yes<sup>33</sup>) | 0.9713 (0.2455) | - | - |
| 4 | *FGF5, PRDM8* | CHD, SBP, DBP | rs13125101<br>(Intergenic) | *FGF5* | Yes<sup>25</sup> (Yes<sup>25</sup>) | 0.9827 (0.4148) | - | - |
| 5 | *MAP3K1* | CHD, HDL, TG, WC, SBP, T2D | rs9686661<br>(Intron) | *C5orf67* | Yes<sup>25</sup> (Yes<sup>25</sup>) | 0.7755 (0.7115) | - | - |
| 6 | *VEGFA* | CHD, HDL, TG, BMI, WC | rs998584<br>(Downstream) | *VEGFA* | Yes<sup>25</sup> (Yes<sup>25</sup>) | 0.8376 (0.9746) | - | - |
| 10 | *TSPAN14, FAM213A* | CHD, RA | rs2343306<br>(Intron) | *TSPAN14* | Yes<sup>25</sup> (No) | 0.9064 (0.7279) | - | - |
| 11 | *ARNTL* | CHD, DBP | rs10832013<br>(Upstream) | *ARNTL* | Yes<sup>25</sup> (Yes<sup>25</sup>) | 0.9403 (0.0823) | - | - |
| 11 | *SIPA1* | CHD, HDL, TG | rs12801636<br>(Intron) | *PCNX3* | Yes<sup>29</sup> (Yes<sup>29</sup>) | 0.8369 (0.8945) | - | - |
| 12 | *HNF1A* | CHD, LDL | rs1169288<br>(Missense) | *HNF1A* | Yes<sup>29</sup> (Yes<sup>29</sup>) | 0.9645 (0.5762) | - | - |
| 13 | *N4BP2L2, PDS5B* | CHD, BMI | rs35193668<br>(Intron) | *N4BP2L2* | Yes<sup>25</sup> (Yes<sup>25</sup>) | 0.6785 (0.0911) | *N4BP2L2*<br>(rs9337) | - |
| 16 | *CDH13* | CHD, DBP | rs7500448<br>(Intron) | *CDH13* | Yes<sup>25</sup> (Yes<sup>25</sup>) | 0.9947 (1) | - | - |
| 16 | *CTRB2, BCAR1* | CHD, T2D | rs55993634<br>(Downstream) | *CTRB2* | Yes<sup>33</sup> (Yes<sup>25</sup>) | 0.8296 (0.3868) | *BCAR1*<br>(rs28595463) | - |
| 17 | *IGF2BP1* | CHD, BMI, T2D | rs11079849<br>(Intron) | *IGF2BP1* | Yes<sup>25</sup> (Yes<sup>25</sup>) | 0.8389 (0.831) | - | - |
| 17 | *PECAM1, DDX5, TEX2* | CHD, SBP, DBP | rs1867624<br>(Upstream) | *RPL31P57* | Yes<sup>29</sup> (Yes<sup>29</sup>) | 0.7963 (0.4276) | - | - |
| New CHD loci shown to share associations with CHD related traits using HyPrColoc and yet to be reported |  |  |  |  |  |  |  |  |
| **6** | *FHL5* | CHD, SBP | rs9486719<br>(Intron) | *FHL5* | - | 0.844 (0.1542) | - | - |
| **10** | *CYP26A1* | CHD, TG | rs2068888<br>(Downstream) | *CYP26A1* | - | 0.8454 (0.7669) | - | - |
| **16** | *ANKRD11* | CHD, WC | rs11643561<br>(Intron) | *ANKRD11* | - | 0.7827 (0.0795) | - | - |
| **19** | *RSPH6A* | CHD, SBP | rs8108474<br>(Intron) | *RSPH6A* | - | 0.7802 (0.1435) | - | - |
| **20** | *PREX1* | CHD, SBP,<br>DBP | rs79044887<br>(Intron) | *PREX1* | - | 0.7237 (0.132) | - | - |
| Colocalization analyses were performed genome-wide using publicly available data (Table S1). Chr: chromosome; Locus: labelled with candidate causal genes as listed by Erdmann et al. <sup>39</sup>; Gene: nearest gene to colocalized SNP; eQTL: gene expression<sup>26</sup>; pQTL: protein expression<sup>27</sup>; Colocalized SNP(consequence); SNP marking the association shared across the traits and its annotation in VEP<sup>48</sup> obtained from PhenoScanner<sup>49</sup>; Locus at time of 2015 CHD data release<sup>16</sup>; region was either known and published in<sup>16</sup> or later identified<sup>25</sup>; PPFC: posterior probability of colocalization; PPE: proportion of PPFC explained by the listed SNP; traits: the traits with the colocalized SNP association. The full results from these analyses are available at https://jrs95.shinyapps.io/hyprcoloc_chd. |  |  |  |  |  |  |  |  |

**Figure 7:**
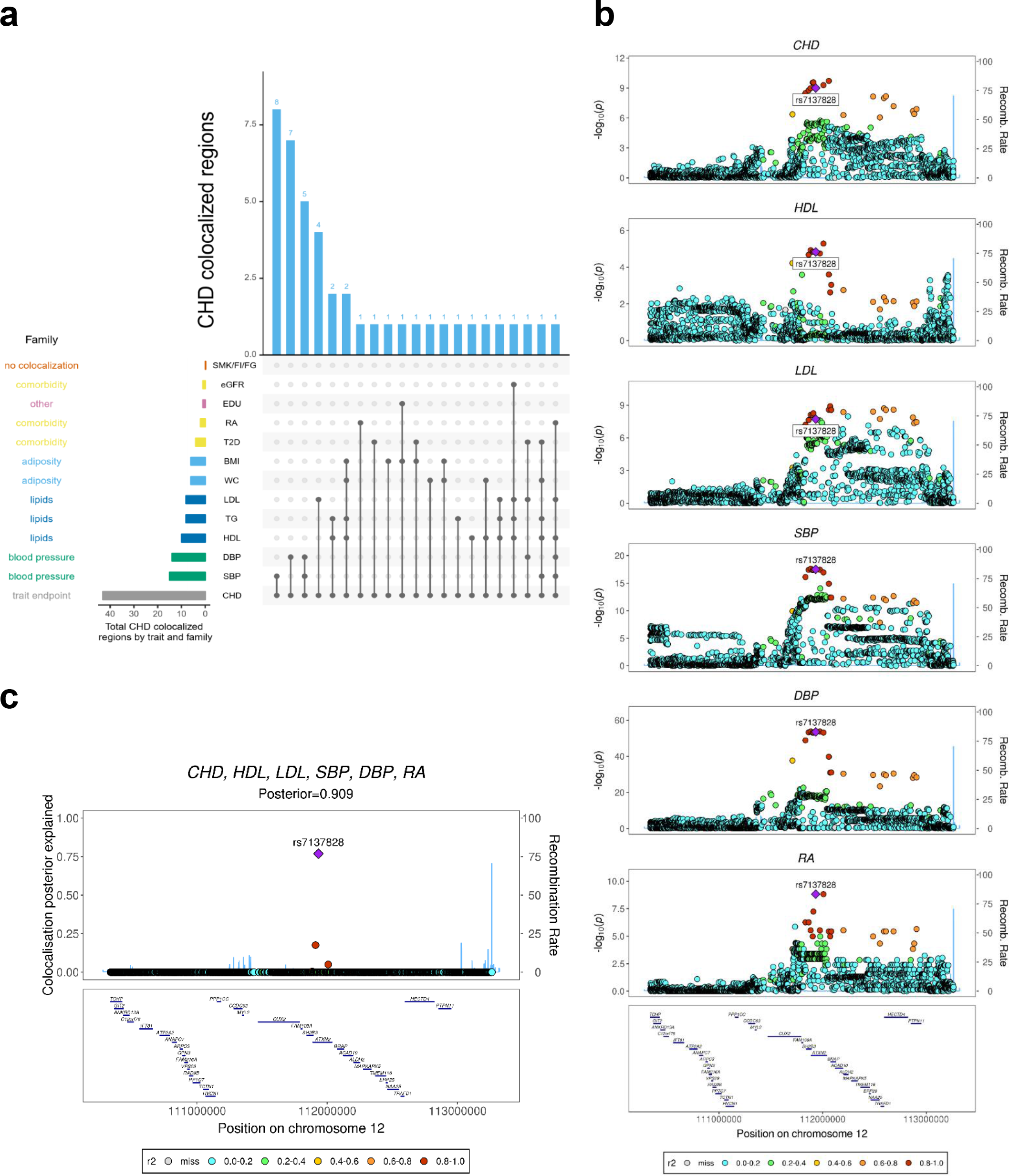
Genome-wide multi-trait colocalization analysis of CHD and fourteen related traits. (a) Summary of the number of regions across the genome in which CHD colocalizes with at least one related trait. Results are aggregated by trait family, e.g. lipid fractions, and by each individual trait. (b) Stacked association plots of CHD with high density lipoprotein (HDL), low density lipoprotein (LDL), systolic blood pressure (SBP), diastolic blood pressure (DBP) and rheumatoid arthritis (RA). HyPrColoc implicated both the *SH2B3-ATXN2* locus and risk variant rs713782, both of which have been previously reported as associated with CHD risk^25^. However, HyPrColoc extended this result by identifying that the risk loci and variant are shared with 5 conventional CHD risk factors^11^. (c) HyPrColoc identified rs713782 as a candidate causal variant explaining the shared association signal between CHD and the 5 related traits, i.e. rs713782 explained over 76% of the posterior probability of colocalization whereas the next candidate variant explained < 20%.

In thirty-eight of the 43 (88%) colocalized regions^15,16,25,29–34^, the candidate causal SNP proposed by HyPrColoc and/or its nearest gene, have been previously identified. Importantly, 20 of these were reported *after* the CARDIoGRAMplusC4D study^16^. For example, *FGF5* was sub-genome-wide significant (*P*>5×10^−8^) with CHD in the 2015 data, but through colocalization with blood pressure, we highlight it as a CHD locus and it is genome-wide significant in the most recent CHD GWAS^25^. The remaining 18 regions were reported previously, but one, *APOA1-C3-A4-A5*, was sub-genome-wide significant in the CARDIoGRAMplusC4D study^16^ despite having been reported previously^34^. However, we used HyPrColoc to show that the association of major lipids colocalize with a CHD signal, highlighting this as a CHD locus in these data (**Table 1** and **Figure** S**6**). The locus has subsequently been replicated^25,30^ and we show below that the signal also colocalizes with circulating apolipoprotein A-V protein levels (**Table 1**). This demonstrates that joint colocalization analyses of diseases and related traits can improve power to detect new associations (an approach which is advocated outside of colocalization studies^35^). Our results also illustrate that multi-trait colocalization analyses can provide further insights into well-known risk-loci of complex disease. For example, at the well-studied *SH2B3*-*ATXN2* region^25,34^, HyPrColoc detected two cholesterol measures (LDL, HDL), two blood pressure measures (SBP, DBP) and rheumatoid arthritis (RA) colocalizing with CHD at the previously reported CHD associated SNP^25^ rs7137828 (PPFC=0.909 of which 76.8% is explained by the variant rs7137828; **Figure 7**). In addition, we newly implicated a candidate SNP and locus in a further 5 CHD regions not previously associated with CHD risk (**Table 1**). In one of the 5 regions, *CYP26A1*, CHD colocalized with tri-glycerides (TG) and HyPrColoc identified a single variant that explained over 75% of the posterior probability of colocalization, supporting this SNP as a candidate shared CHD/TG variant.

For each of the 43 regions that shared genetic associations across CHD and related traits, we further integrated whole blood gene^26^ and protein^27^ expression into the colocalization analyses. We tested *cis* eQTL for 1,828 genes and *cis* pQTL from the 854 published proteins across the 43 loci for colocalization with CHD and the related traits. Of the 43 listed variants (**Table 1**), 27 were associated with expression of at least one gene (*P*<5×10^−8^) and a total of 125 such genes were identified. HyPrColoc refined this, identifying six regions colocalizing with eQTL for one expressed gene and one region, the *FHL3* locus, colocalizing with expression of three genes (*SF3A3*, *UTP11L*, *RNU6-510P*) (**Table 1**). The *GUCY1A3* locus has previously been associated with BP^36^ and with CHD^15^. Here we show that these associations are likely to be due to the same variant, rs72689147 (PPFC=0.93), with the G allele increasing DBP and risk of CHD. We furthermore show that the association colocalizes with expression of *GUCY1A1* in whole blood, with the G allele reducing *GUCY1A1* expression (PPFC=0.77; **Table 1**). The *GUCY1A1* gene is ubiquitously expressed in heart tissues, including in the coronary and aortic arteries^37^. In the mouse, higher expression of *GUCY1A1* has been correlated with less atherosclerosis in the aorta^38^. *GUCY1A1* is a likely candidate gene in this locus^39^, illustrating the utility of HyPrColoc to help prioritise candidate causal genes. The *CTRB2*-*BCAR1* locus was not known at the time of the release of the 2015 CARDIoGRAMplusC4D data, however we find the association at this locus is shared with T2D (PPFC=0.83) and that *BCAR1* expression colocalized with the CHD association (PPFC=0.86). Other studies have implicated the locus in CHD33 and suggested *BCAR1* as the causal gene in carotid intimal thickening^40,41^. We note that two CHD loci also colocalize with circulating plasma proteins, *APOA1-C3-A4-A5*, with apolipoprotein A-V and the *APOE* locus with apolipoprotein E (Table 1).

Of the 38 known CHD loci that colocalized with a related trait, 8 are reported to have a single causal variant^25^, of these we identified the same CHD-associated variant (or one in LD with either r^2^>0.8 or |D’|>0.8)^14^ at seven loci (*SORT1*, *PHACTR1*, *ZC3HC1*, *CDKN2B-AS1*, *KCNE2*, *CDH13, APOE*). Despite the possible presence of multiple causal associations at other loci, HyPrColoc was still able to pick out single shared associations across traits: a result supported by our simulation study when additional distinct causal variants explain less trait variation than that explained by a shared causal variant between colocalized traits (**Supplementary Material**).

## Discussion

We have developed and applied a deterministic Bayesian colocalization algorithm, HyPrColoc, for multi-trait statistical colocalization analyses. HyPrColoc is based on the same underlying statistical model as COLOC^2^, but for the first time enables colocalization analyses to be performed across massive numbers of traits, owing to the novel insight that the posterior probability of colocalization at a single causal variant can be accurately approximated by enumerating only a small number of putative causal configurations. The HyPrColoc algorithm was validated using simulations and used to assess genetic risk shared across CHD and related traits. Using CHD data from 2015^16^, in which 46 regions were genome-wide significant (*P*<5×10^−8^), our multi-trait colocalization analysis identified 43 regions in which CHD colocalized with ≥1 related trait. With this approach, we were able to identify CHD loci that were not known at the time of the data release (2015), demonstrating the benefit of synthesising data on related traits to uncover potential new disease-associated loci^8,35^. A further five regions, we postulate, may be identified as CHD loci in the future. Others have considered pleiotropic effects of CHD loci previously^42^, but our formal colocalization analyses are more robust, *e.g*. in the *ABO* region we show colocalization of T2D and DBP in addition to the previously reported pleiotropic effect with LDL. We integrated eQTL and pQTL data to prioritise candidate genes at some loci, e.g. *GUCY1A1*, *BCAR1* and *APOE*.

The HyPrColoc algorithm identifies regions of the genome where there is evidence of a shared causal variant (by dissecting the genome into distinct regions) and also allows for a targeted analysis of a specific genomic locus of primary interest, e.g. when aiming to identify the perturbation of a biological pathway through the influence of a particular gene. Moreover, these region-specific analyses can highlight candidate causal genes, which will help improve biological understanding and may indicate potential drug targets to inform medicines development^43^.

We have described HyPrColoc under the assumption of at most one causal variant per trait. Future work is required to extend this methodology and algorithm to multiple-causal variants. However, we note that the reliability of results under the single causal variant assumption only break down when secondary causal variants explain as much trait variation as the shared variant (**Supplementary Material**). An example of which is the expression of *SH2B3*, where multiple causal variants for the expression of this gene masks colocalization with the CHD signal. We note that misspecification of LD between causal variants has a major impact on correct detection of multiple causal variants in a region^44^, making a single causal variant assessment the most reliable when accurate study-level LD information is not available. To overcome challenges when specifying the prior probability of a causal configuration, we have suggested two different parsimonious configuration priors that allow a sensitivity analysis to the type of prior and the choice of hyper-parameters to be performed (**Methods**). Nevertheless, other priors may be more appropriate for particular applications.

In summary, we have developed a computationally efficient method that can perform multi-trait colocalization on a large scale. As the size and scale of available data on genetic associations with traits increase, computationally scalable methods such as HyPrColoc will be increasingly valuable in prioritizing causal genes and revealing causal pathways.

## Software availability

We developed an R package for performing the HyPrColoc analyses (https://github.com/jrs95/hyprcoloc). The regional association plots (as seen in Figure 7) were created using gassocplot (https://github.com/jrs95/gassocplot) and LD information from 1000 Genomes^14^.

## Acknowledgements

The authors would like to thank Prof Frank Dudbridge, University of Leicester, who provided helpful comments on the manuscripts and Dr Robin Young, Robertson Centre for Biostatistics, University of Glasgow, for help with the simulation study. This work was funded by the UK Medical Research Council (MR/L003120/1, MC UU 00002/7), British Heart Foundation (RG/13/13/30194), and the UK National Institute for Health Research Cambridge Biomedical Research Centre. The LD information was computed using the phased haplotypes from the 1000 Genomes study (http://www.internationalgenome.org/). The data on coronary artery disease, glycaemic traits, lipid measures, smoking, education, renal function and arthritis have been contributed by CARDIoGRAMplusC4D (www.cardgogramplusc4d.org), MAGIC (www.magicinvestigators.org), GLGC (www.lipidgenetics.org), TAG (https://www.med.unc.edu/pgc/results-and-downloads), SSAGC (www.thessgac.org), DIAGRAM (www.diagram-investigators.org) and CKDGen (http://ckdgen.imbi.uni-freiburg.de) and Okada *et al*. (plaza.umin.ac.jp/~yokada/datasource/software.htm) investigators, respectively. The data on adiposity measures and blood pressure are from the first release of the Neale Lab’s GWAS analysis of UK-Biobank (http://www.nealelab.is/uk-biobank). The data on gene expression and protein expression in whole blood have been contributed by eQTLGen (http://www.eqtlgen.org/cis-eqtls.html) and Sun *et al*. (https://www.phpc.cam.ac.uk/ceu/proteins/), respectively.

## Author contributions

C.N.F. developed the mathematical and statistical methodology, developed the statistical software and applied the methods to the analysis of CHD and related risk factors. J.R.S advised on the statistical methodology and software, developed the bioinformatical software and command-line tool, designed and applied the methods to the analysis of CHD and related risk factors. P.G.B. contributed to the statistical methodology. B.B.S. designed the analysis of CHD and related risk-factors. P.D.W.K. and S.B. revised and reviewed the statistical methodology and scientific content. J.M.M.H contributed to the overall scientific content and goals of the project. All authors contributed to the writing of the manuscript.

## Methods

### SNP association models

Let *Y*_*i*_ denote one of *i* = 1,2, …, *m*, traits assessed in a maximum of *m* studies, i.e. two or more traits can be measured in the same study, and *G*_*ij*_ denote the genotype of the *j*^th^ genetic variant.

It is assumed that the outcome model for *Y*_*i*_ is given by

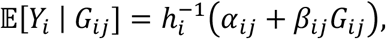

where *α*_*ij*_ is the intercept term and *h*_*i*_ is a function linking the *i*^*th*^ outcome to the genotype *G*_*ij*_, for all *j* = 1,2, …, *Q* genetic variants in the genomic region. The function *h*_*i*_ is typically taken as the identity function for continuous traits and the logit function for binary traits. The aim of colocalization analyses is to identify genomic loci where there exists an *G*_*ij*_ that is causally associated with at least two of the *m* traits. For each of the *m* traits and *Q* genetic variants, we assume that GWAS summary statistics 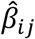 and 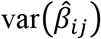 are available. We use these data to perform colocalization analyses in genomic loci.

### Colocalization posterior probability

Using binary vectors to indicate whether a variant putatively causally influences a trait, we can define causal configurations (*S*) that can be grouped into sets 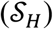 which belong to a single data generating hypothesis (*H*). We use the notation 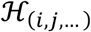 to denote a *set* of hypotheses in which a collection of *i* traits share a causal variant, a separate collection of *j* traits share a distinct causal variant, and so on (Figure 1). For, example, 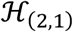 denotes the set of hypotheses in which each hypothesis specifies uniquely 2 traits that share a causal variant, a single trait has a distinct causal variant and all remaining *m* − 3 traits do not have a causal variant in the region. Assuming at most one causal variant for each trait these data generating hypotheses can be combined to generate a hypothesis space (Ω). The posterior probability of hypothesis *H*, given the combined data *D* from all *m* studies, can therefore be computed using (**Supplementary Material**),

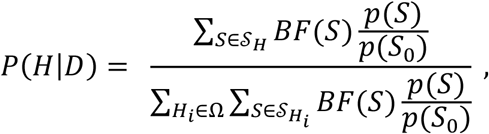

where *p*(*S*)/*p*(*S*_0_) is the prior-odds of configuration 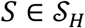 compared with the null-configuration *S*_0_, *i.e*. no genetic association with any trait. See^2^ for a derivation with *m* = 2 traits. *BF*(*S*) is a Bayes factor which is the likelihood of the data being generated under 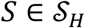 relative to the likelihood of the data being generated *S*_0_.

### Computing Bayes Factors: independent studies

If the trait associations are calculated using independent studies (i.e. no overlapping samples in the GWAS datasets), the Bayes factors can be computed using Wakefield’s Approximate Bayes Factors^13^ (*ABF*) for each trait *i* and genetic variant *j*, i.e.

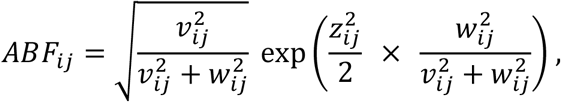

where *z*_*ij*_, *ν*_*ij*_ and *w*_*ij*_ are the Z-statistic, standard error and the prior standard deviation for 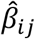, respectively. Following^2^, for continuous variables *w*_*ij*_ is set to 0.15 while for binary traits it is set to 0.2. As an example, the *ABF* for the hypothesis that all *m* traits colocalize at genetic variant 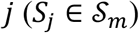 is given by,

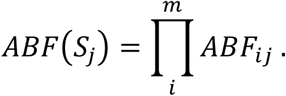

### Calculating Bayes Factors: non-independent studies

If the trait associations are not calculated using independent studies i.e. there are overlapping samples, the Bayes factor for each causal configuration can be computed using a Joint *ABF* (*JABF*) (**Supplementary Material**). The *JABF* for causal configuration *S* is given by,

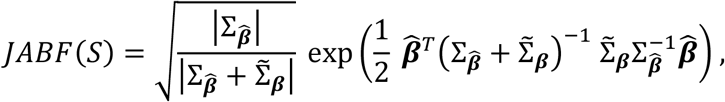

where 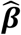 is the vector of regression coefficients for all *m* traits, 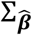 is an *m* × *m* variance-covariance matrix of the regression coefficients (i.e. 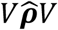, where *V*^2^ is a diagonal matrix of variances for the regression coefficients, e.g. with *i* ^th^ diagonal element 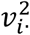, and 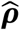 is the observed correlation matrix for the regression coefficients) and 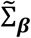 is the ‘adjusted’ prior variance-covariance matrix (i.e. 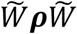, where 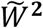 is a diagonal matrix of prior variance divided by estimated variance, e.g. with *i* ^th^ diagonal element 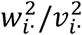, and ***ρ*** is the prior correlation matrix between traits). The correlation matrix 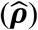 is computed using the tetrachoric correlation method^45^ and we discuss our approach to setting ***ρ*** in the **Supplementary Material**.

### Configuration prior probabilities

We consider two different strategies for determining the priors for different hypotheses: variant-level priors and uniform priors.

### Variant-level prior probabilities

The prior probability space for a single genetic variant can be fully partitioned into the prior probability that the genetic variant is not associated with any of the *m* traits, *p*_0_, the prior probability that the genetic variant is associated with only the first trait, *p*_1_,…, the prior probability that the SNP is associated with a subset of *k* traits {*j*_1_, *j*_2_, …, *j*_*k*_}, 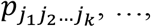 the prior probability that the genetic variant is associated with all traits, *p*_12…*m*_. Hence,

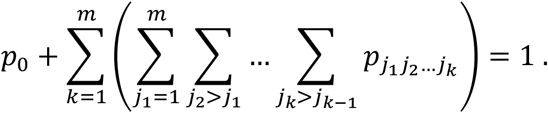

Following^2,8^ we set that the prior probability to not vary by genetic variant, nor by the specific collection of colocalized traits of a given size, but by the number of colocalized traits, i.e. a SNP associated with a total of *k* traits has a prior probability that depends on the number *k* but not the specific collection of traits. To allow for the assessment of large numbers of traits we propose variant-level priors where the prior probability that a genetic variant is associated with *k* traits is given by,

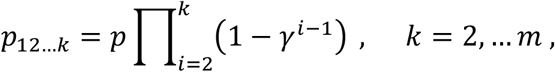

where *p* is the probability of the genetic variant being associated with one trait and *γ* is a parameter which controls the probability that a genetic variant is associated with an additional trait. Notably, 1 − *γ* is the probability of a variant being causal for a second trait given it is causal for one trait, 1 − *γ*^2^ is the probability it is causal for a third trait given it is causal for two traits, and so on. It follows that,

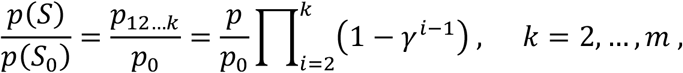

for configurations 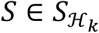, where *k* traits share a causal variant and the remaining *m* − *k* traits do not have a casual variant, and

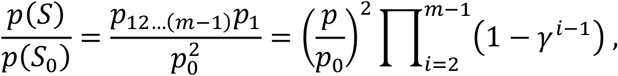

for configurations 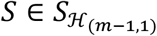, where *m* − 1 traits share a causal variant and the remaining trait has a distinct causal variant. This prior set-up allows evidence to grow in favour of *k* traits colocalizing conditional on evidence supporting *k* − 1 traits colocalizing (**Supplementary Material**). For example, if the first *k* traits are believed to share a causal variant *a priori*, then the prior probability that the (*k* + 1)^*th*^ is also colocalized, conditional on the other *k* traits, increases as the number of colocalized traits *k* grows. The marginal prior probability of *k* traits colocalizing is always very small, however, which controls the false positive rate (**Figures 6** and **S3; Supplementary Tables S2-3**). Conditional growth limits the loss of power when assessing colocalization across a large number of traits. A loss in power *necessarily* occurs when analysing large numbers of colocalized traits, due to the rapid growth in the number of hypotheses in which a subset of traits can colocalize relative to all traits colocalizing. Evidence supporting these ‘subset’ hypotheses will eventually overwhelm evidence in favour of the maximum number of truly colocalized traits for fixed sample size (**Figure 5A**).

### Conditionally uniform prior probabilities

An alternative prior strategy is to assume uniform priors for each configuration within a hypothesis^46^. This strategy benefits from: (i) not setting variant-level information and (ii) implicitly accounting for large differences in the causal configuration space between hypotheses, which limits the loss in power of the *PPFC* for very large *m*. These priors take the form,

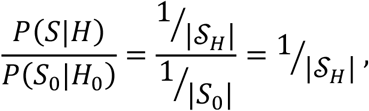

where 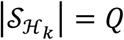 and

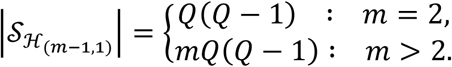

Through simulations, we identified the conditionally uniform prior as less conservative than variant-level priors, having an increased false detection rate of colocalization. (**Supplementary Material; Figures S2-4**). This could lead to an increased false positive detection rate in practice.

### HyPrColoc posterior approximation

To compute the posterior probability of full colocalization across a large number of traits we propose the HyPrColoc posterior approximation. Let *P*(*H*_*m*_|*D*), *P*_*scν*_, *P*_(*m*−1,1)_ and *P*_*all*_ denote: (i) the *posterior probability* of *full colocalization*; (ii) the sum of the posterior probabilities in which no traits have a causal variant, a subset of *m* − 1 traits *share* a *causal variant* (the remaining trait does not have a causal variant) and all *m* traits colocalize (*P*_*scν*_); (iii) the sum of posterior probabilities in which a subset of *m* − 1 traits share a causal variant and the remaining trait has a *distinct* causal variant (*P*_(*m*−1,1)_) and; the sum of *all* posterior probabilities of at most one causal variant per trait (*P*_*all*_). That is,

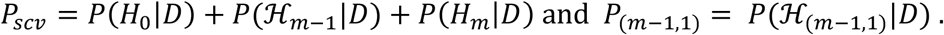

The HyPrColoc posterior is computed in two steps. Step 1 computes the regional association probability *P*_*R*_, defined as:

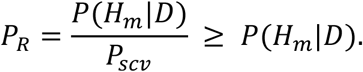

Step 2 computes the alignment probability *P*_*A*_, defined as:

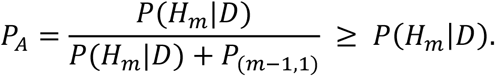

Note that *P*_*R*_ is computed using (*m* + 1)*Q* causal configurations and *P*_*A*_ is computed using an additional *mQ*(*Q* − 1) causal configurations. Hence, computation of *P*_*R*_ and *P*_*A*_ has 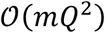 computational cost. We let 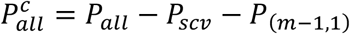, then it follows that the posterior probability of all traits sharing a single causal variant is given by

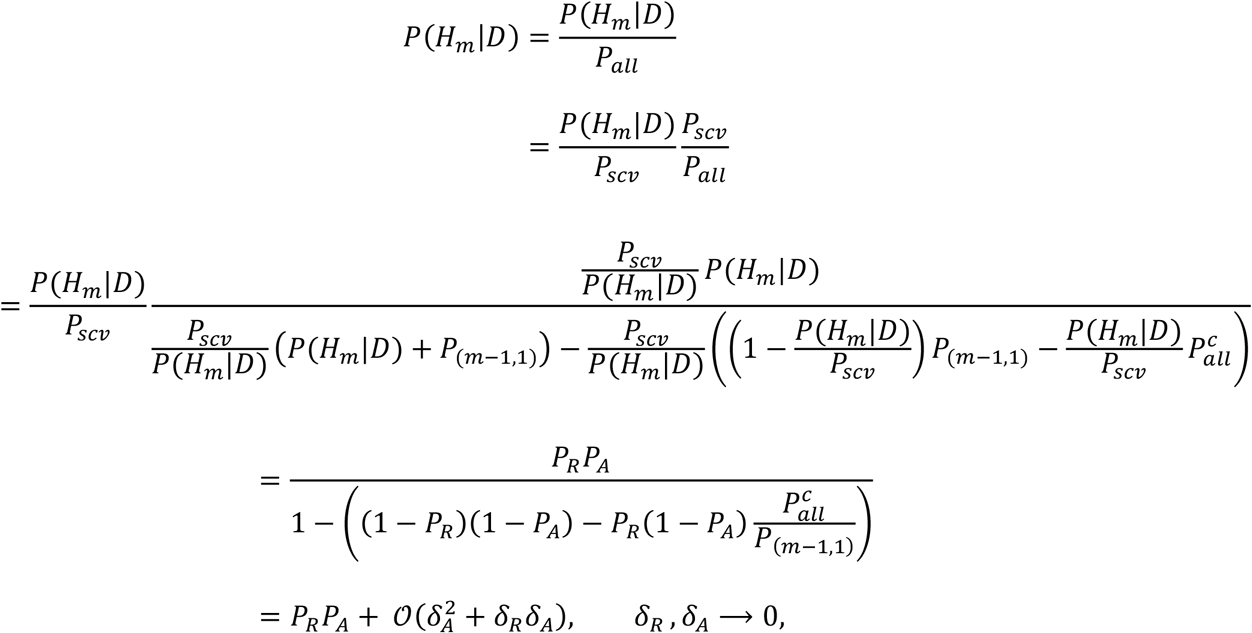

where *δ*_*R*_ = 1 − *P*_*R*_, *δ*_*A*_ = 1 − *P*_*A*_ and

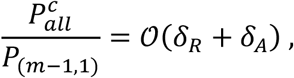

(**Supplementary Material**). By definition, *P*(*H*_*m*_|*D*) → 1 ⇔ *P*_*R*_ → 1 and *P*_*A*_ → 1. Hence together the regional and alignment probabilities when multiplied form a statistic that is sufficient to accurately assess evidence of the full colocalization hypothesis. The objects *P*_*R*_ and *P*_*A*_ can be defined for various collections of hypotheses that partition *P*_*all*_. However, the major insight is that the hypotheses contained in *P*_*R*_ and *P*_*A*_ are computed with minimal computation burden, i.e. computed using ≤ *mQ*^2^ causal configurations, amongst all alternatives, making the HyPrColoc approximation tractable for very large numbers of traits *m*. Our software allows for the assessment of the HyPrColoc approximation by increasing the number of hypotheses used to approximate *P*_*R*_, e.g. we can compute

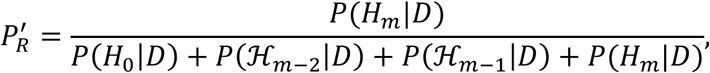

which is computed from 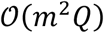 causal configurations and assess the relative difference between *P*_*R*_ and 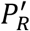. We show that 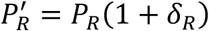 (**Supplementary Material**) and through simulations that there very close correspondence between 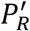 and *P*_*R*_ (**Supplementary table S4**).

### Branch and Bound divisive clustering algorithm

To identify complex patterns of colocalization amongst all traits, we propose a branch and bound (BB) divisive clustering algorithm that utilizes the HyPrColoc approximation to identify a cluster of traits with the greatest evidence of colocalization at each iteration (Figure 3 and **Supplementary Material**). Starting with all of the traits in a single cluster, the algorithm explores evidence supporting any of 2*m* branches - a branch represents a hypothesis whereby *m* − 1 traits share a causal variant and either the remaining trait does not have a causal variant or has a causal variant elsewhere in the region - against the full colocalization hypothesis. These branches represent the hypotheses used in the computation of the regional and alignment probabilities *P*_*R*_ and *P*_*A*_. There are two bounds: (i) the minimum probability required to accept evidence that all *m* traits are regionally associated 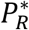 and (ii) the minimum probability required to accept that the causal variant for all *m* traits aligns at a single variant 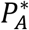. The BB algorithm accepts evidence supporting all *m* traits sharing a single causal variant if 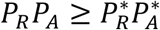, after which the algorithm returns the HyPrColoc estimate of *PPFC* and stops. If either 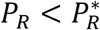 or 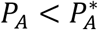 there is insufficient evidence supporting all traits sharing a causal variant and the BB algorithm moves to the branch with maximum evidence supporting *m* − 1 traits sharing a causal variant. At this point the traits are partitioned into two clusters: one containing *m* − 1 traits deemed most likely to share a causal variant and a second cluster containing the remaining trait. We repeat this process of branch selection and partitioning on the cluster of *m* − 1 traits until we identify either: (A) a cluster of traits of size *k* ≥ 2 whose regional and alignment statistics satisfy 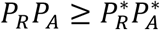, or (B) there is one trait left in the cluster. In scenario A, the HyPrColoc posterior probability that all *k* traits colocalize is presented and the remaining *m* − *k* traits are assessed for evidence of colocalization using the branch selection and partitioning scheme. In scenario B, the trait is deemed not colocalize with any other trait in the sample and the BB selection algorithm is repeated using *m* − 1 traits. The entire process is repeated until all clusters of colocalized traits, whereby each cluster of traits colocalize at a distinct causal variant, have been identified, all other traits are deemed not to share a causal variant with any other trait.

### Simulation study

To create genomic loci with realistic patterns of LD, for each simulation scenario we simulated 1,000 datasets and for each dataset we resampled phased haplotypes from the European samples in 1000 Genomes^14^ and randomly chose one of the first 50 regions confirmed to be associated with CHD^15^. Unless stated otherwise, for traits that have a causal variant in the region, the variant explains 1% of trait variance and each trait was assumed to be measured in studies with a sample size of *N* = 10,000. Variant-level priors were chosen for the simulation study with the stringent choice of *γ* = 0.98 and setting *p* = 10^−4^ as in^2^.

### Application to CHD and cardiovascular risk factors

The GWAS results used in the assessment of colocalization of CHD with related traits were taken from large-scale analyses of CHD^16^, blood pressure (http://www.nealelab.is/uk-biobank), adiposity measures (http://www.nealelab.is/uk-biobank), glycaemic traits^17^, renal function^18^, type II diabetes^19^, lipid measurements^20^, smoking^21^, rheumatoid arthritis^22^ and educational attainment^23^ (**Table S1**). All datasets had either been imputed to 1000 Genomes^14^ prior to GWAS analyses or were imputed up to 1000 Genomes from the summary results using DIST^47^ (INFO>0.8). We performed colocalization analyses in two steps. In step one, we assessed colocalization of CHD with the 14 risk-factors in pre-specified LD blocks from across the genome^24^. We used a conservative variant-level prior structure with *p* = 1 × 10^−4^ and *γ* = 0.95, i.e. 1 in 200,000 variants are expected to be causal for two traits, and set strong bounds for the regional and alignment probabilities, i.e. 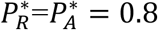 so that the algorithm identified a cluster of colocalized traits only if *P*_*R*_*P*_*A*_ > 0.64. The full results from this analysis are available at https://jrs95.shinyapps.io/hyprcoloc_chd.

To prioritise candidate causal genes in regions where CHD and at least one related trait colocalized, we re-ran the colocalization analysis and included whole blood *cis* eQTL^26^ (31,684 samples) and *cis* pQTL^27^ (3,301 samples) data in addition to the primary traits, in a second step. A colocalization analysis was performed for every transcript with data within each region. *cis* eQTL were defined 1MB upstream and downstream of the centre of the gene probe (1,828 genes were analysed across the 43 regions). *cis* pQTL were defined 5MB upstream and downstream of the transcript start site (854 proteins were analysed across the 43 regions). We integrated gene expression information taken from whole blood tissue as: (i) the eQTLGen dataset^26^ has a large sample size relative to other publicly available gene expression data resources and; (ii) the pQTL data were also measured in whole blood tissues, so there was consistency in the tissue analysed.

## References

1. Nica A. C. & Dermitzakis E. T. Using gene expression to investigate the genetic basis of complex disorders. Hum. Mol. Genet. 17, 129–134 (2008).

2. Giambartolomei C. et al. Bayesian Test for Colocalisation between Pairs of Genetic Association Studies Using Summary Statistics. PLoS Genet. 10, (2014).

3. Guo H. et al. Integration of disease association and eQTL data using a Bayesian colocalisation approach highlights six candidate causal genes in immune-mediated diseases. Hum. Mol. Genet. 24, 3305–3313 (2015).

4. Hauberg M. E. et al. Large-Scale Identification of Common Trait and Disease Variants Affecting Gene Expression. Am. J. Hum. Genet. 100, 885–894 (2017).

5. Hormozdiari F. et al. Colocalization of GWAS and eQTL Signals Detects Target Genes. Am. J. Hum. Genet. 99, 1245–1260 (2016).

6. Wen X., Pique-Regi R. & Luca F. Integrating molecular QTL data into genome-wide genetic association analysis: Probabilistic assessment of enrichment and colocalization. PLoS Genet. 13, 1–25 (2017).

7. Jaffe A. et al. Mapping DNA methylation across development, genotype, and schizophrenia in the human frontal cortex. Nat. Neurosci. 19, 40–47 (2016).

8. Giambartolomei C. et al. A Bayesian framework for multiple trait colocalization from summary association statistics. Bioinformatics 34, 2538–2545 (2018).

9. Plagnol V., Smyth D. J., Todd J. A. & Clayton D. G. Statistical independence of the colocalized association signals for type 1 diabetes and RPS26 gene expression on chromosome 12q13. Biostatistics 10, 327–334 (2009).

10. Wallace C. et al. Statistical colocalization of monocyte gene expression and genetic risk variants for type 1 diabetes. Hum. Mol. Genet. 21, 2815–2824 (2012).

11. Hippisley-Cox J. et al. Predicting cardiovascular risk in England and Wales: Prospective derivation and validation of QRISK2. Bmj 336, 1475–1482 (2008).

12. Rodondi N. et al. Framingham Risk Score and Alternatives for Prediction of Coronary Heart Disease in Older Adults. 7, (2012).

13. Wakefield J. Bayes Factors for Genome-Wide Association Studies: Comparison with P-values. 86, 79–86 (2009).

14. The 1000 Genomes Project Consortium. A global reference for human genetic variation. Nature 526, 68–74 (2015).

15. The CARDIoGRAMplusC4D Consortium. Large-scale association analysis identifies new risk loci for coronary artery disease. Nat. Genet. 45, 25–33 (2012).

16. Nikpay M., Goel A., Won H.-H. & Hall L. M. A comprehensive 1000 Genomes-based genome-wide association meta-analysis of coronary artery disease. Nat. Genet. 47, 1121–1130 (2015).

17. Dupuis J. et al. New genetic loci implicated in fasting glucose homeostasis and their impact on type 2 diabetes risk. Nat Genet 42, 105–116 (2010).

18. Gorski M. et al. 1000 Genomes-based meta-analysis identifies 10 novel loci for kidney function. Sci. Rep. 7, 1–10 (2017).

19. Scott R. A. et al. An Expanded Genome-Wide Association Study of Type 2 Diabetes in Europeans. Diabetes 66, 2888–2902 (2017).

20. Teslovich T. M. et al. Biological, Clinical, and Population Relevance of 95 Loci for Blood Lipids. Nature 466, 707–713 (2010).

21. The Tobacco and Genetics Consortium. Genome-wide meta-analyses identify multiple loci associated with smoking behavior. Nat. Genet. 42, 441–447 (2010).

22. Okada Y. et al. Genetics of rheumatoid arthritis contributes to biology and drug discovery. Nature 113, 190–196 (2014).

23. Okbay A., Beauchamp J. P., Fontana M. A., Lee J. J. & Pers T. H. Genome-wide association study identifies 74 loci associated with educational attainment. Nature 533, 539–542 (2016).

24. Berisa T. & Pickrell J. K. Approximately independent linkage disequilibrium blocks in human populations. Bioinformatics 32, 283–285 (2015).

25. Van Der Harst P. & Verweij N. Identification of 64 novel genetic loci provides an expanded view on the genetic architecture of coronary artery disease. Circ. Res. 122, 433–443 (2018).

26. Võsa U. et al. Unraveling the polygenic architecture of complex traits using blood eQTL meta-analysis. bioRxiv 18, 10 (2018).

27. Sun B. B. et al. Genomic atlas of the human plasma proteome. Nature 558, 273–79 (2018).

28. Forouzanfar M. H. et al. Global burden of hypertension and systolic blood pressure of at least 110 to 115mmHg, 1990-2015. JAMA - J. Am. Med. Assoc. 317, 165–182 (2017).

29. Howson J. M. M., Zhao W. & Barnes D. R. Fifteen new risk loci for coronary artery disease highlight arterial wall-specific mechanisms. Nat Genet 49, 1113–1119 (2017).

30. Nelson C. P. et al. Association analyses based on false discovery rate implicate new loci for coronary artery disease. Nat. Genet. 49, 1385–1391 (2017).

31. The IBC 50K CAD Consortium. Large-scale gene-centric analysis identifies novel variants for coronary artery disease. PLoS Genet. 7, (2011).

32. The Coronary Artery Disease (C4D) Genetics Consortium. A genome-wide association study in Europeans and South Asians identifies five new loci for coronary artery disease. Nat. Genet. 43, 339–346 (2011).

33. Klarin D. et al. Genetic Analysis in UK Biobank Links Insulin Resistance and Transendothelial Migration Pathways to Coronary Artery Disease. Nat Genet 49, 1392–1397 (2017).

34. Schunkert H. et al. Large-scale association analyses identifies 13 new susceptibility loci for coronary artery disease. Nat Genet 43, 333–338 (2011).

35. Turley P. et al. Multi-trait analysis of genome-wide association summary statistics using MTAG. Nat Genet 50, 229–237 (2018).

36. International Consortium for Blood Pressure Genome-Wide Association Studies. Genetic Variants in Novel Pathways Influence Blood Pressure and Cardiovascular Disease Risk. Nature 478, 103–109 (2011).

37. GTEx Consortium. Genetic effects on gene expression across human tissues. Nature 550, 204–213 (2017).

38. Kessler T., Wobost J., Wolf B., Eckhold J. & Vilne B. Functional characterization of the GUCY1A3 coronary artery disease risk locus. Circulation 136, 476–489 (2017).

39. Erdmann J., Kessler T., Venegas L. M. & Schunkert H. A decade of genome-wide association studies for coronary artery disease : the challenges ahead. Cardiovasc. Res. 49, 1241–1257 (2018).

40. Gertow K. et al. Identification of the BCAR1-CFDP1-TMEM170A Locus as a Determinant of Carotid Intima-Media Thickness and Coronary Artery Disease Risk. Circ. Cardiovasc. Genet. 5, 656–665 (2012).

41. Boardman-Pretty F. et al. Functional Analysis of a Carotid Intima-Media Thickness Locus Implicates BCAR1 and Suggests a Causal Variant. Circ. Cardiovasc. Genet. 8, 696–706 (2015).

42. Webb T. R. et al. Systematic Evaluation of Pleiotropy Identifies 6 Further Loci Associated With Coronary Artery Disease. J. Am. Coll. Cardiol. 69, 735–1097 (2017).

43. Nelson M. R. et al. The support of human genetic evidence for approved drug indications. Nat. Genet. 47, 856–860 (2015).

44. Benner C. et al. Prospects of Fine-Mapping Trait-Associated Genomic Regions by Using Summary Statistics from Genome-wide Association Studies. Am. J. Hum. Genet. 101, 539–551 (2017).

45. Province M. A. & Borecki I. B. A correlated meta-analysis strategy for data mining ‘OMIC’ scans. Pac. Symp. Biocomput. 236–46 (2013).

46. Pickrell J. K. et al. Detection and interpretation of shared genetic influences on 42 human traits. Nat Genet 48, 709–717 (2016).

47. Lee D., Bigdeli T. B., Riley B. P., Fanous A. H. & Bacanu S. A. DIST: Direct imputation of summary statistics for unmeasured SNPs. Bioinformatics 29, 2925–2927 (2013).

48. McLaren W. et al. The Ensembl Variant Effect Predictor. Genome Biol. 17, 1–14 (2016).

49. Staley J. R. et al. PhenoScanner: A database of human genotype-phenotype associations. Bioinformatics 32, 3207–3209 (2016).

