## Supplementary_material for "A fast and efficient colocalization algorithm for identifying shared genetic risk factors across multiple traits"

### Overview

*HyPrColoc* is a model and algorithm that uses summary association statistics to assess whether multiple traits colocalize. Given a genomic region and a (potentially large) collection of traits, *HyPrColoc* seeks to identify clusters of traits, such that Trait A and Trait B are allocated to the same cluster if and only if they share a causal variant within the region. To do this, *HyPrColoc* uses a novel model-based Bayesian divisive clustering algorithm. The algorithm starts with all traits in the same cluster, and iteratively divides the cluster into sub-clusters. Whether or not a cluster should be divided into sub-clusters is determined using a Bayesian model selection approach, which requires the calculation of (approximate) Bayes factors. A key quantity that must often be calculated by the algorithm is the posterior probability that a collection of $m$ traits all share a causal variant within a genomic region (we refer to this quantity as the *posterior probability of full co-localization*, or PPFC, across the $m$ traits). We derive a tightly bounded approximation of this quantity, which greatly accelerates computation and enables *HyPrColoc* to scale to large numbers of traits.

**Outline**

We start in Section S1 by providing the mathematical notation that will be used throughout. In Section S2, we provide details of the Bayesian model selection approach, which requires a number of novel mathematical derivations in order to allow us to efficiently calculate the posterior probability associated with different colocalization hypotheses. In particular: in Section S2.1, we derive expressions for Joint Approximate Bayes Factors (JABFs); in Section S2.2. we consider strategies for specifying the prior probabilities associated with different causal hypotheses; and in Section S2.3 we introduce the idea of *hypothesis prioritisation*, as well as deriving the *HyPrColoc* approximation of the PPFC, which allows *HyPrColoc* to scale to large numbers of traits. In Section S3, we describe the *HyPrColoc* Bayesian divisive clustering algorithm. Finally, in Section S4 we provide further simulation results and mathematical details.

### S1. Notation

Let $D = \{D_1, D_2, \ldots, D_m\}$ be a set of datasets obtained from $m$ studies (some or all may contain overlapping participants), where $D_i = \{\boldsymbol{y}_i, \boldsymbol{X}_i, \boldsymbol{G}_i\}$ is the combined dataset comprising: outcome (trait), covariate, and genetic data from study $i$. More precisely, if the $i^{th}$ dataset comprises measurements on $n_i$ participants, then:

- $\boldsymbol{y}_i$ is an $n_i$ dimensional vector of outcome data;
- $\boldsymbol{X}_i$ is a $p_i \times n_i$ matrix of covariate data, where $p_i$ is the number of covariates measured in the $i^{th}$ dataset; and
- $\boldsymbol{G}_i$ is a $Q \times n_i$ matrix of genotype data, where $Q$ is the number of SNPs.

Note that the number of covariates, $p_i$, may vary by study, however, the number of SNPs, $Q$, in the genomic region is fixed and constant across studies.

Let $\boldsymbol{X}_{ij}$ and $\boldsymbol{G}_{ij}$ be the $j^{th}$ column vectors for the covariate and genotype data respectively, then we assume the following outcome model for individual $j$ in study $i$

$$\mathbb{E}[y_{ij}] = h_i^{-1}\left(\boldsymbol{\alpha}_i^T \boldsymbol{X}_{ij} + \boldsymbol{\beta}_i^T \boldsymbol{G}_{ij}\right), \quad i = 1,2,\ldots m \quad \text{and} \quad j = 1,2,\ldots,n_i,$$

( 1 )

where

$h_i$ is the link function for study $i$,
$\boldsymbol{\alpha}_i$ is a $p_i$ dimensional vector of (nuisance) covariate effects,
$\boldsymbol{\beta}_i$ is a $Q$ dimensional vector of genotype effects.

The choice of link function, $h_i$, will depend on the type of outcome, $y_i$, in the $i^{th}$ dataset. For example, if $y_i$ is continuous, e.g. gene expression levels, then $h_i$ might be the identity function; or if $y_i$ is binary, e.g. a disease outcome, then $h_i$ might be a logit model.

The aim of *HyPrColoc* is to assess evidence supporting colocalization across clusters of traits, under the assumption that each of the $m$ traits has a maximum of one causal variant in a genomic region. In order to do this, *HyPrColoc* explores different hypotheses regarding: (i) which variants may be causal; and (ii) which traits may share the same causal variant.

For simplicity, here we only consider a single genomic region. Let $Q$ be the number of SNPs in the region of interest. A colocalization hypothesis $H$ can be expressed in terms of a colocalization mechanism, *i.e.* no causal variant (CV), one CV, or several CVs in the region. Moreover, a CV can be uniquely associated with just one trait or can be shared by a subset of the $m$ traits, including being shared across all $m$-traits. We refer to a CV shared across all $m$ traits as '*full colocalization*'. For each trait, SNP causality can be summarised by a binary vector of length $Q$, where a 1 at the $j^{th}$ position indicates that the $j^{th}$ variant is causally associated with the trait. A collection of $m$ such vectors, one per trait, defines an $m \times Q$ *causal configuration* matrix $S$, where element $(i, j)$ defines the causal status of SNP $j$ for trait $i$, i.e.

$$\{S\}_{ij} = \begin{cases} 1, & \text{SNP } j \text{ causal for trait } i \\ 0, & \text{SNP } j \text{ not causal for trait } i \end{cases}.$$

For example, if $H_m^{(j)}$ is the hypothesis that *the* $j^{th}$ *SNP is causal for all m traits*, then the corresponding configuration matrix, $S_j$, is a matrix with 1s in the $j^{th}$ column, and 0s elsewhere:

$$S_j = \underbrace{\begin{pmatrix} 0 & \cdots & 1 & 0 & \cdots & 0 \\ 0 & \cdots & 1 & 0 & \cdots & 0 \\ \vdots & \cdots & \vdots & \vdots & \ddots & \vdots \\ 0 & \cdots & 1 & 0 & \cdots & 0 \end{pmatrix}}_{j^{th} \text{ column is } 1}$$

In practice, most hypotheses correspond to multiple configuration matrices, e.g. if $H_m$ is the hypothesis that *there is a SNP in the region that is causal for all m traits*, then $H_m$ corresponds to the set of configuration matrices, $\mathcal{S}_{H_m} = \{ S_j \} : j = 1,2, \dots, m,$ , where each $S_j$ is as defined above. We denote by $H_0$ the null hypothesis that *no variant in the region is causal for any of the traits*, and note that $H_0$ corresponds to a single configuration matrix $S_0$, which is the $m \times Q$ matrix of zeros.

#### S2. Bayesian model selection for colocalization hypotheses

As in [1], we adopt a Bayesian model selection approach for choosing between different colocalization hypotheses. As outlined above, any colocalization hypothesis $H$ can be specified by a set of causal configurations, $\mathcal{S}_H$. Consequently, the posterior probability (PP) of $H$ can be written as

$$P(H|D) = \frac{P(D|H)P(H)}{P(D)}$$

$$= \frac{\sum_{S \in \mathcal{S}_H} P(D|S)\, P(S)}{\sum_{H^* \in \Omega} \sum_{S \in \mathcal{S}_{H^*}} P(D|S)\, P(S)},$$

where Ω denotes the space of all possible colocalization hypotheses (discussed further in Section S2.3).

In practice, it is useful to rewrite the right-hand side of the above equation in terms of the Bayes Factor *BF*(*S*) in favour of configuration *S* relative to the null configuration,

$$BF(S) = \frac{P(D|S)}{P(D|S_0)},$$

which is the (marginal) likelihood associated with the data being generated under the causal configuration $S$, relative to the (marginal) likelihood associated with the data being generated under the null configuration $S_0$.

Dividing the numerator and denominator in the equation above by $P(H_0|D) = \frac{P(D|S_0)P(S_0)}{P(D)}$, i.e. the posterior of the null-hypothesis $H_0$, we obtain

$$P(H|D) = \frac{\sum_{S\in\mathcal{S}_H} BF(S)\frac{P(S)}{P(S_0)}}{\sum_{H^*\in\Omega}\sum_{S\in\mathcal{S}_{H^*}} BF(S)\frac{P(S)}{P(S_0)}}.$$

( 2 )

In the remainder of this section, we consider how the various terms in Equation (2) above can be efficiently evaluated or estimated. In Section S2.1, we extend the work of [2] to derive approximations for the Bayes factors, *BF*(*S*). In Section S2.2, we consider strategies for specifying the causal configuration priors, *P*(*S*). Finally, in Section S2.3 we derive a novel hypothesis prioritization (*HyPr*) approach, which enables us to efficiently evaluate the denominator in Equation (2) and thereby allows *HyPrColoc* to scale to large numbers of traits.

#### S2.1. Joint approximate Bayes factors

In practice, individual participant data are often unavailable (and so $D$ is unknown), and we must work instead with study-level summary data. In this case, the Bayes Factor *BF*(*S*) is also unavailable, but we may instead calculate an *Approximate Bayes Factor* (ABF) using $\hat{\beta}_{j_i}$, the estimated effect of the $j$th SNP on the outcome in the $i$th study, and its associated standard error, $\hat{V}_{j_i}$ [2]. Here we extend the approach of [2] and derive a *joint* ABF (*JABF*) for all $m$ studies under causal configuration $S, JABF(S)$, which allows for overlapping samples between studies.

We assume in our derivations that there are $Q$ variants in the region and a maximum of $q = 1$ variants are causal per trait. We denote the number of CVs that trait $i$ has under configuration $S$ by $q_{i_s} : q_{i_s} \in \{0,1\}$ and define $q_s = \sum_{i=1}^{m} q_{i_s}$. To simplify notation, we assume without loss of generality that the first $i = 1,2, \dots, q_s$ traits have a casual variant under configuration $S$. Moreover, to simplify our discussion we assume that the $k^{th}$ variant is causal for traits $i \leq q_s$. Under configuration $S$, the linear predictor in Equation (1) may be written in terms of a $p_i$ dimensional vector of covariate effects $\boldsymbol{\alpha}_{i_s}$ and a parameter $\beta_{k_i}$ denoting the genetic effect of the $k^{th}$ variant, i.e.

$$\mathbb{E}[y_{ij}] = h_i^{-1}\left(\boldsymbol{\alpha}_{i_s}^T \boldsymbol{X}_{ij} + \beta_{k_i} G_{ijk}\right),$$

where $G_{ijk}$ denotes the $k^{th}$ genotype for participant $j$ in the $i^{th}$ study. For studies $i > q_s$ the genetic effect $\beta_{k_i} = 0$. Recall that $D$ denotes the combined data across all $m$ studies. It follows, therefore, that

$$P(D|S) = \int \int P(D|\,\boldsymbol{\alpha}, \boldsymbol{\beta}, S)\pi(\boldsymbol{\alpha}, \boldsymbol{\beta}|S) d\boldsymbol{\alpha} d\boldsymbol{\beta}$$

$$\approx \int \int P\left(\widehat{\boldsymbol{\alpha}}, \widehat{\boldsymbol{\beta}}|\,\boldsymbol{\alpha}, \boldsymbol{\beta}, S\right)\pi(\boldsymbol{\alpha}, \boldsymbol{\beta}|S) d\boldsymbol{\alpha} d\boldsymbol{\beta} \;:\quad n_i \gg 1\,,\;\; i = 1,2, \dots m,$$

where:

- $\pi$ denotes the prior distribution for the covariate and genetic effect parameters.
- $\{\widehat{\boldsymbol{\alpha}}, \widehat{\boldsymbol{\beta}}\}$ denote the *joint* summary data for each of the $m$ studies under configuration $S$.

The validity of the approximation in the second line of the equation above relies on each study having a large sample size, a situation that is almost always satisfied in GWAS.

Using standard results about asymptotic distributions of maximum likelihood estimators, it follows that – in the asymptotic limit – the sampling distribution for $\{\widehat{\boldsymbol{\alpha}}, \widehat{\boldsymbol{\beta}}\}$ is then:

$$\begin{pmatrix}\widehat{\boldsymbol{\alpha}}\\ \widehat{\boldsymbol{\beta}}\end{pmatrix} \mid S \sim N_{p+m}\left(\begin{pmatrix}\boldsymbol{\alpha}_s\\ \boldsymbol{\beta}_s\end{pmatrix}, I^{-1}\right), \quad n_i \longrightarrow \infty, \quad i = 1,2,\dots m,$$

where $p = \sum_{i=1}^{m} p_i$, $\boldsymbol{\alpha}_s = \mathbb{E}[\widehat{\boldsymbol{\alpha}}|S]$, $\boldsymbol{\beta}_s = \mathbb{E}[\widehat{\boldsymbol{\beta}}|S]$ and $I$ is the Fisher information which may be written as

$$I = \begin{pmatrix} \overbrace{I_{00}}^{p\times p} & \overbrace{I_{01}}^{p\times m} \\ \underbrace{(I_{01})^T}_{m\times p} & \underbrace{I_{11}}_{m\times m} \end{pmatrix}$$

If the covariate $\boldsymbol{X}_{ij}$ and genotypic information $\boldsymbol{G}_{ij}$, conditional on configuration $S$, are independent for each of the $i = 1,2,\dots,m$ studies then the information $I_{01} = 0$. However, this is unlikely to be true in general. In what follows, we therefore generalise the approach of [2] to the dependent case. Following [2], we first define the re-parameterisation matrix A to be

$$A = \begin{pmatrix} \mathbb{I}_p & (I_{00})^{-1} I_{01} \\ \boldsymbol{0} & \mathbb{I}_m \end{pmatrix},$$

where $\mathbb{I}_r$ denotes the $r \times r$ identity matrix.

Applying $A$ to $\begin{pmatrix}\widehat{\boldsymbol{\alpha}}\\ \widehat{\boldsymbol{\beta}}\end{pmatrix}$, we obtain

$$A\begin{pmatrix}\widehat{\boldsymbol{\alpha}}\\ \widehat{\boldsymbol{\beta}}\end{pmatrix} \mid S = \begin{pmatrix}\widehat{\boldsymbol{\tau}}\\ \widehat{\boldsymbol{\beta}}_s\end{pmatrix} \sim N_{p+m}\left(\begin{pmatrix}\boldsymbol{\tau}\\ \boldsymbol{\beta}_s\end{pmatrix}, \begin{bmatrix}\Sigma_{\widehat{\boldsymbol{\tau}}} & \boldsymbol{0}\\ \boldsymbol{0}^T & \Sigma_{\widehat{\boldsymbol{\beta}}_s}\end{bmatrix}\right),$$

where we have used $\widehat{\boldsymbol{\beta}}_s$ as shorthand to denote the $m$ dimensional vector of estimated genetic effects under configuration $S$,

$$\widehat{\boldsymbol{\beta}}_s = \{\hat{\beta}_{k_1}, \hat{\beta}_{k_2}, \dots, \hat{\beta}_{k_m}\}$$

with $\hat{\beta}_{k_i}$ denoting the estimated effect of the $k^{th}$ SNP on the $i^{th}$ trait and $\Sigma_{\widehat{\boldsymbol{\beta}}_s}$ denoting the estimated $m \times m$ covariance matrix of the combined study effect vector $\widehat{\boldsymbol{\beta}}_s$,

$$\Sigma_{\widehat{\boldsymbol{\beta}}_s} = \begin{pmatrix} \hat{V}_{k_1}^2 & \cdots & \widehat{\sigma}_{k_1 k_m} \\ \vdots & \ddots & \vdots \\ \widehat{\sigma}_{k_1 k_m} & \cdots & \hat{V}_{k_m}^2 \end{pmatrix},$$

where $\widehat{\sigma}_{k_j k_l}$ denotes the covariance between the estimated effects $\hat{\beta}_{k_j}$ and $\hat{\beta}_{k_l}$. If the studies do not contain overlapping participants, then $\Sigma_{\widehat{\boldsymbol{\beta}}_s}$ is a diagonal matrix whose $i^{th}$ diagonal element is the estimated variance $\left(\hat{V}_{k_i}^2\right)$. The (marginal) likelihood $P(D|S)$ then becomes

$$P(D|S) \approx \int\int P\left(\widehat{\boldsymbol{\alpha}}, \widehat{\boldsymbol{\beta}} \mid \boldsymbol{\alpha}, \boldsymbol{\beta}, S\right)\pi(\boldsymbol{\alpha}, \boldsymbol{\beta}|S)d\boldsymbol{\alpha}d\boldsymbol{\beta}$$

$$= |A| \int P(\hat{\boldsymbol{\tau}} \mid \boldsymbol{\tau})\pi(\boldsymbol{\tau})d\boldsymbol{\tau} \int P\left(\widehat{\boldsymbol{\beta}}_s \mid \boldsymbol{\beta}_s\right)\pi(\boldsymbol{\beta}_s)d\boldsymbol{\beta}_s, \qquad n_i \gg 1, \;\; i = 1,2, \dots m,$$

where $|A|$ is the Jacobian, resulting from the transformation of variables, and the prior distribution $\pi(\boldsymbol{\tau}, \boldsymbol{\beta}_s) = \pi(\boldsymbol{\tau})\pi(\boldsymbol{\beta}_s)$ assumes $\boldsymbol{\tau}$ and $\boldsymbol{\beta}_s$ are independent. In terms of the re-parameterised model above, it follows that

$$BF(S) = \frac{P(D|S)}{P(D|S_0)} \approx \frac{\int P\left(\widehat{\boldsymbol{\beta}} \mid \boldsymbol{\beta}, S\right)\pi(\boldsymbol{\beta}|S)d\boldsymbol{\beta}}{\int P\left(\widehat{\boldsymbol{\beta}} \mid \boldsymbol{\beta}, S_0\right)\pi(\boldsymbol{\beta}|S_0)d\boldsymbol{\beta}} = \frac{P\left(\widehat{\boldsymbol{\beta}}_s\right)}{P\left(\widehat{\boldsymbol{\beta}}_{s_0}\right)} = JABF(S).$$

If we define the marginal prior distribution on $\boldsymbol{\beta}_s$ to be

$$\boldsymbol{\beta}_s \sim N_m\left(\mathbf{0}\,, \Sigma_{\boldsymbol{\beta}_s}\right)$$

under configuration $S$, it follows that the JABF can be written as

$$JABF(S) = \sqrt{\frac{\left|\Sigma_{\widehat{\boldsymbol{\beta}}_s}\right|}{\left|\Sigma_{\widehat{\boldsymbol{\beta}}_s} + \Sigma_{\boldsymbol{\beta}_s}\right|}}\; \exp\left\{\frac{1}{2}\, \widehat{\boldsymbol{\beta}}_s^{\;T}\left(\Sigma_{\widehat{\boldsymbol{\beta}}_s} + \Sigma_{\widehat{\boldsymbol{\beta}}_s}\Sigma_{\boldsymbol{\beta}_s}^{-1}\Sigma_{\widehat{\boldsymbol{\beta}}_s}\right)^{-1}\widehat{\boldsymbol{\beta}}_s\right\}.$$

We will find it useful later to write the above in terms of $Z$-scores. Let

$$\widehat{\boldsymbol{Z}}_s = \hat{V}_s^{-1}\widehat{\boldsymbol{\beta}}_s : \quad \hat{V}_s = \mathrm{diag}\left\{\hat{V}_{k_1}, \hat{V}_{k_2}, \dots, \hat{V}_{k_m}\right\}$$

be the vector of $Z$-scores for the $k^{th}$ variant in each study. We further define

$$\boldsymbol{Z}_s = W_s^{-1}\boldsymbol{\beta}_s : \quad W_s = \mathrm{diag}\left\{W_{k_1}, W_{k_2}, \dots, W_{k_m}\right\},$$

where $W_j = \sqrt{(\Sigma_{\boldsymbol{\beta}_s})_{jj}}$ is the square root of the $j^{th}$ diagonal element of the covariance matrix $\Sigma_{\boldsymbol{\beta}_s}$.

Then,

$$JABF(S) = \sqrt{\frac{|\Sigma_{\widehat{\boldsymbol{Z}}_s}|}{|\Sigma_{\widehat{\boldsymbol{Z}}_s} + \tilde{\Sigma}_{\boldsymbol{Z}_s}|}} \exp\left\{\frac{1}{2}\, \widehat{\boldsymbol{Z}}_s{}^T \left(\Sigma_{\widehat{\boldsymbol{Z}}_s} + \Sigma_{\widehat{\boldsymbol{Z}}_s} \tilde{\Sigma}_{\boldsymbol{Z}_s}^{-1} \Sigma_{\widehat{\boldsymbol{Z}}_s}\right)^{-1} \widehat{\boldsymbol{Z}}_s\right\},$$

*( 3 )*

where

$$\tilde{\Sigma}_{\boldsymbol{Z}_s} = W_s \Sigma_{\boldsymbol{Z}_s} W_s \quad \text{and} \quad W_s = \text{diag}\left\{\frac{W_{k_1}}{\widehat{V}_{k_1}}, \frac{W_{k_2}}{\widehat{V}_{k_2}}, \dots, \frac{W_{k_m}}{\widehat{V}_{k_m}}\right\}$$

is a diagonal matrix whose elements are ratios of prior standard deviation (i.e. $W_{k_l}$) relative to estimated marginal effect standard deviations for trait $l$ and variant $k$ under configuration $S$. We call $\tilde{\Sigma}_{\boldsymbol{Z}_s}$ the *adjusted* prior covariance matrix, as it contains both estimated and prior information. Equation (3) is the general form for an ABF for a collection of $m \geq 2$ correlated traits.

**The independent studies case**

If all $m$ studies are independent, then both $\Sigma_{\widehat{\boldsymbol{Z}}_s}$ and $\tilde{\Sigma}_{\boldsymbol{Z}_s}$ are diagonal matrices and it follows that the $i^{th}$ element

$$\left\{\Sigma_{\widehat{\boldsymbol{Z}}_s} + \Sigma_{\widehat{\boldsymbol{Z}}_s} \tilde{\Sigma}_{\boldsymbol{Z}_s}^{-1} \Sigma_{\widehat{\boldsymbol{Z}}_s}\right\}_i = \frac{\widehat{V}_{k_i}^2 + W_{k_i}^2}{W_{k_i}^2}$$

which means that the approximate Bayes factor for the $i^{th}$ study, given configuration $S$, is given by

$$ABF_i(S) = \sqrt{\frac{\widehat{V}_{k_i}^2}{\widehat{V}_{k_i}^2 + W_{k_i}^2}} \exp\left\{\frac{1}{2} \hat{Z}_{k_i}^2 \frac{W_{k_i}^2}{\widehat{V}_{k_i}^2 + W_{k_i}^2}\right\},$$

which matches the results[1–3], and

$$JABF(S) = \prod_{i=1}^{m} ABF_i(S).$$

**Specification of the prior on the genetic effects, $\boldsymbol{\beta}$**

To compute a JABF for each causal configuration, $O(m^2Q^2)$ hyperparameters need to be set in order to specify the adjusted prior matrices $\tilde{\Sigma}_{\boldsymbol{Z}_s}$. As the number of studies $m$ increases, this makes prior specification challenging. We propose the following strategy to attenuate the burden of setting such a large amount of hyperparameters.

Recall that the $i^{\text{th}}$ diagonal element of the diagonal matrix $W_s$ is the ratio of the marginal effect prior and sample standard deviation $\frac{W_{k_i}}{\hat{V}_{k_i}}$ from study $i$ and variant $k$, under configuration $S$. If we adopt an empirical Bayes approach and set the prior correlation matrix $\boldsymbol{\rho}(\Sigma_{\boldsymbol{Z}_s})$ equal to a correlation matrix, $\boldsymbol{\rho}(\Sigma_{\hat{\boldsymbol{Z}}_s})$, estimated from the data, then the entries of the adjusted prior matrix $\tilde{\Sigma}_{\boldsymbol{Z}_s}$ follow immediately once we have set the $m$ prior parameters $\left(W_{k_i} : i = 1,2,\dots,m\right)$ in $W_s$ only. Hence, computing a JABF for all causal configurations requires setting $mQ$ prior variance components only, which can be tuned to values used in previous studies[1,3], avoiding setting $O(m^2Q^2)$ potentially irrelevant covariance pairs. The assumption that the prior correlation matrix $\boldsymbol{\rho}(\Sigma_{\boldsymbol{Z}_s})$ is equal to the observed correlation matrix of the marginal *Z*-scores follows from the assumption that the estimated marginal effects correlation matrix and the prior marginal effects correlation matrix are generated from the same linkage disequilibrium (LD) pattern within each study.

We now discuss different choices for the empirical correlation matrix $\boldsymbol{\rho}(\Sigma_{\hat{\boldsymbol{Z}}_s})$ estimated from the data.

<u>Choices for the between-study empirical correlation matrix</u>

Correlation between $Z$-scores from two separate studies can arise when both studies share participants, i.e. there are overlapping samples. We refer to this generally as 'trait correlation', as it describes the origins of correlation in both the prior and observed correlation matrices. To approximate the correlation between any two $Z$-scores $Z_{j_i}$ and $Z_{k_l}$, where $i$ and $l$ denote the $i^{\text{th}}$ and $l^{\text{th}}$ traits, and $j$ and $k$ denote the $j^{\text{th}}$ and $k^{\text{th}}$ variants, we follow the approach of [3]. We assume that both traits have been standardised to have mean zero and unit variance, and both variants are centred at zero. Then, for two continuous traits, or one or two binary traits (assuming the genetic effect on the binary outcome is 'small', i.e. that a causal SNP accounts for a small proportion of trait variation), it follows that

$$\rho(Z_{j_i}, Z_{k_l}) = E\left[\frac{\sum_{n=1}^{N_i} y_{in} g_{inj}}{\hat{\sigma}_i \sqrt{\sum_{n=1}^{N_i} g_{inj}^2}} \frac{\sum_{n=1}^{N_k} y_{ln} g_{lnk}}{\hat{\sigma}_l \sqrt{\sum_{n=1}^{N_k} g_{lnk}^2}}\right] \approx \frac{N}{N_i N_k} \rho_{il} \hat{\rho}_{jk},$$

where $N$ denotes the number of overlapping samples (i.e. common individuals) between studies $i$ and $l$, $\rho_{il}$ denotes the correlation between traits $i$ and $l$ and $\hat{\rho}_{jk}$ is the squared correlation based on genotypic allele counts. Note we have assumed that $\hat{\sigma}_i \approx 1$ and $\hat{\sigma}_l \approx 1$, which follows on assuming that a causal SNP accounts for only a small proportion of trait variation. The parameter $\hat{\rho}_{jk}$ is approximated using the LD $r_{jk}^2$ between genotypes, which is estimated from haplotype frequencies. These parameters are not identical, but typically very similar. We estimate non-zero trait correlation, e.g. $\rho_{il} \neq 0$, in one of two ways, which we now discuss.

<u>Estimation of $\rho_{il}$</u>

The between-study covariance structure of the estimated $Z$-scores is unknown and is estimated in one of two ways:

(i). Estimating the Pearson's correlation between any two $Z$-score pairs[3]

(ii). Estimating the between study *tetrachoric correlations* of $Z$-score pairs[4]

Under the biologically plausible assumption that most genetic effects are drawn under the null-hypothesis of no association with a trait, the tetrachoric correlation approach[4] accounts for the possibility that a false positive in one study might reflect a false-positive in the corresponding results from other correlated studies. In doing so, the tetrachoric correlation approach avoids inflating the Type I error whilst also reducing the impact of correlations between true positives, allowing evidence to accumulate in favour of the alternative. This contrasts with correlating the $Z$-scores using Pearson's correlation coefficient which is conservative against the alternative, thus reducing power[3].

Tetrachoric correlation is computed by correlating the sign (direction) of the $Z$-scores, between studies, and therefore requires both the estimated effect size of the genotype $j$ on trait $i$, $\hat{\beta}_{j_i}$, and associated standard error, $\hat{V}_{j_i}$. For samples in which these data are available, our *HyPrColoc* software approximates the covariance parameters in $\Sigma_{\hat{\mathbf{Z}}_S}$ using the estimated tetrachoric correlations between traits by default.

#### S2.2. Causal configuration priors

The previous section addressed the challenge of evaluating the Bayes factor terms that appear in Equation (2). We now consider how the causal configuration priors, $P(S)$, that appear in Equation (2) may be specified. We consider two different specifications of $P(S)$: (1) Variant specific (VS) configuration priors; and (2) Conditionally uniform (CU) configuration priors.

***(1) Variant specific (VS) configuration priors***

For each SNP (i.e. for each column of the configuration matrix $S$), there are many different causal configurations. Each such configuration describes whether or not the SNP is causal for a particular combination of the *m* traits, and may be expressed as a binary column vector of length *m*. For example, the SNP might not be associated with any of the *m* traits (corresponding to a column vector of zeros), or the SNP might be associated with trait 1 only (corresponding to a binary vector whose first element is 1, and whose other elements are 0), …, and so on. Since there are $2^m$ binary vectors of length *m*, it follows that there are $2^m$ different causal configurations. We wish to specify the prior probability associated with each of these.

To do this, we first introduce some notation: let $p_0$ denote the prior probability that the SNP is not associated with any of the $m$ traits; let $p_1$ denote the prior probability that the SNP is associated with trait one only; let $p_2$ denote the prior probability that the SNP is associated with trait two only; ...; let $p_{j_1 j_2 \dots j_k}$ denote the prior probability that the SNP is associated with the $k$ traits $\{j_1, j_2, \dots, j_k\}$ (and no others); …; and let $p_{12\dots m}$ denote the prior probability that the SNP is associated with all traits. All such probabilities must clearly sum to 1, i.e.

$$p_0 + \sum_{k=1}^{m} \left( \sum_{j_1=1}^{m} \sum_{j_2 > j_1} \cdots \sum_{j_k > j_{k-1}} p_{j_1 j_2 \dots j_k} \right) = 1.$$

To avoid having to specify all of these prior probabilities separately (i.e. to avoid having to specify each of $p_0$, $p_1$, …, $p_{j_1 j_2 \dots j_k}$,…, in turn), we instead define the prior probabilities as follows,

$$p_{j_1 j_2 \dots j_k} = \begin{cases} p, & k = 1, \\ p \prod_{i=2}^{k} \left(1 - \gamma^{i-1}\right), & k \geq 2, \end{cases}$$

( 4 )

where $j_\ell \in \{1, \dots, m\}$, and $p$ and $\gamma$ are parameters that must be specified. We note that this formulation also implicitly fixes the value of $p_0$, since

$$p_0 = 1 - \sum_{k=1}^{m}\left(\sum_{j_1=1}^{m}\sum_{j_2>j_1}\cdots\sum_{j_k>j_{k-1}} p_{j_1 j_2 \ldots j_k}\right).$$

Equation (4) provides a general formula for specifying the prior on each column of the configuration matrix $S$, using just two parameters: $p$ and $\gamma$. The parameter $p$ is the probability that the SNP is associated with trait $\ell$ only (for any $\ell \in \{1, \ldots, m\}$). The role of the parameter $\gamma \in [0,1]$ is revealed by considering the conditional prior probability of an additional association as follows. Let $T_k$ denote trait $k$ and $T_{1,2,\ldots(k-1)}$ denote the collection of traits 1 to $k-1$, then

$$P(\text{SNP is causal for } T_k \mid \text{SNP is causal for } \mathrm{T}_{1,2\ldots(k-1)})$$

$$= \frac{\sum_{j=0}^{m-k}\left(\binom{m-k}{j}\prod_{i=1}^{k+j}(1-\gamma^{i-1})\right)}{\sum_{j=0}^{m-k+1}\left(\binom{m-k+1}{j}\prod_{i=1}^{k+j-1}(1-\gamma^{i-1})\right)}$$

$$\approx 1 - \gamma^{k-1}.$$

Hence, $\gamma$ controls the prior probability of an additional association given that the SNP is associated with at least one trait. Notably, large values of $\gamma$ make an additional colocalization unlikely whereas small values make additional colocalizations likely. The properties of Equation (4) can be summarised as follows:

i. All possible SNP-trait prior probabilities are specified using two easily interpretable parameters: $p$ and $\gamma$.
ii. Recovers the approach of [1] when $m = 2$, $p = 10^{-4}$ and $\gamma = 0.99$, i.e.

$$P(\text{SNP causal for } T_2 \mid \text{causal for } T_1) = \frac{1-\gamma}{1+(1-\gamma)}$$

$$\approx 1 - \gamma$$

$$= 0.01.$$

That is, of all SNPs associated with one trait, 1 in 100 of them will be associated with another trait.

iii. Allows evidence to grow in favour of $k$ traits co-localising conditional on evidence supporting $k-1$ traits co-localising, i.e.

$$P(\text{SNP causal } T_k \mid \text{causal } T_{1,2,\ldots(k-1)}) \geq P(\text{SNP causal } T_{k-1} \mid \text{causal } T_{1,2,\ldots(k-2)}).$$

iv. The limiting behaviour of an additional colocalization given that $m-1$ traits colocalize at a given SNP is given by

$$\lim_{m\to\infty} P(\text{SNP causal } T_m \mid \text{causal } T_{1,2,\ldots(m-1)}) = \lim_{m\to\infty} \frac{1-\gamma^{m-1}}{1+(1-\gamma^{m-1})} = 0.5.$$

That is, of all SNPs associated with $m-1$ traits, half will be associated with all $m$ traits as $m \to \infty$. Thus we are agnostic about whether a SNP co-localizing with $m-1$ traits also colocalizes with an additional trait.

So far, we have just considered specification of the prior for a single column of the configuration matrix, *S*, i.e. for just a single SNP. In principle, we could specify SNP-specific values for $p$ and $\gamma$, which might depend on, say, MAF or function of the SNP. Here, we assume a common value of $p$ and $\gamma$ for all SNPs.

***(2) Conditionally uniform (CU) configuration priors***

In the absence of (SNP) level prior information, a typical strategy is to assume uniform priors; i.e. that all matrices *S* are equally probable *a priori*. However, this would mean that hypotheses comprising larger numbers of configurations would be more probable *a priori*. To avoid issues with massive differences in the cardinality of the configuration space between hypotheses, we introduce the CU prior, which ensures that all non-null hypotheses are equally probable *a priori*. That is, we want the following property: $P(H_i) = P(H_j)$ for all non-null hypotheses $H_i$ and $H_j$. We do this by specifying the prior probability of *S* as follows

$$P(S|S \in \mathcal{S}_H) \propto {}^{1}/_{|\mathcal{S}_H|}.$$

We set the constant of proportionality by additionally requiring that $P(H_j) = \varepsilon P(H_0)$, and recalling that the prior probabilities must sum to 1. Throughout, we set $\varepsilon = 10^{-4}$ (used similarly to $p_1$ in [1]).

Under the CU prior, the configuration prior probabilities depend on the cardinality of the hypothesis configuration spaces. This means that the set of hypotheses $\mathcal{H}_{(k)}$ (which assume a collection of $k \geq 1$ traits share a CV all remaining $m - k$ traits do not have a CV in the region) have associated causal configuration sets $\mathcal{S}_{\mathcal{H}_{(k)}}$ with the same cardinality $Q$ and therefore have the same CU prior probability. Hence, these priors vary by the size of the region $Q$ only. In contrast, the set of hypotheses $\mathcal{H}_{(m-1,1)}$ have a causal configuration set with cardinality $\left|\mathcal{S}_{\mathcal{H}_{(m-1,1)}}\right| = mQ(Q-1)$, which varies in both the number of traits and the size of the genomic region.

Two benefits of the CU causal configuration priors are:

- Prior specification depends only on a single unknown parameter $\varepsilon$, which here we set to $10^{-4}$.
- It automatically adapts to the ‘scale’ of the problem, i.e. number of traits $m$ and size of the genomic region $Q$.

An observed drawback, from our simulations, of the CU prior is that there is a small increase in the false-positive rate (i.e. falsely detecting colocalization) relative to the VS prior which does not suffer from these issues (**Figure S3**).

#### S2.3. Hypothesis prioritisation: the *HyPrColoc* approximation

We now complete the exposition of our Bayesian model selection approach by considering how the denominator on the right hand side of Equation (2), i.e. $\sum_{H^* \in \Omega} \sum_{S \in \mathcal{S}_{H^*}} BF(S) \frac{P(S)}{P(S_0)}$, may be efficiently evaluated.

##### Hypothesis space

We use the notation $\mathcal{H}_{(i,j,\ldots)}$ to denote a *set* of hypotheses in which a collection of $i$ traits share a causal variant, a separate collection of $j$ traits share a distinct causal variant, and so on (**Main text Figure 1**). For example, $\mathcal{H}_{(2,1)}$ denotes the set of hypotheses in which 2 traits share a causal variant, a single trait has a distinct causal variant and all remaining $m-3$ traits do not have a causal variant in the region. A single hypothesis is denoted by $H$, e.g. $H_m$ denotes the hypothesis in which all $m$ traits colocalize. Assuming at most one causal variant for each trait these data generating hypotheses can be combined to generate a hypothesis space ($\Omega$). The cardinality, or size, of the set containing all data generating hypotheses for $m$ traits follows

$$|\Omega| = \sum_{k=0}^{m} \binom{m}{k} Bell(k) = Bell(m+1),$$

where $Bell(.)$ denotes the Bell number, which in combinatorial mathematics represents the number of partitions of a set. This identity can be proved via induction using the intermediate identity above. Heuristically, however, on noting that $\binom{m}{k}$ is the number of ways of choosing $k$ traits from a possible $m$ traits and $Bell(k)$ is the number of ways that the $k$ traits can be partitioned into shared and distinct causal mechanisms, the identity follows naturally. We show later that we need only compute contributions from the sets of hypotheses in $(\mathcal{H}_{(m-1)} \cup H_m)$ and $\mathcal{H}_{(m-1,1)}$ to approximate the posterior probability that all traits colocalize. The cardinalities of these sets are

$$\left|\mathcal{H}_{(m-1)} \cup H_m\right| = m + 1,$$

$$\left|\mathcal{H}_{(m-1,1)}\right| = m.$$

Note that,

$$\lim_{m\to\infty} \frac{2m+1}{Bell(m+1)} = 0.$$

That is, for large $m$ the sets $(\mathcal{H}_{(m-1)} \cup H_m)$ and $\mathcal{H}_{(m-1,1)}$ are extremely small relative to $\Omega$. For example, suppose there are $m = 3$ traits under consideration, then there are 15 mutually exclusive hypotheses of which 7 are contained within the sets $(\mathcal{H}_{(m-1)} \cup H_m)$ and $\mathcal{H}_{(m-1,1)}$, accounting for around 47% of all hypotheses. However, when $m = 20$ there are approximately $4.74 * 10^{14}$ mutually exclusive hypotheses and the collections $(\mathcal{H}_{(m-1)} \cup H_m)$ and $\mathcal{H}_{(m-1,1)}$ account for around $0.0000000000086\%$ of all hypotheses. Therefore almost $100\%$ of the data generating hypotheses are contained within the set $\Omega \setminus \{ \mathcal{H}_{(m-1)} \cup H_m \cup \mathcal{H}_{(m-1,1)} \}$.

As noted from Figure 4 in the main text, a method (e.g. MOLOC[5]) which aims to compute contributions from all hypotheses becomes computationally impractical beyond $m > 4$ traits, highlighting the significant benefits of the hypothesis prioritisation approach. Our major result, HyPrColoc, is that the posterior probability of full co-localization (PPFC) across $m$ traits, $P(H_m|D)$, can be closely approximated for a general number of traits $m$ and SNPs $Q$ by computing the denominator in Equation (2), from the sets of hypotheses $(\mathcal{H}_{(m-1)} \cup H_m)$ and $\mathcal{H}_{(m-1,1)}$ only.

#### Posterior probability of colocalization

Let $P_{all}$, $P_{scv}$, $P_m$, $P_{dcv}$ and $P_{(m-1,1)}$ be given by

$$P_{all} = \frac{P(\Omega|D)}{P(H_0|D)},$$

$$P_{scv} = \frac{P(H_0|D) + \sum_{k=1}^{m-1} P(\mathcal{H}_k|D) + P(H_m|D)}{P(H_0|D)},$$

$$P_{dcv} = P_{all} - P_{scv},$$

$$P_m = \frac{P(H_m|D)}{P(H_0|D)},$$

and

$$P_{(m-1,1)} = \frac{P(\mathcal{H}_{(m-1,1)}|D)}{P(H_0|D)}.$$

Note that $P_{scv}$ is composed from the set of all subsets of hypotheses which assume that a collection of $k \geq 1$ traits have and share a causal variant (*SCV*), the remaining $m - k$ traits do not have a CV in the region. $P_{dcv}$ is composed from sets of hypotheses which assume two or more traits have distinct causal variants (*DCVs*).

We use these statistics to introduce two rapidly computable values that quantify the probability that two criteria necessary for colocalization are satisfied. The first of these criteria is that all the traits must share an association with one or more variants within the region. $P_R$, which we refer to as the *regional association probability*, is the probability that this criterion is satisfied:

$$P_R = \frac{P_m}{P_{scv}} > \frac{P_m}{P_{all}} = P(H_m|D) \qquad (= PPFC)$$

*( 5 )*

$P_R$ is greater than or equal to the PPFC of all traits being colocalized, thus, a small value for $P_R$ provides evidence against the PPFC. Large values, however, implicate all traits sharing one or more genetic predictors in the genomic region. For each trait the predictor might: (i) be a causal variant or (ii) appear owing to LD with a causal variant. To distinguish between evidence supporting a single shared CV across all traits, from multiple shared causal variants or LD between distinct causal variants, we have a second criterion that ensures the shared associations between all traits are owing to a single shared putative causal variant,

$$P_A = \frac{P_m}{P_m + P_{(m-1,1)}} > \frac{P_m}{P_{all}} = P(H_m|D).$$

( 6 )

We call $P_A$ the *alignment* probability as it differentiates between all traits having a single shared CV (i.e. alignment of the CV for each trait) and an alternative in which a subset of $m-1$ traits share a CV with the remaining trait has a distinct CV elsewhere in the region. Following the same approach as in the main text **Methods**, the regional and alignment probabilities can be used to accurately and efficiently assess evidence of a full colocalization hypothesis:

$$P(H_m|D) = \frac{P_m}{P_{scv} + P_{dcv}} = \frac{P_m}{P_{scv}} \frac{P_{scv}}{P_{scv} + P_{dcv}}$$

$$= P_R \frac{P_m}{\left(P_m + P_{(m-1,1)}\right) - \left((1 - P_R)P_{(m-1,1)} - P_R P^c_{(m-1,1)}\right)}$$

$$= \frac{P_R P_A}{1 - \left((1 - P_R)(1 - P_A) - P_R(1 - P_A)\frac{P^c_{(m-1,1)}}{P_{(m-1,1)}}\right)}$$

$$= P_R P_A + \mathcal{O}(\delta_A^2 + \delta_R \delta_A), \qquad \delta_R, \delta_A \longrightarrow 0,$$

( 7 )

where,

$$\delta_R = 1 - P_R,$$

$$\delta_A = 1 - P_A$$

and the *deprioritised hypotheses* in $P^c_{(m-1,1)}$ are shown (see **S4.4 Properties of the HyPrColoc approximation**) to satisfy

$$\frac{P^c_{(m-1,1)}}{P_{(m-1,1)}} = \mathcal{O}(\delta_A + \delta_R).$$

*( 8 )*

The *HyPrColoc approximation* is given by

$$P_{hypr}(H_m|D) = P_R P_A > \big(P(H_m|D)\big)^2,$$

which is bounded below by the square of the PPFC. Moreover, our result shows that

$$P(H_m|D) \to 1 \quad \Leftrightarrow \quad P_R \to 1 \ \text{ and } \ P_A \to 1,$$

i.e. that is necessary and sufficient to identify that the regional and alignment statistics tend to unity in order to know that the PPFC tends to unity. We show later that $P_R$ satisfies

$$P_R = \frac{P_m}{1 + P_m + P_{(m-1)}}\big(1 + \mathcal{O}(\delta_R^2)\big), \quad \delta_R \to 0.$$

Consequently, $P_R$ can be accurately computed using $m + 1$ hypotheses ($\mathcal{H}_{(m-1)} \cup H_m$), each with $Q$ causal configurations. Thus $P_R$ has $\mathcal{O}(mQ)$ computational complexity. $P_A$ is computed from an additional $m$ hypotheses ($\mathcal{H}_{(m-1,1)}$) each with $Q(Q-1)$ causal configurations, thus has $\mathcal{O}(mQ^2)$ computational complexity. In total therefore, the PPFC can be accurately approximated by computing only $\mathcal{O}(mQ^2)$ causal configurations. This contrast with the computationally prohibitive process of enumerating all $(Q+1)^m$ causal configurations.

We therefore obtain an approximation to the PPFC in two stages:

1. Assessment of the probability that there exists shared association *region* across all traits via $P_R$.

2. Assessment of the probability that the genetic signals *align* at a causal variant shared across all traits via $P_A$.

### S3. The *HyPrColoc* Bayesian divisive clustering algorithm

The general goal of a colocalization analysis is to identify patterns of colocalization amongst a large list of traits. *HyPrColoc* does this using a *branch and bound* model-based Bayesian divisive clustering algorithm.

**Algorithm design**

Let $\boldsymbol{T}_{j_1 j_2 \dots j_k}$ denote a specific collection of $k$ traits. We define the regional probability for the collection $\boldsymbol{T}_{j_1 j_2 \dots j_k}$ to be

$$P_R\left(\boldsymbol{T}_{j_1 j_2 \dots j_k}\right) = \frac{P_k}{1 + P_k + P_{(k-1)}}$$

$$= \frac{P\left(D \middle| H_{\boldsymbol{T}_{j_1 j_2 \dots j_k}}\right) p_{j_1 j_2 \dots j_k}}{P(D|H_0)p_0 + P\left(D \middle| H_{\boldsymbol{T}_{j_1 j_2 \dots j_k}}\right) p_{j_1 j_2 \dots j_k} + \sum_{l=1}^{k} P\left(D \middle| H_{\boldsymbol{T}_{j_1 j_2 \dots j_{(l-1)} j_{(l+1)} \dots j_k}}\right) p_{j_1 j_2 \dots j_{(l-1)} j_{(l+1)} \dots j_k}}$$

and the alignment probability to be

$$P_A\left(\boldsymbol{T}_{j_1 j_2 \dots j_k}\right) = \frac{P_k}{P_k + P_{(k-1,1)}}$$

$$= \frac{P\left(D \middle| H_{\boldsymbol{T}_{j_1 j_2 \dots j_k}}\right) p_{j_1 j_2 \dots j_k}}{P\left(D \middle| H_{\boldsymbol{T}_{j_1 j_2 \dots j_k}}\right) p_{j_1 j_2 \dots j_k} + \sum_{l=1}^{k} P\left(D \middle| H_{\left(\boldsymbol{T}_{j_1 j_2 \dots j_{(l-1)} j_{(l+1)} \dots j_k}, T_{j_l}\right)}\right) p_{j_l} p_{j_1 j_2 \dots j_{(l-1)} j_{(l+1)} \dots j_k}}$$

where $H_{\boldsymbol{T}_{j_1 j_2 \dots j_k}}$ denotes the hypothesis that traits $\boldsymbol{T}_{j_1 j_2 \dots j_k}$ share a causal variant and $H_{\left(\boldsymbol{T}_{j_1 j_2 \dots j_{(l-1)} j_{(l+1)} \dots j_k}, T_{j_l}\right)}$ denotes the hypothesis that the collection of traits $\boldsymbol{T}_{j_1 j_2 \dots j_{(l-1)} j_{(l+1)} \dots j_k}$ share a causal variant and the trait $T_{j_l}$ has a distinct causal variant. In addition, we let $K = \{\}$ be an empty set used to store the set of colocalized and $L = \{\}$ be an empty set used to store the traits which do not colocalize with any other trait.

Stage 1 of the BB algorithm aims to identify the cluster of traits $\boldsymbol{T}_{j_1 j_2 \dots j_k}$ with the strongest evidence of a regional association from the set of all traits $\boldsymbol{T}_{12\dots m}$. The cluster $\boldsymbol{T}_{j_1 j_2 \dots j_k}$ is then assessed for alignment at a single causal variant. If all traits in the cluster do not align at a single causal variant, then the cluster is partitioned into two clusters: one cluster contains the trait least likely to share a causal variant the other cluster contains the remaining traits. This process is described by 4 main steps:

1. Computation of the regional association probability for all traits $\boldsymbol{T}_{12\dots m}$ which involves computing $m+1$ probabilities for each of the hypotheses considered in $P_R(\boldsymbol{T}_{12\dots m})$. If $P_R(\boldsymbol{T}_{12\dots m}) \geq P_{\mathrm{R}}^{*}$, where $P_{\mathrm{R}}^{*}$ denotes a user defined probability with which we accept evidence of a regional association across the traits, then $\boldsymbol{T}_{12\dots m}$ is a candidate cluster of colocalized traits and we move to step 4.
2. If $P_R(\boldsymbol{T}_{12\dots m}) < P_{\mathrm{R}}^{*}$ dismiss the possibility of all $m$ traits colocalizing and partition the cluster $\boldsymbol{T}_{12\dots m}$ by either:
   a) removing trait $k^*$ that is least likely to share one or more genetic predictors with all other traits,

$$\max_{l \in \{1,2,\dots,k\}} P\left(D \middle| H_{\boldsymbol{T}_{12\dots(l-1)(l+1)\dots m}}\right) = P\left(D \middle| H_{\boldsymbol{T}_{12\dots(k^*-1)(k^*+1)\dots m}}\right).$$

   This process has $\mathcal{O}(mQ)$ cost.

   b) removing trait $k^*$ that is least likely to align at a SCV with the other $m-1$ traits, i.e.

$$\max_{l \in \{1,2,\dots,k\}} P\left(D \middle| H_{(\boldsymbol{T}_{12\dots(l-1)(l+1)\dots m}, T_l)}\right) = P\left(D \middle| H_{\left(\boldsymbol{T}_{12\dots(k^*-1)(k^*+1)\dots m}, T_{k^*}\right)}\right).$$

   This process has $\mathcal{O}(mQ^2)$ cost.

3. Repeat steps 1-2 until a collection of traits $\boldsymbol{T}_{j_1 j_2 \dots j_k}$ is identified which satisfies

$$P_R\left(\boldsymbol{T}_{j_1j_2\dots j_k}\right) \geq P_{\mathrm{R}}^*.$$

The default *HyPrColoc* BB cluster algorithm uses the more computationally efficient 2(a) to reduce the traits down to $\boldsymbol{T}_{j_1j_2\dots j_k}$

4. Assess colocalization of the traits $\boldsymbol{T}_{j_1j_2\dots j_k}$, *i.e.* compute

$$P_R\left(\boldsymbol{T}_{j_1j_2\dots j_k}\right)P_A\left(\boldsymbol{T}_{j_1j_2\dots j_k}\right).$$

a) If $P_R\left(\boldsymbol{T}_{j_1j_2\dots j_k}\right)P_A\left(\boldsymbol{T}_{j_1j_2\dots j_k}\right) \geq P_R^*P_A^*$ then $\boldsymbol{T}_{j_1j_2\dots j_k}$ are identified as being colocalized. We set

$$K \to K \cup \boldsymbol{T}_{j_1j_2\dots j_k}$$

remove $\boldsymbol{T}_{j_1j_2\dots j_k}$ from the sample and repeat steps 1 - 4 using $\boldsymbol{T}_{12\dots m}\backslash\{K, L\}$.

b) If $P_A\left(\boldsymbol{T}_{j_1j_2\dots j_k}\right) < P_A^*$ and $k > 2$ we partition the cluster $\boldsymbol{T}_{j_1j_2\dots j_k}$ by removing the trait least likely to align at a single causal variant using 2(b) and repeat 1 - 3.

c) If $P_A\left(\boldsymbol{T}_{j_1j_2\dots j_k}\right) < P_A^*$ and $k = 2$ we identify the trait $T^* \in \{T_{j_1}, T_{j_2}\}$ least likely to align at a SCV using 2(b). $T^*$ is deemed to not colocalize with any other trait, we

$$L \to L \cup T^*$$

and remove $T^*$ from the sample. Steps 1 – 3 are repeated using traits $\boldsymbol{T}_{12\dots m}\backslash\{K, L\}$.

The whole process is repeated until

$$K \cup L = \boldsymbol{T}_{12\dots m}.$$

**Figure S1** illustrates the regional and alignment branch selection algorithm at phase 2. **Figure 3** displays the BB algorithm pipeline to assess collections of colocalized traits from all $m$ traits.

### S4. Additional simulation results and further mathematical details

#### S4.1. Additional simulation results

The simulation strategy is the same as defined in the main text and we summarise this again here. To create genomic loci with realistic patterns of LD, for each simulation scenario we simulated 1,000 datasets by resampling phased haplotypes from the European samples in 1000 Genomes[6] and for each dataset we randomly selected one of the first 50 regions confirmed to be associated with CHD[7]. Unless stated otherwise, for traits that have a CV in the region the variant explains 1% of trait variance and all studies considered have a sample size of $N = 10{,}000$.

**Reworking the BB clustering algorithm assessment with a reduced number of traits**

To assess the performance of HyPrColoc for varying numbers of traits, we repeated the simulation scenario described in the main text, but with a reduced number of traits. Instead of 100 traits, in the present assessment we simulated 20 traits from non-overlapping datasets with 10,000 individuals under three situations: in all scenarios there exists a cluster of 10 traits sharing a single causal variant and either the remaining 10 traits (i) do not have a causal variant; (ii) form a separate cluster of 10 traits sharing a distinct causal variant or; (iii) separately have distinct causal variants. In all scenarios, the causal variant for each trait explained 1% of trait variance and the probability parameters were set to $P_R^* = P_A^* = 0.6$. Results are stratified by posterior probability in scenario (i) and by LD between causal variants in scenarios (ii) and (iii). In scenario (i), the probability of detecting the true cluster of colocalized traits was greater than 0.9 and, when considering only results where $P_R P_A > 0.7$, this probability increased to greater than 0.98 (**Figure S2**). When a second cluster of colocalized traits were added, scenario (ii), the detection probability decreased by only $\approx 0.01$, however this was owing to strong LD between the causal variants, i.e. $r^2 > 0.95$. Considering only situations in which $r^2 < 0.95$ between the two causal variants returns similar performance to scenario (i) (**Figure S2**). Finally, when all the remaining traits had a distinct causal variant, scenario (iii), the detection probability dropped markedly ($\approx$ $0.15$) owing to the increased chance that the causal variant from a non-colocalized trait is in strong LD with the colocalized causal variant, i.e. $r^2 > 0.95$, a scenario in which no algorithm is likely to perform well. In scenarios where $r^2 \leq 0.95$, the detection probability was found to be ≈ 0.95.

In summary, on reducing the number of traits from 100 (in the main text) to 20 the overall performance of the algorithm is improved, i.e. approximately a 5% increase in true detection rates over all three scenarios. These results highlight some sensitivity in the performance of the algorithm to the number of traits under consideration.

#### Comparison of conditionally uniform and variant level prior frameworks

Here we compare the conditionally uniform (CU), e.g. [3], and variant specific (VS) configuration prior frameworks when either all traits share a causal variant or a subset of traits do.

When all traits share a causal variant, we simulated $m = 2, 5, 10$ quantitative and independent traits in which the CV explains $0.5\%, 1\%$ and $2\%$ of trait variance, and across study sample sizes $N = 5{,}000, 10{,}000, 20{,}000$ (**Figure S3**). The results indicate that in all scenarios the CU causal configuration prior has markedly better distribution properties, i.e. a tighter distribution between the first, fifth (median), and ninth deciles, and a higher overall median posterior probability of true colocalization, relative to the VS configuration priors.

When a subset of traits colocalize, the CU configuration prior *can* lead to an increased false positive rate. To highlight this, we simulated 10 traits from non-overlapping studies (we deal with overlapping participants and correlated traits shortly) whereby 7 of 10 traits share a CV and the remining 3 traits do not have a CV in the region (**Figure S4)**. Generally speaking, the BB algorithm performs well under both configuration prior scenarios, regularly identifying the true cluster of colocalized traits, i.e. $> 0.9$ true detection probability in most scenarios. However, the detection rate can increase or decrease depending on the regional and alignment thresholds and we note that the optimal values vary depending on the type of configuration prior employed (**Figure S4**). For example, the true detection probability is $> 0.95$ when setting $P_R^* = P_A^* = 0.5$ for the VS prior and $P_R^* = P_A^* = 0.7$ for the CU prior (**Table S3**). This impacts on the false positive and negative detection rate as well. Across all values of the regional and alignment thresholds considered, the BB algorithm using the CU configuration prior identified a cluster of ≥ 8 traits sharing a CV in 3% of cases, when $P_R^* = P_A^* = 0.7$, and in 8% of cases when $P_R^* = P_A^* = 0.5$. The cluster of 7 truly colocalized traits were always included in these collections. The false positive rate using the VS configuration prior was nominal across all threshold values however, instead for the number of traits considered the VS prior appears conservative, e.g. the cluster of traits identified as sharing a CV under the VS prior has a false negative rate of 10% when $P_R^* = P_A^*$=0.7, this reduced to $<$ 4% when setting $P_R^* = P_A^*$=0.5.

To attenuate sensitivity to the choice of configuration prior, we can change the regional and alignment threshold parameters to suit the choice of configuration prior used. For example, when deducing that a cluster of traits are colocalized using the CU prior we can set $P_R^* = P_A^* \geq 0.7$, whereas under the VS prior these values can be reduced $P_R^* = P_A^* \geq 0.5$. This might help balance the performance of the BB algorithm between the different choices of configuration prior (**Figure S4** and **Table S3)**. In general, increasing the regional and alignment thresholds will reduce the probability of a cluster including one or more false positives under both priors.

#### Analysing traits from studies with overlapping samples

When two or more studies have common participants, *i.e.* overlapping participants (sometimes referred to as overlapping datasets), and there exists correlation between traits measured in each study, between study-level summary effects are correlated and we may wish to account for these correlations in our analysis: under the assumption that doing so might control the false positive detection rate. Here we discuss the impact of adjusting analyses to account for this. Our discussion focuses on assessing three questions:

(i) If traits are *known* a priori to be correlated, should the prior probability of a causal configuration account for trait correlation?
(We demonstrate that the answer to this can be yes, but a general strategy to account for known trait correlation at the level of the prior configuration probabilities is complex. Hence, we aim to assess the following as well)

(ii) What are the consequences of ignoring known trait correlation at the level of the prior configurations but adjusting analyses to account for (observed) correlation of the summary effects (via the likelihood terms) when computing the posterior probability of colocalization?

(iii) What are the consequences of ignoring known trait correlation at the level of prior configurations as well as ignoring (observed) correlation of summary effects when computing the posterior probability of colocalization, i.e. incorrectly assuming independence between studies (non-overlapping samples)?

**Trait correlation and prior configuration probabilities**

We define a symmetric genetic mechanism between two or more traits as either a colocalization configuration or a configuration in which each trait has a distinct causal variant and these variants are in strong LD with one another. We show that configurations that reflect symmetric genetic mechanisms are *a priori* more probable than asymmetric configurations as the magnitude of correlation between the two traits increases. We illustrate our findings via a simple, but widely used, model. Suppose traits $T_1$ and $T_2$ are given by the linear models

$$T_1 = \beta_1 g_1 + \epsilon_1,$$

$$T_2 = \beta_2 g_2 + \epsilon_2,$$

where the genetic information is assumed mean zero and independent of the residual information, i.e.

$$g_i \perp \epsilon_j, \quad i = 1,2 \text{ and } j = 1,2,$$

and residual terms are mean zero random variables. Let $\rho_{12} = cor(T_1, T_2)$ denote the *marginal* Pearson's correlation between the traits, i.e. taking both $g_i$ and $\epsilon_i$ as random. As the magnitude of correlation between the traits increases, the traits can increasingly be expressed as a linear multiple of one another, e.g.

$$\lim_{\rho_{12} \to \pm 1} T_2 = \kappa T_1,$$

for some constant $\kappa$. If the traits are standardised to have zero mean and unit variance, then $\kappa = \pm 1$ and, owing to exogeneity between the genetic and residual information, matching the genetic terms reveals that

$$g_2 = \pm \frac{\beta_1}{\beta_2} g_1$$

and consequently

$$T_2 = \pm \beta_1 g_1 \pm \epsilon_1 \quad : \quad \rho_{12} \to \pm 1.$$

Thus, as $\rho_{12} \to \pm 1$ the causal variant for trait 1 is *also* the causal variant for trait 2. In terms of prior configuration probabilities, therefore, if two quantitative traits are *known* to be correlated then it follows that

$$\lim_{\rho_{12} \to \pm 1} \begin{cases} P\big(g_j \text{ causal } T_2 \mid g_i \text{ causal } T_1\big) = 1 \quad : \quad j = i & \text{(colocalization prior)} \\ P\big(g_j \text{ causal } T_2 \mid g_i \text{ causal } T_1\big) = 0 \quad : \quad j \neq i \text{ and } g_j \perp g_i. & \text{(misalignement prior)} \end{cases}$$

The result generalises to discrete traits, e.g. a disease outcome, derived by linearizing the link function in Equation (1) under the assumption that a causal variant explains only a small amount of trait variance.

The key point is: known correlation between two or more traits can be incorporated into the prior configuration probabilities, upweighting a colocalization mechanism relative to a non-symmetric (i.e. un-correlated) mechanism. We show later that ignoring known trait correlation in the prior configuration probabilities can reduce power to detect a cluster of colocalized in colocalization analyses.

**Ignoring trait correlation in the likelihood terms**

Recall that $JABF(S)$ in Equation (3) approximates the (null-weighted) likelihood contribution to the posterior probability of causal configuration $S$ in Equation (2). We re-write $JABF(S)$ for clarity:

$$JABF(S) = \sqrt{\frac{|\Sigma_{\hat{\boldsymbol{Z}}_s}|}{|\Sigma_{\hat{\boldsymbol{Z}}_s} + \tilde{\Sigma}_{\boldsymbol{Z}_s}|}} \exp\left\{\frac{1}{2}\,\hat{\boldsymbol{Z}}^T\left(\Sigma_{\hat{\boldsymbol{Z}}_s} + \Sigma_{\hat{\boldsymbol{Z}}_s}\tilde{\Sigma}_{\boldsymbol{Z}_s}^{-1}\Sigma_{\hat{\boldsymbol{Z}}_s}\right)^{-1}\hat{\boldsymbol{Z}}\right\}.$$

Let $\Sigma'$ denote the argument of the exponential term, i.e.

$$\Sigma' = \Sigma_{\hat{\boldsymbol{Z}}_s} + \Sigma_{\hat{\boldsymbol{Z}}_s}\tilde{\Sigma}_{\boldsymbol{Z}_s}^{-1}\Sigma_{\hat{\boldsymbol{Z}}_s}.$$

and $\Sigma'_{mis}$ be the mis-specified analogue of $\Sigma'$. If misspecification of the correlation structure artificially increases a likelihood term, then the 'true' likelihood is less than the misspecified likelihood, i.e.

$$JABF(S) < JABF_{mis}(S).$$

Assuming that the dominant behaviour of $JABF(S)$ is owing to the exponential term, the above inequality can be approximated by

$$\widehat{\boldsymbol{Z}}^T \Sigma'^{-1} \widehat{\boldsymbol{Z}} < \widehat{\boldsymbol{Z}}^T \Sigma'^{-1}_{mis} \widehat{\boldsymbol{Z}}$$

$$\Rightarrow \quad \widehat{\boldsymbol{Z}}^T \left( \Sigma'^{-1} - \Sigma'^{-1}_{mis} \right) \widehat{\boldsymbol{Z}} < 0.$$

Hence, if the matrix $\left( \Sigma'^{-1} - \Sigma'^{-1}_{mis} \right)$, which we refer to as the *contrast* matrix, is strictly *negative definite* then accounting for true correlation between the summary effects can shrink the likelihood relative to misspecifying trait correlation. Consequently, the misspecified framework can lead to an inflated false positive rate. The opposite is also true, if the contrast matrix is positive definite then misspecification can lead to an increased false positive rate.

We use these properties to assess the performance of the BB algorithm when correlated traits are wrongly assumed independent, by varying the contrast matrix to be either negative or positive definite.

**Results**

We simulated 10 quantitative traits with correlated *residual error* terms, correlated under a compound symmetric correlation framework, and assuming *complete overlap* of participants between studies (i.e. participants are the same in all 10 studies). Note that marginal trait correlations are not necessarily compound symmetric owing to the presence of genetic correlation. To assess performance across a range of misspecification scenarios, the correlation parameter $\rho$ was set to either:

- $\rho \in \{0.5, 0.8\}$ (negative definite contrast matrix ~ increase false positive rate)
- $\rho = -0.1$ (positive definite contrast matrix ~ increase false negative rate)

In all scenarios, 7 of the 10 traits had a shared CV explaining 1% of trait variance, the remaining 3 traits had no CV in the region.

To illustrate the impact of accounting for trait correlation through configuration prior terms, we constructed correlation *aware* prior configuration probabilities as follows. Under the CU prior, we assume that asymmetric genetic configurations are $1-\rho^2$ less likely, i.e.

$$P\ (S_H) = \begin{cases} \epsilon/_Q, & H \in \mathcal{H}_{(i)} \cup H_m \ \text{(colocalization)} \\ (1-\rho^2)\left(\epsilon/_{mQ(Q-1)}\right), & H \in \mathcal{H}_{(m-1,1)} \ \text{(misalignment)} \end{cases}.$$

Under the VS prior, Equation (4), we let the probability of an additional colocalization be a function of the correlation, e.g. in the analysis of $m = 2$ traits

$$P(\text{SNP is causal for } T_2 \mid \text{SNP is causal for } \mathrm{T}_1) = 1-\gamma = \rho^2,$$

and, analogous to a CU prior, asymmetric configurations are multiplied by $1-\rho^2$.

**$\underline{\boldsymbol{Residual\ correlation}: \boldsymbol{\rho = 0.5}}$**

We analysed the data in three ways:

(i) Ignoring all correlation, i.e. assuming non-overlapping participants between pairs of studies and ignoring correlation in configuration priors (bottom **Figure S4** and **Table S2**)

(ii) Adjusting for correlation between the summary data in the computation of the likelihood (**Figure S5** and **Table** S2)

(iii) Adjusting for correlation between the summary data in the computation of the likelihood and adjusting for known trait correlation via correlated configuration priors (**Figure S5** and **Table S2**)

As anticipated, the correlation adjusted likelihood (scenario (ii)) is smaller than the likelihood computed assuming independence (scenario (i)) (**Table S2**). Consequently, adjusting for observed correlation but ignoring correlation at the level of the configuration prior shrinks the probability of correctly identifying all colocalized traits (scenarios (ii) vs (iii)). For example, the probability of correctly identifying the cluster of colocalized traits is: 0.03 when *only* adjusting for observed correlation; 0.93 when adjusting for both observed and *a priori* ('known') correlation and; 0.95 when ignoring observed and *a priori* correlation (all results derived using VS configuration priors) (**Table S2**). We note how similar performance is when either *ignoring* evidence of observed *and* a priori correlation or adjusting for them *both*. We also note a significant difference in the performance of the BB algorithm between the type of configuration prior assumed. This is because the CU framework places the same configuration prior probability for clusters of two or more colocalized traits whereas the VS configuration prior is smaller for larger clusters of colocalized traits.

We might anticipate that the independence assumption would inflate the false positive detection rate, when a colocalization likelihood is inflated (by assuming independence between study summary data) relative to a correlation adjusted likelihood. However, this does not appear to be the case: both the distribution of the posterior probability of colocalization and the true detection rate of the BB algorithm closely follows that when correctly assuming independence between the traits. For example, when we compare the results in which we have (a) wrongly assumed the study samples are non-overlapping and that there is no (a priori) trait correlation and (b) correctly assumed the study samples are non-overlapping and there is no (a priori) trait correlation, we find the false positive rate was 0.003 under (a) and 0.01 under (b) and the probability of correctly identifying all colocalized traits was 0.97 under (a) and 0.95 under (b) using the VS prior framework (**Table S2**). Using the CU prior, the results were similar. In summary, these results illustrate that when *correctly* or *wrongly* assuming between study samples are non-overlapping and that there is no (a priori) trait correlation the BB algorithm performs similar well. We show this conclusion holds across a wider range of scenarios.

We further assessed the performance of the BB algorithm across various strengths and directions of the correlation parameter $\rho$, i.e. when $\rho = -0.1$ and $\rho = 0.8$ and wrongly assuming non-overlapping samples between studies (**Table S3**). Notably, when $m = 10$ and $\rho = -0.1$, the contrast matrix $\left(\Sigma'^{-1} - \Sigma'^{-1}_{mis}\right)$ is positive definite and the correlation adjusted likelihood of colocalization is larger than when the correlation is ignored (results omitted). In this scenario we might anticipate that an independence assumption leads to a reduction in the true positive detection rate. However, once again, the distribution of the posterior probability of colocalization and the probability of correctly detecting the cluster of colocalized traits is maintained under independence assumption (**Table S**3).

#### Ignoring trait correlation: some remarks

Ignoring observed correlation of between study summary data, due to overlapping participants between studies, and ignoring *a priori* correlation at the level of the configuration priors has *two* main advantages when performing multi-trait colocalization analyses. Firstly, adjusting for observed trait correlation via the likelihood in Equation (2) requires repeatedly inverting potentially high-dimensional matrices in Equation (3). This process incurs a non-linear time cost which necessarily impacts on the computational efficiency of HyPrColoc (**Tables S2-S3**), making the analysis of hundreds of traits *realistic* only under the assumption of non-overlapping samples between studies. Secondly, in our investigation into the performance of the BB algorithm (**Tables S2-S3**) we conclude that analyses which adjust only for observed trait correlation have a significantly reduced probability of correctly detecting all colocalized traits. Adjusting for 'known' (*a priori*) correlation through the configuration priors can help to remedy this issue. However, a general approach to do this is not forthcoming and any approach will necessarily introduce a posterior sensitivity to both the reference LD information and the quality of the prior knowledge of trait correlation. This detail is avoided under an independence assumption.

Our investigation into these types of misspecification is brief and limited to compound symmetric correlation matrices relating the residuals in the outcome model. It is possible that our findings are consistent across a broader range of correlation structures. However, this property requires further investigation (but is a very attractive result if it holds).

##### Performance time of HyPrColoc when accounting for trait correlation

**Tables S2** displays the 1st, 5th (median) and 9th deciles of CPU time when assessing 7 of 10 traits sharing a CV, 3 traits not having a CV in the region, and all 10 traits being correlated (under the variety of scenarios outlined previously). For reference, the results when analysing independent traits are also included.

For scenarios in which analyses adjusted for observed correlation between summary data, the BB algorithm computed an inverse correlation matrix (Equation (3)) using a Cholesky decomposition, which is $\mathcal{O}(m^3)$ computationally expensive, for each of $\mathcal{O}(mQ^2)$ causal configurations. Recall that in our simulations we resampled phased haplotypes from one of the first 50 regions confirmed to be associated with CHD for each of 1000 datasets. These regions vary markedly in size, i.e. $Q$ varies between regions, hence the number of causal configurations considered varies between regions. This is reflected in the results (**Table S2**). For example, across all 50 regions, the median time to correctly identify the set of colocalized traits when adjusting for correlation in both the between study-level summary data and configuration priors was around 19 seconds. For larger regions ($Q \geq 2000$) the median time grew to over 100 seconds. These performance times contrast with the results from the algorithm when assuming independence between studies, thereby avoiding repeatedly computing $\mathcal{O}(m^3)$ expensive matrix inverses, in which the median algorithm time is $< 0.01$ seconds.

#### S4.2. Multiple causal variants per trait

In this section we focus on an assessment of the performance of the single causal variant (CV) assumption when one or more traits have two CVs in the genomic region. Our goal is to highlight the circumstances in which the BB algorithm regularly detects a true cluster of colocalized traits despite some or all traits having more than one CV (and vice versa, i.e. when the BB algorithm fails to identify a cluster of truly colocalized traits owing to violations of the single CV assumption).

#### Robustness of the single CV assumption when one or more traits have multiple CVs per region

Here we assess the performance of the single CV assumption when one or more traits have an additional CV in the region. As noted from **Figures 5-6**, the distribution of the HyPrColoc posterior probability of colocalization and the performance of the BB algorithm is related to: (i) the sample size of each study; (ii) trait variance explained by a CV and; (iii) LD between distinct CVs. In our present assessment, we focus on variations of (ii) and (iii), i.e. the impact of variance explained by an additional CV and the strength of LD between CVs. We omit assessing the influence of sample size here, i.e. scenario (i), owing to its implicit relationship trait variance explained by a CV (**Figures 5 and S3**).

Following the simulating strategy outlined previously (see main text **Model validation using simulations**), we generated 10 quantitative traits all of which share a single CV explaining 1% of trait variation and either 1, 2 or all 10 traits have an additional distinct CV. The additional CV explains either 0.5% or 1% of trait variation. This highlights the variability in performance when the secondary genetic association is either weaker (0.5%) or equal (1%) in strength to the shared CV. Let $r_i^2$ denote the LD between the shared CV (across all traits) and the additional CV for the $i^{th}$ trait only. To assess the impact of LD between the shared CV and any of the secondary CVs, we assigned a variant to be causal under two scenarios: (i) each additional CV for trait $i$ was restricted to $r_i^2 \leq 0.5$, reflecting scenarios in which traits have (approximately) a conditionally independent secondary signal and; (ii) no restriction on LD between the shared and any additional CV, i.e. $r_i^2 \leq 1$. Results are presented in **Table S5**.

**Linkage disequilibrium (LD) between additional CVs and a shared CV**

When the LD between an additional CV, for the $i^{th}$ trait, and the shared CV is bounded, i.e. $r_i^2 \leq 0.5$, the results indicate a sensitivity to (i) trait variation explained by an additional CV and (ii) the number of traits with additional CVs. Results appear to be more strongly dependent on trait variance explained, however (**Table S5**). For example, when each additional CV explains less trait variation than the shared variant, i.e. $0.5\%$ relative to 1%, the BB algorithm had continued good performance in detecting the true collection of colocalized traits: the probability of correctly identifying all colocalized traits is $\{0.86, 0.743, 0.704\}$ for $\{1, 5, 10\}$ traits with an additional CV respectively. Moreover, regardless of the number of traits with an additional CV, the probability of detecting at least 9 of 10 truly colocalized traits was $> 0.92$. Highlighting that the mis-specified single CV assumption is somewhat robust to the number of traits with additional CVs (provided the additional CVs explain less trait variation than the shared CV). When trait variance explained by an additional CV was comparable to the shared CV, i.e. both an additional and shared CV explained $1\%$ of trait variation, there was a steep decline in the BB algorithms ability to correctly identify all colocalized traits: the probability of correctly identifying all colocalized traits is $\{0.517, 0.06, 0.007\}$ for $\{1, 5, 10\}$ traits with an additional CV respectively. In general, any trait with two causal variants explaining equal amounts of trait variation were deemed not to colocalize with any other trait. This matches the theoretical results, that is $P_A \rightarrow 1$ if the Bayes Factor for the single shared causal variant is maximally dominant over all other SNP Bayes Factors per trait in the region (see **Alignment: distinguishing a shared causal variant from distinct causal variants in strong LD**).

When LD is unrestricted, i.e. $r_i^2 \leq 1$, performance is again more strongly affected by trait variation explained by an additional CV and the number of traits with additional CVs, than correlation between CVs. There is some effect of strong correlation, however. For example, the probability of correctly identifying all colocalized traits is $\{0.86, 0.743, 0.704\}$ when LD was restricted ($r_i^2 \leq 0.5$) and $\{0.846, 0.614, 0.46\}$ when LD was unrestricted ($r_i^2 \leq 1$), for $\{1, 5, 10\}$ traits with an additional CV respectively.

Overall, we conclude that the single CV assumption in multi-trait colocalization analyses recovers sensible results when either: (i) each trait has at most one CV in the region or; (ii) each trait has multiple CVs and a shared CV explains more trait variation than an additional CV, for each trait. In the following section we illustrate and example property (ii) using real data.

#### S4.3. CHD example: additional results

Here we present additional stacked association and fine-mapping plots from our applied analysis of CHD with 14 established risk-factors. We do this for three reasons:

i. To highlight that joint colocalization analyses can boost detection when subthreshold GWAS results are included in analyses (**Figure S6-8**).
ii. To highlight heterogeneity in the relationship between the colocalization posterior probability and fine-mapping evidence, i.e. strong support for a colocalization hypothesis does not imply the existence of a strong single causal variant fine-mapping probability (**Figure S7**).
iii. To illustrate potential in the continued good performance of HyPrColoc under a single causal variant framework despite empirical evidence suggesting the existence of multiple causal variants in the region for one or more traits (**Figure S8**).

#### S4.4. Properties of the HyPrColoc approximation

In this section, we explore the following:

i. Identification of the data generating mechanism(s) in which $\delta_R \longrightarrow 0$.
ii. Identification of the data generating mechanism in which $\delta_A \longrightarrow 0$.

iii. Verification that $P^c_{(m-1,1)} \longrightarrow 0$ as $\delta_R, \delta_A \longrightarrow 0$ from (11).

We address each point in turn. Note that (i) and (ii) are considered to inform our understanding of the regional and alignment statistics whereas (iii) underpins the validity the HyPrColoc approximation.

#### Regional and alignment statistics: a single shared causal variant across all traits

##### Regional statistic: shared genetic predictors across all traits

We write the joint Bayes Factor at SNP $i$, across all $m$ studies, as the product of BFs from each study, e.g.

$$BF\left(S_i \in \mathcal{S}_{H_m}\right) = \prod_{k=1}^{m} BF_i^{(k)}$$

( 9 )

This is the setup when the $k = 1,2, \ldots, m$ studies are independent (i.e. non-overlapping samples). When there are overlapping samples between studies, the JABF in Equation (3) can be written in the above form by orthogonalizing the $Z$-scores using a Cholesky factorisation. In this case, $k$ would index each of the $m$ transformed variables. For simplicity, henceforth we assume the traits are measured in independent studies.

Recall that $\delta_R = (1 - P_R)$. We re-write Equation (5), to yield

$$\delta_R = 1 - P_R = \frac{\sum_{i=0}^{m-1} P_{(i)}}{\sum_{i=0}^{m} P_{(i)}} = \frac{\chi_{m-1}}{1 + \chi_{m-1}},$$

Where $P_{(i)} = \frac{P(\mathcal{H}_{(i)}|D)}{P(H_0|D)}$ and

$$\chi_{m-1} = \sum_{i=0}^{m-1} \frac{P_{(i)}}{P_m} = \underbrace{\frac{P_{(m-1)}}{P_m}}_{\Delta\ 1\ \text{trait}\ w/CV} \left[ 1 + \underbrace{\frac{P_{(m-2)}}{P_{(m-1)}}}_{\Delta\ 1\ \text{trait}\ w/CV} \left[ 1 + \underbrace{\frac{P_{(m-3)}}{P_{(m-2)}}}_{\Delta\ 1\ \text{trait}\ w/CV} [1 + \cdots] \right] \right].$$

We will show later that $\frac{P_{(m-k)}}{P_{(m-k-1)}} = \mathcal{O}(\delta_R)$, hence the above illustrates that $\chi_{m-1}$ can be written as a power series in $\delta_R$. From the above, it follows therefore that:

$$\lim_{\delta_R \to 0} \left( \frac{1-\delta_R}{\delta_R} \right) \chi_{m-1} = 1$$

$$\Longrightarrow \chi_{m-1} \sim \delta_R,$$

$$\Longrightarrow \frac{P_{(m-1)}}{P_m} \sim \delta_R,$$

where $\sim$ denotes an asymptotic equivalence relation. The last line holds under the assumption of fixed prior configuration information. In this scenario, as $\delta_R \to 0$ only the likelihood contributions in each $P_{(i)}$ change and it follows (via a proof by contradiction) that:

$$\frac{P_{(m-1)}}{P_m} \gg \frac{P_{(m-2)}}{P_m} \gg \cdots \gg \frac{P_{(1)}}{P_m}, \qquad \delta_R \to 0.$$

The aim is now to identify the types of causal configurations satisfying the equivalence relation $\frac{P_{(m-1)}}{P_m} \sim \delta_R$. To do this we re-write $\frac{P_{(m-1)}}{P_m}$ in terms of prior configurations and Bayes Factors by combining Equations (2), (5) and (9), i.e.

$$\sum_{k=1}^{m} \alpha_k \sum_{i=1}^{Q} \frac{1 \big/ BF_i^{(k)}}{1 + \sum_{j \neq i} \frac{BF_j^{(1)} BF_j^{(2)} \cdots BF_j^{(m)}}{BF_i^{(1)} BF_i^{(2)} \cdots BF_i^{(m)}}} \sim \delta_R \ , \qquad \delta_R \longrightarrow 0$$

*( 10)*

where

$$\alpha_k = \frac{p_{12\cdots(k-1)(k+1)\cdots m}}{p_{12\cdots m}}.$$

Under the conditionally uniform prior $\alpha_k = 1$, which is independent of trait $k$. With $\alpha_k$ a fixed constant, Equation (10) makes clear that the parameter $\delta_R$ is a function of the Bayes Factors, *i.e.* $\delta_R = \delta_R(\boldsymbol{BF})$, where $\boldsymbol{BF}$ denotes the collection of all trait and SNP specific Bayes factors. On noting that

$$1 + \sum_{j \neq i} \frac{BF_j^{(1)} BF_j^{(2)} \cdots BF_j^{(m)}}{BF_i^{(1)} BF_i^{(2)} \cdots BF_i^{(m)}} \geq 1,$$

it follows that there must exist at least one SNP $j$ satisfying

$$BF_j^{(k)} \geq \mathcal{O}(\delta_R^{-1}), \quad j \in \{1,2,\dots,Q\}, \qquad k = 1,2,\dots,m,$$

$$\text{and a trait } k' : \quad BF_j^{(k')} = \mathcal{O}(\delta_R^{-1}), \quad k' \in \{1,2,\dots,m\}.$$

The above illustrates that the largest Bayes-Factors for each trait can grow at different asymptotic scales. To avoid this unnecessary complication, we perform a *model order reduction* technique that assumes there are two classes of BFs for *each* SNP $j$ and trait $k$, i.e.

$$BF_j^{(k)} = \begin{cases} \varepsilon_{jk}, & j \text{ not causal for trait } k \\ 1/\varepsilon_{jk}, & j \text{ causal (or in high LD w/causal variant) for trait } k \end{cases}$$

*( 11)*

where $\varepsilon_{ij}$ is given by

$$\varepsilon_{jk} = \tau_{jk}\delta_R + \mathcal{O}(\delta_R^2), \qquad \delta_R \longrightarrow 0,$$

and $\tau_{jk}$ is a positive $\mathcal{O}(1)$ constant that allows each BF to vary by SNP and trait. We identify causal configurations that satisfy the asymptotic equivalence relation by placing Equation (11) into Equation (10) under different data-generating mechanisms: (i) all traits share at least one causal variant; (ii) at most $m - 1$ traits share at least one CV; and so on. For each scenario, if this does not lead to a contradiction of the equivalence relation Equation (10) then we conclude that the mechanism is a candidate solution to $\delta_R = 0 \iff P_R = 1$.

1. **<u>All traits have at least one CV</u>**

Let SNP $i \in q$, where $q$ is the *set* of variants that are shared across all traits. If $|q| > 1$ this can be due to either: (i) strong LD between distinct CVs for each of the $m$ traits; (ii) multiple shared CVs across all traits or; (iii) a combination of both (i) and (ii).

Combining Equations (10) and (11), it follows that

$$\frac{1/BF_i^{(k)}}{1 + \sum_{j \neq i} \frac{BF_j^{(1)} BF_j^{(2)} \cdots BF_j^{(m)}}{BF_i^{(1)} BF_i^{(2)} \cdots BF_i^{(m)}}} \sim \tau_{ik} \left(1 + K_{i_q}\right) \delta_R, \qquad k = 1,2,\cdots,m,$$

where $K_{i_q}$ is a constant that appears when $|q| > 1$. However, if $i \notin q$, i.e. SNP $i$ is not causal (or in strong LD with a CV), then

$$\frac{1/BF_i^{(k)}}{1 + \sum_{j \neq i} \frac{BF_j^{(1)} BF_j^{(2)} \cdots BF_j^{(m)}}{BF_i^{(1)} BF_i^{(2)} \cdots BF_i^{(m)}}} \leq \mathcal{O}\left({\delta_R}^2\right), \qquad k = 1,2,\cdots,m.$$

These results reveal that

$$\sum_{k=1}^{m} \alpha_k \sum_{i=1}^{Q} \frac{1/BF_i^{(k)}}{1 + \sum_{j \neq i} \frac{BF_j^{(1)} BF_j^{(2)} \cdots BF_j^{(m)}}{BF_i^{(1)} BF_i^{(2)} \cdots BF_i^{(m)}}} \sim \delta_R \sum_{k=1}^{m} \alpha_k \sum_{i \in q} \tau_{ik} \left(1 + K_{i_q}\right) = \mathcal{O}(\delta_R).$$

That is, the equivalence relation Equation (10) is balanced when all traits contain at least one causal variant or when causal variants between traits are not identical but are in *strong LD* with one another.

2. **A maximum of $m-1$ traits have one or more CVs**

Following the same approach as before but now assuming the $i^{\text{th}}$ SNP is causal for all traits except trait *k*, it follows that

$$\frac{1/BF_i^{(k)}}{1+\sum_{j\neq i}\frac{BF_j^{(1)}BF_j^{(2)}\cdots BF_j^{(m)}}{BF_i^{(1)}BF_i^{(2)}\cdots BF_i^{(m)}}} \sim \frac{1+K_{i_q}}{\tau_{ik}\delta_R} \nsim \delta_R$$

which violates the equivalence relation Equation (10). Hence, as $P_R \longrightarrow 1$ the data generating mechanism cannot be one in which a maximum of $m-1$ traits share at least one causal variant; or subsets of the $m-1$ traits are driven by mutually exclusive causal variants that are in strong LD. The results for a maximum of $m-2$ traits; $m-3$ traits…follow similarly.

In summary, as $P_R \longrightarrow 1$, any of the following causal configurations are possible:

i. All traits colocalize at a single causal variant (main text **Figure 3**).
ii. There exists a region of two or more genetic predictors shared across all traits, owing to a complex LD structure. Within this shared region, subsets of all traits are driven by mutually exclusive causal variants (main text **Figure 3**).
iii. All traits share multiple causal variants in the region.
iv. A combination of (i) or (iii) and (ii).

**Alignment: distinguishing a shared causal variant from distinct causal variants in strong LD**

The alignment probability $P_A$ quantifies evidence between the scenarios (i)-(iv) above. We show this by identifying the data generating mechanism(s) which allows $\delta_A \longrightarrow 0$. We follow the same approach that identified the mechanisms which recover small values of $\delta_R$. To avoid unnecessary algebra, we outline the main steps here.

Re-writing Equation (6) as

$$\delta_A = 1 - P_A = \frac{P_{(m-1,1)}}{P_m + P_{(m-1,1)}} = \frac{\chi_{(m-1,1)}}{1+\chi_{(m-1,1)}},$$

returns

$$\chi_{(m-1,1)} = \frac{P_{(m-1,1)}}{P_m} \sim \delta_A, \quad \delta_A \to 0.$$

Writing the above in terms of Bayes Factors and configuration prior probabilities, by combing Equations (2), (6) and (9), we identify that

$$\sum_{k=1}^{m} \alpha'_k \sum_{i=1}^{Q} \frac{\sum_{j\neq i} \frac{BF_j^{(k)}}{BF_i^{(k)}}}{1 + \sum_{j\neq i} \frac{BF_j^{(1)} BF_j^{(2)} \cdots BF_j^{(m)}}{BF_i^{(1)} BF_i^{(2)} \cdots BF_i^{(m)}}} \sim \delta_A , \qquad \delta_A \longrightarrow 0,$$

*( 12 )*

with

$$\alpha'_k = \frac{p_{12\cdots(k-1)(k+1)\cdots m}\, p_k}{p_{12\cdots m}}.$$

The main difference between Equations (10) and (12) is in the numerator, which is now written in terms of Bayes Factor ratios $\frac{BF_j^{(k)}}{BF_i^{(k)}}$. We modify the model order reduction setup in Equation (11) to accommodate this, i.e. for SNPs $i$ and $j$ and the $k^{th}$ trait we assume that

$$BF_j^{(k)} \Big/ BF_i^{(k)} = \begin{cases} \varepsilon'_{ijk}, & i \text{ causal } j \text{ not causal for trait } k \\ \tau'_{ijk}, & i \text{ and } j \text{ not causal for trait } k \\ 1/\varepsilon'_{ijk}, & i \text{ not causal } j \text{ causal for trait } k \end{cases} \quad : \ \varepsilon'_{ijk} = \tau'_{ijk}\delta_A + \mathcal{O}(\delta_A^2) \quad \text{and} \quad \delta_A \longrightarrow 0.$$

*( 13)*

Note, for example, that "$i$ causal $j$ not causal for trait $k$" should also be interpreted as "SNP $i$ in strong LD with a causal variant whilst SNP $j$ is not causal and in low LD with a casual SNP for trait $k$". To determine the data generating mechanism which returns a large value of the alignment probability $P_A$, we place Equation (13) into (12) under the following options: (i) all traits share one CV; (ii) all traits share two or more CVs; or (iii) all traits have at least one causal variant and at most $m-1$ traits share a CV. As previously, a mechanism which violates the equivalence relation (16) is dismissed as solution to $\delta_A = 0$. The results (omitted) reveal that: large values of the alignment statistic, i.e. as $P_A \longrightarrow 1$, can be *only* be explained by each trait having a single shared causal variant against the alternatives (ii) and (iii).

**Conclusion**

Small values of the regional probability $\delta_R$ guarantee that all traits must have at least one causal variant in the region and these causal variants are shared or in strong LD across all $m$ traits. Whereas, small values of the alignment probability $\delta_A$ guarantee that all traits with at least one causal variant in the region must only share a single causal variant. Hence, taken in combination, as $\delta_R \longrightarrow 0$ and $\delta_A \longrightarrow 0$ the data are generated *only* under a single causal variant colocalization mechanism shared across all traits.

#### Deprioritised hypotheses are monotonic decreasing in regional and alignment probabilities

The ratio $\frac{P^c_{(m-1,1)}}{P_{(m-1,1)}}$ in Equation (7) can be written as the product of ratios in which there is a change in the maximum number of traits with a causal variant, e.g. $\frac{P_{(j-1,1)}}{P_{(j,1)}}$, multiplied by a product sum of ratios in which there is a change in the location of one causal SNP, e.g. $\frac{P_{(j-2,1,1)}}{P_{(j-1,1)}}$,

$$\frac{P^c_{(m-1,1)}}{P_{(m-1,1)}} = \underbrace{\frac{P_{(m-2,2)} + P_{(m-2,1,1)}}{P_{(m-1,1)}}}_{\Delta\ \text{CV position}} \left[1 + \frac{P_{(m-3,3)} + P_{(m-3,2,1)} + \cdots P_{(m-3,1,1,1)}}{P_{(m-2,2)} + P_{(m-2,1,1)}} [[1 + \cdots]]\right]$$

$$+ \underbrace{\frac{P_{(m-2,1)}}{P_{(m-1,1)}}}_{\Delta\ \text{traits with CV}} \left( \left[ 1 + \frac{P_{(m-3,2)} + P_{(m-3,1,1)}}{P_{(m-2,1)}} [[1 + \cdots]] \right] + \right.$$

$$\left. \cdots \frac{P_{(j-1,1)}}{P_{(j,1)}} \left[ 1 + \frac{P_{(j-2,2)} + P_{(j-2,1,1)}}{P_{(j-1,1)}} [1 + \cdots] \right] \right).$$

*(14)*

After some algebra, it can be shown, using Equations (10) and (12), that the ratio in which there is a change in the number of traits with a causal variant is controlled by $\delta_R$, e.g.

$$\frac{P_{(m-2,1)}}{P_{(m-1,1)}} = \mathcal{O}(\,\delta_R)$$

and a change in the position of a causal variant is controlled by $\delta_A$, e.g.

$$\frac{P_{(m-2,2)} + P_{(m-2,1,1)}}{P_{(m-1,1)}} = \mathcal{O}(\,\delta_A).$$

Placing the above into Equation (14), it follows that as $\delta_R, \delta_A \longrightarrow 0$, the posterior probability of colocalization Equation (7) can be written as:

$$P\left(H_{scv}^{(m)} \middle| D\right) = \frac{P_{scv}^{(m)}}{P_{scv} + P_{dcv}}$$

$$= \frac{P_R P_A}{1 - \left( (1 - P_R)(1 - P_A) - P_R(1 - P_A) \frac{P_{dcv}^{(m;\, m-1)^c}}{P_{dcv}^{(m;\, m-1)}} \right)}$$

$$= P_R P_A + \mathcal{O}(\delta_A^2 + \delta_R \delta_A), \qquad \delta_R\,, \delta_A \longrightarrow 0.$$

#### Linearization of the regional statistic

A complete computation of the regional probability is $\mathcal{O}(2^m Q)$ expensive in the number of traits $m$ and variants in a region $Q$. This becomes computationally impractical as the number of traits becomes large, e.g. $m \geq 20$. In keeping with the spirit of hypothesis prioritisation, we now show that the regional probability is accurately approximated by a probability that has linear computational cost in the number of traits $m$. This maintains computationally tractability of the HyPrColoc approximation and algorithm across large numbers of traits. The regional probability $P_R$ from Equation (5) is given by

$$P_R = \frac{P_m}{1 + P_m + P_{(m-1)} + \sum_{k=1}^{m-2} P_{(i)}} = \frac{P_m}{1 + P_m + P_{(m-1)}} \left( \frac{1}{1 + \chi'_{m-2}} \right)$$

with

$$\chi'_{m-2} = \sum_{k=1}^{m-2} \frac{P_{(i)}}{1 + P_m + P_{(m-1)}} = \mathcal{O}(\delta_R^2), \quad \delta_R \to 0,$$

on using Equation (10). Consequently,

$$P_R = \frac{P_m}{1 + P_m + P_{(m-1)}} \left(1 + \mathcal{O}(\delta_R^2)\right).$$

The denominator $1 + P_m + P_{(m-1)}$ is computed using only $\mathcal{O}(mQ)$ causal configurations, which is linear in the number of traits and variants in the region. We therefore use

$$P'_R = \frac{P_m}{1 + P_m + P_{(m-1)}} = P_R\left(1 + \mathcal{O}(\delta_R^2)\right)$$

to approximate the regional statistic. As an option, users of HyPrColoc can assess the accuracy of $P'_R$ against $P_R$ by increasing the number of hypotheses used to approximate $P_R$, i.e. we can include $P_{(m-2)}$ in the denominator (e.g. using $1 + P_m + P_{(m-1)} + P_{(m-2)}$) which increases the number of causal configurations used to approximate $P_R$ from $\mathcal{O}(mQ)$ up to $\mathcal{O}(m^2Q)$. **Table S4** illustrates the performance of the linear approximation (using the simulation setup outlined in the **Online Methods)**, indicating a very close correspondence between $P'_R$ and $P_R$ when a shared CV explains only 0.5% of trait variation for each trait (median absolute difference

0.0017), and when we increase trait variance explained to 1%, $P_R'$ and $P_R$ are numerically identical.

### References

1. Giambartolomei, C. *et al.* Bayesian Test for Colocalisation between Pairs of Genetic Association Studies Using Summary Statistics. *PLoS Genet.* **10**, (2014).

2. Wakefield, J. Bayes Factors for Genome-Wide Association Studies : Comparison with P -values. **86**, 79–86 (2009).

3. Pickrell, J. K. *et al.* Detection and interpretation of shared genetic influences on 42 human traits. *Nat Genet* **48**, 709–717 (2016).

4. Province, M. A. & Borecki, I. B. A correlated meta-analysis strategy for data mining 'OMIC' scans. *Pac. Symp. Biocomput.* 236–46 (2013).

5. Giambartolomei, C. *et al.* A Bayesian framework for multiple trait colocalization from summary association statistics. *Bioinformatics* **34**, 2538–2545 (2018).

6. The 1000 Genomes Project Consortium. A global reference for human genetic variation. *Nature* **526**, 68–74 (2015).

7. The CARDIoGRAMplusC4D Consortium. Large-scale association analysis identifies new risk loci for coronary artery disease. *Nat. Genet.* **45**, 25–33 (2012).

8. Nikpay, M., Goel, A., Won, H.-H. & Hall, L. M. A comprehensive 1000 Genomes-based genome-wide association meta-analysis of coronary artery disease. *Nat. Genet.* **47**, 1121–1130 (2015).

### Tables

Table S1: GWAS datasets used to map the risk of CHD.

| **Phenotype** | **Abbreviation** | **Data Source** | **PubMed ID** | **N** |
|---|---|---|---|---|
| **Body mass index** | BMI | UK Biobank | NA | 336,107 |
| **Coronary heart disease** | CHD | CARDIoGRAM plusC4D | 26343387 | 184,305 |
| **Diastolic blood pressure** | DBP | UK Biobank | NA | 317,756 |
| **Education years** | EDU | SSAGC | 27225129 | 328,917 |
| **Estimated glomerular filtration rate** | eGFR | CKDGen | 28452372 | 110,517 |
| **Fasting glucose** | FG | MAGIC | 20081858 | 46,186 |
| **Fasting insulin** | FI | MAGIC | 20081858 | 46,186 |
| **High density lipoprotein** | HDL | GLGC | 20686565 | 99,900 |
| **Low density lipoprotein** | LDL | GLGC | 20686565 | 95,454 |
| **Rheumatoid arthritis** | RA | Okada *et al.* | 24390342 | 58,284 |
| **Smoking** | SMK | TAG | 20418890 | 74,035 |

| | | | | |
|---|---|---|---|---|
| **Systolic blood pressure** | SBP | UK Biobank | NA | 317,754 |
| **Triglycerides** | TG | GLGC | 20686565 | 96,598 |
| **Type II diabetes** | T2D | DIAGRAM | 28566273 | 159,208 |
| **Waist circumference** | WC | UK Biobank | NA | 336,639 |

The UK Biobank results were obtained from the first release of the Neale Lab's GWAS analysis of UK Biobank (http://www.nealelab.is/uk-biobank).

**Table S2:** Performance of the BB algorithm when 7 of 10 traits share a CV (the remaining 3 traits do not have a CV in the region), all traits are correlated (compound symmetric residual correlation matrix with correlation parameter $\rho = 0.5$) and the participants are *completely* overlapping between the studies. For comparison, the results when all traits are independent, i.e. $\rho = 0$, and the participants are non-overlapping between studies are presented. When the traits are correlated, the data were analysed in three ways: (i) accounting for correlation through the likelihood terms only; (ii) through the likelihood and prior configurations and; (iii) wrongly assuming independence. Presented are the detection probabilities and the 1st, 5th (median) and 9th deciles of the algorithm computation time, and the distribution of the posterior probability of colocalization (when the algorithm correctly identified all colocalized traits only).

| **Prior** | **Trait correlation ($\rho$)** | **Likelihood adjusted for trait correlation** | **Prior adjusted for trait correlation** | **Regional / alignment threshold** | **Multiple clusters of colocalized traits** | **Cluster contains 6 of the 7 colocalized traits** | **Correctly identified cluster of colocalized traits** | **Cluster has 1 false positive** | **Cluster has 2 or more false positives** | **CPU time (seconds)** | **Distribution of posterior** |
|---|---|---|---|---|---|---|---|---|---|---|---|
| | | | | | **Detection Probability** | | | | | $1^{st}$ ,$5^{th}$ and $9^{th}$ deciles | |
| **VS** | Yes (0.5) | Yes | No | 0.5 | 0.855 | 0.108 | 0.035 | 0.002 | 0 | (2.4, 3.9, 106) | (0.5, 0.54, 0.62) |
| **VS** | Yes (0.5) | Yes | Yes | 0.5 | 0.005 | 0.023 | 0.938 | 0.03 | 0.004 | (5.8, 19.3, 106) | (0.7, 0.87, 94) |
| **VS** | Yes (0.5) | No | No | 0.5 | 0.009 | 0.013 | 0.975 | 0.003 | 0 | $< 0.1$ | (0.79,0.96,1) |
| **VS** | No (0) | -- | -- | 0.5 | 0.017 | 0.017 | 0.955 | 0.011 | 0 | $< 0.1$ | (0.73,0.94,1) |

| | | | | | | | | | | | |
|---|---|---|---|---|---|---|---|---|---|---|---|
| **CU** | Yes (0.5) | Yes | No | 0.5 | 0.074 | 0.21 | 0.678 | 0.033 | 0.004 | (5.8, 19.3, 106) | (0.52, 0.61, 0.71) |
| **CU** | Yes (0.5) | Yes | Yes | 0.5 | 0.004 | 0.023 | 0.935 | 0.03 | 0.004 | (5.8, 19.4, 106) | (0.72, 0.84, 0.9) |
| **CU** | Yes (0.5) | No | No | 0.7 | 0.002 | 0.005 | 0.947 | 0.044 | 0.002 | <0.1 | (0.94,0.99,1) |
| **CU** | No (0) | -- | -- | 0.7 | 0.002 | 0.008 | 0.944 | 0.044 | 0.002 | <0.1 | (0.90,0.98,1) |

**Table S3:** Performance of the BB algorithm when 7 of 10 traits share a CV, the remaining 3 traits do not have a CV in the region, all traits are correlated (approximately compound symmetric with correlation parameter $\rho \in \{-0.1, 0.8\}$) and the genetic samples are completely overlapping between the studies. For comparison, the results when all traits are independent, i.e. $\rho = 0$, are shown. In all scenarios the traits were analysed as independent. Present are detection probabilities and the 1st, 5th (median) and 9th deciles of the posterior probability of colocalization (when the algorithm correctly identified all colocalized traits only).

| **Prior** | **Trait correlation ($\boldsymbol{\rho}$)** | **Likelihood adjusted for trait correlation** | **Prior adjusted for trait correlation** | **Regional / alignment threshold** | **Multiple clusters of colocalized traits** | **Cluster contains 6 of the 7 colocalized traits** | **Correctly identified cluster of colocalized traits** | **Cluster has 1 false positive** | **Cluster has 2 or more false positives** | **Distribution of posterior** |
|---|---|---|---|---|---|---|---|---|---|---|
| | | | | | **Detection Probability** | | | | | 1st ,5th and 9th deciles |

| | | | | | | | | | | |
|---|---|---|---|---|---|---|---|---|---|---|
| **VS** | Yes (-0.1) | No | No | 0.5 | 0.021 | 0.014 | 0.957 | 0.008 | 0 | (0.718, 0.95, 1) |
| **VS** | Yes (0.8) | No | No | 0.5 | 0 | 0.0015 | 0.995 | 0.003 | 0 | (0.84, 0.97, 1) |
| **VS** | No (0) | -- | -- | 0.5 | 0.017 | 0.017 | 0.955 | 0.011 | 0 | (0.73,0.94,1) |
| **CU** | Yes (-0.1) | No | No | 0.7 | 0 | 0.002 | 0.953 | 0.045 | 0 | (0.9, 0.987, 1) |
| **CU** | Yes (0.8) | No | No | 0.7 | 0 | 0 | 0.982 | 0.015 | 0.002 | (0.95, 0.994, 1) |
| **CU** | No (0) | -- | -- | 0.7 | 0.002 | 0.008 | 0.944 | 0.044 | 0.002 | (0.90,0.98,1) |

**Table S4:** The absolute difference between the linearized approximation of the regional statistic (which computes $mQ$ causal configurations) and the full computation (which computes $\mathcal{O}(2^m Q)$ causal configurations), for 10 traits which share a CV under two scenarios: the CV explains $0.5\%$ or $1\%$ of trait variance with sample sizes of $N = 10000$ and $5000$ respectively. Presented are the 1st , 5th and 9th deciles. By construct the absolute difference is smaller under the CU prior framework than the VS framework, i.e. $|P'_R - P_R|_{CU} \leq |P'_R - P_R|_{VS}$, we therefore do not present these results.

| **Prior** | **Sample size** ($\boldsymbol{N}$) | **Number of traits** ($\boldsymbol{m}$) | **Variance explained by CV** | **Correlation between traits** ($\boldsymbol{\rho}$) | **Absolute difference** ($\boldsymbol{P'_R - P_R}$) |
|---|---|---|---|---|---|
| | | | | | 1st, 5th and 9th deciles |
| **VS** | 10000 | 10 | 1% | 0.1 | (0, 0, 0) |
| **VS** | 5000 | 10 | 0.5% | 0.1 | (0.00003, 0.0017, 0.03) |

**Table S5:** Simulation of 10 traits sharing a CV with either 1, 5 or all 10 traits having an additional CV in the region. In all scenarios, each additional CV is distinct between traits (i.e. not shared). The shared CV explains 1% of trait variation and any additional CV explains either 0.5% or 1% of trait variance. The additional CVs were selected to have either restricted LD with the shared CV, i.e. $r^2 < 0.5$, or unrestricted LD, i.e. $r^2 \leq 1$. Presented are results from the BB algorithm using variant specific prior configuration probabilities ($\gamma = 0.98$), setting the regional and alignment thresholds to $P_R^* = P_A^* = 0.5$ and assuming each of the 10 traits has at most a single CV in the region.

| **Linkage disequilibrium between CVs within each trait** | **Traits with a shared CV** | **Traits with an additional *distinct* CV** | **No colocalized traits** | **Multiple clusters of colocalized traits** | **Cluster contains 7 of the 10 colocalized traits** | **Cluster contains 8 of the 10 colocalized traits** | **Cluster contains 9 of the 10 colocalized traits** | **Correctly identified cluster of colocalized traits** | **Distribution of posterior for correctly identified traits** |
|---|---|---|---|---|---|---|---|---|---|
| | CV explains ($\alpha$ %) of trait variance | | **Detection Probability** | | | | | | $1^{st}$, $5^{th}$ and $9^{th}$ deciles |
| $\mathbf{r^2 < 0.5}$ | 10 (1%) | 1 (0.5%) | 0 | 0.008 | 0.003 | 0.005 | 0.124 | 0.86 | (0.75,0.95,1) |
| $\mathbf{r^2 < 0.5}$ | 10 (1%) | 5 (0.5%) | 0 | 0.001 | 0.011 | 0.043 | 0.202 | 0.743 | (0.783, 0.97, 1) |
| $\mathbf{r^2 < 0.5}$ | 10 (1%) | 10 (0.5%) | 0 | 0.002 | 0.013 | 0.06 | 0.221 | 0.704 | (0.818, 0.98, 1) |

| | | | | | | | | | |
|---|---|---|---|---|---|---|---|---|---|
| $\mathbf{r^2 < 0.5}$ | 10 (1%) | 1 (1%) | 0 | 0.009 | 0.004 | 0.005 | 0.465 | 0.517 | (0.66,0.94,1) |
| $\mathbf{r^2 < 0.5}$ | 10 (1%) | 5 (1%) | 0 | 0.143 | 0.256 | 0.311 | 0.23 | 0.06 | (0.56,0.84,0.99) |
| $\mathbf{r^2 < 0.5}$ | 10 (1%) | 10 (1%) | 0 | 0.632 | 0.181 | 0.124 | 0.056 | 0.007 | (0.63, 0.78, 0.96) |
| $\mathbf{r^2 \leq 1}$ | 10 (1%) | 1 (0.5%) | 0 | 0.008 | 0 | 0.006 | 0.146 | 0.84 | (0.67, 0.94, 1) |
| $\mathbf{r^2 \leq 1}$ | 10 (1%) | 5 (0.5%) | 0 | 0.027 | 0.031 | 0.094 | 0.234 | 0.614 | (0.66, 0.94, 1) |
| $\mathbf{r^2 \leq 1}$ | 10 (1%) | 10 (0.5%) | 0 | 0.084 | 0.07 | 0.141 | 0.245 | 0.46 | (0.62, 0.93, 1) |
| $\mathbf{r^2 \leq 1}$ | 10 (1%) | 1 (1%) | 0 | 0.017 | 0.005 | 0.015 | 0.452 | 0.511 | (0.64,0.92,1) |
| $\mathbf{r^2 \leq 1}$ | 10 (1%) | 5 (1%) | 0 | 0.203 | 0.267 | 0.297 | 0.188 | 0.045 | (0.59,0.9,1) |
| $\mathbf{r^2 \leq 1}$ | 10 (1%) | 10 (1%) | 0 | 0.77 | 0.128 | 0.078 | 0.02 | 0.004 | (0.54,0.61,0.88) |

### Figures

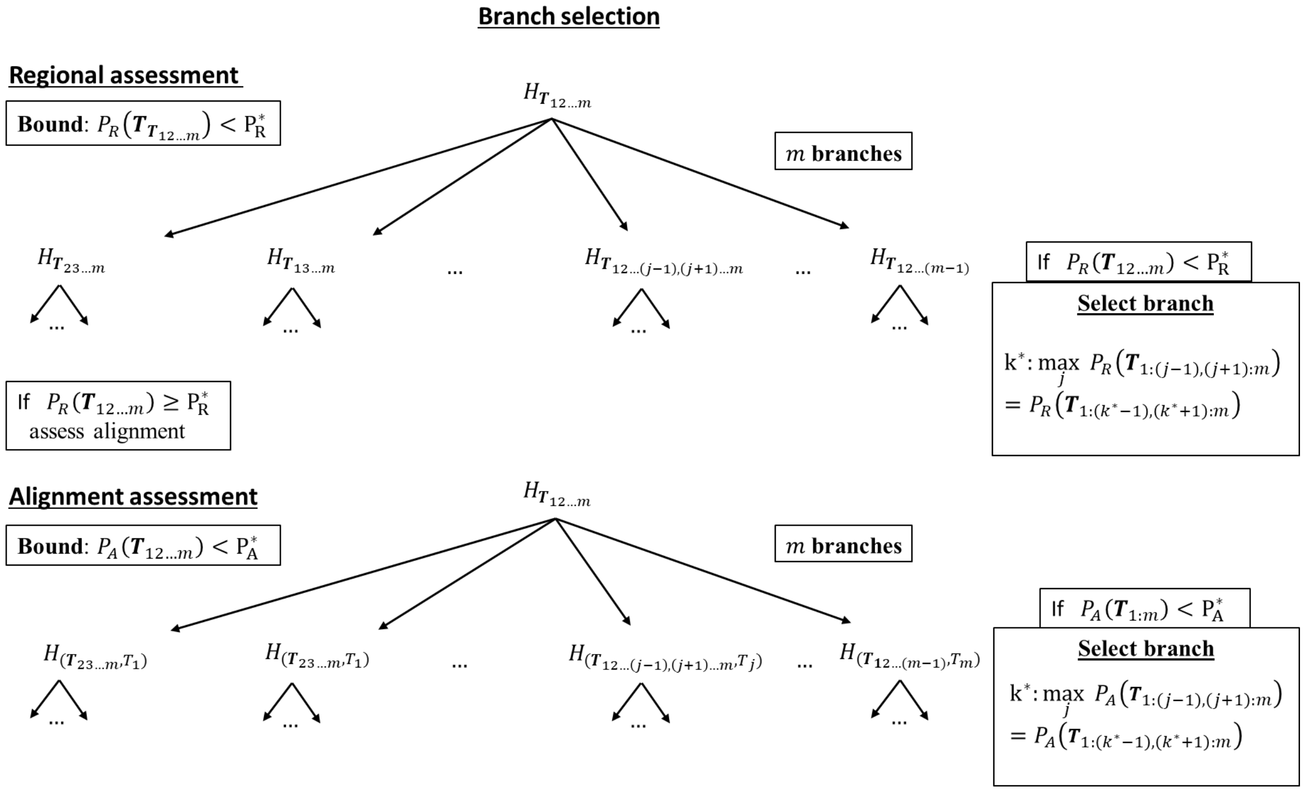

**Figure S1:** Illustration of the regional and alignment assessment in phase 2 of the BB algorithm. The algorithm starts by assessing evidence supporting the hypothesis that all $m$ traits share a CV, i.e. $H_{\boldsymbol{T}_{12\ldots m}}$, by computing a regional statistic $P_R(\boldsymbol{T}_{12\ldots m})$ from $m$ additional hypotheses $H_{\boldsymbol{T}_{12\ldots(j-1)(j+1)\ldots m}} : \; j = 1, 2, \ldots, m$. If $P_R(\boldsymbol{T}_{12\ldots m}) < P_R^*$, the default selection algorithm choses the branch (i.e. hypothesis) with the greatest support from the $m$ hypotheses in which $m-1$ traits share a CV and the remaining trait does not have a CV in the region, i.e. $H_{\boldsymbol{T}_{12\ldots(k^*-1)(k^*+1)\ldots m}}$, and repeats the process but now by assessing evidence supporting the hypothesis $H_{\boldsymbol{T}_{12\ldots(k^*-1)(k^*+1)\ldots m}}$. If $P_R(\boldsymbol{T}_{12\ldots m}) \geq P_R^*$, alignment at a single CV is assessed from $the\ m$ hypotheses $H_{(\boldsymbol{T}_{12\ldots(j-1)(j+1)\ldots m}, T_j)} : \; j = 1, 2, \ldots, m$. If $P_A(\boldsymbol{T}_{12\ldots m}) < P_A^*$ the selection algorithm choses the branch with the greatest support from all $m$ hypotheses in which $m-1$ traits share a CV and the remaining trait has a distinct CV, i.e. $H_{(\boldsymbol{T}_{12\ldots(k^*-1)(k^*+1)\ldots m}, T_{k^*})}$. These steps are then repeated, but now assessing evidence supporting the hypothesis $H_{\boldsymbol{T}_{12\ldots(k^*-1)(k^*+1)\ldots m}}$.

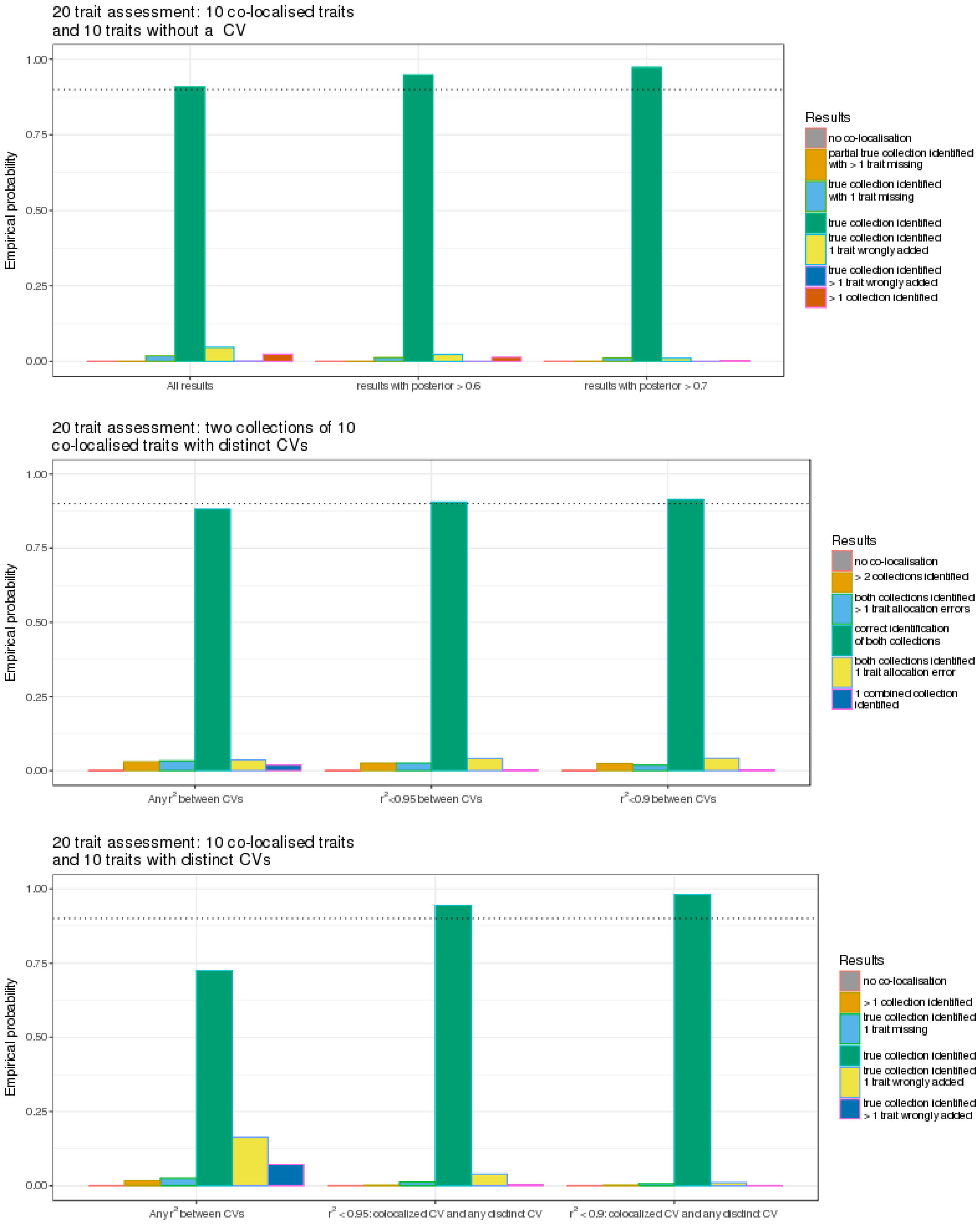

**Figure S2: Assessing the performance of the BB algorithm**. Three scenarios are simulated, in each scenario $m = 20$ traits with non-overlapping samples were generated, all traits had a study sample size of $N = 10000$, and variant-level priors were used in all analyses. In each scenario there exists at least one collection of 10 traits which share a causal variant and either: the remaining 10 traits do not have a causal variant in the region (top); there exists another collection of 10 traits which share a distinct causal variant (middle) and; all remaining traits have distinct causal variants from one another (bottom). Where indicated, detection probabilities are presented by linkage disequilibrium ($r^2$) between the causal variant, shared across the 10 (default) colocalized traits, and any other causal variant, i.e. when $r^2 \leq (1, 0.95, 0.9)$. When only the default colocalized causal variant is present, the detection probability is presented by posterior probability of colocalization, i.e. posterior $\geq (0.5, 0.6, 0.7)$.

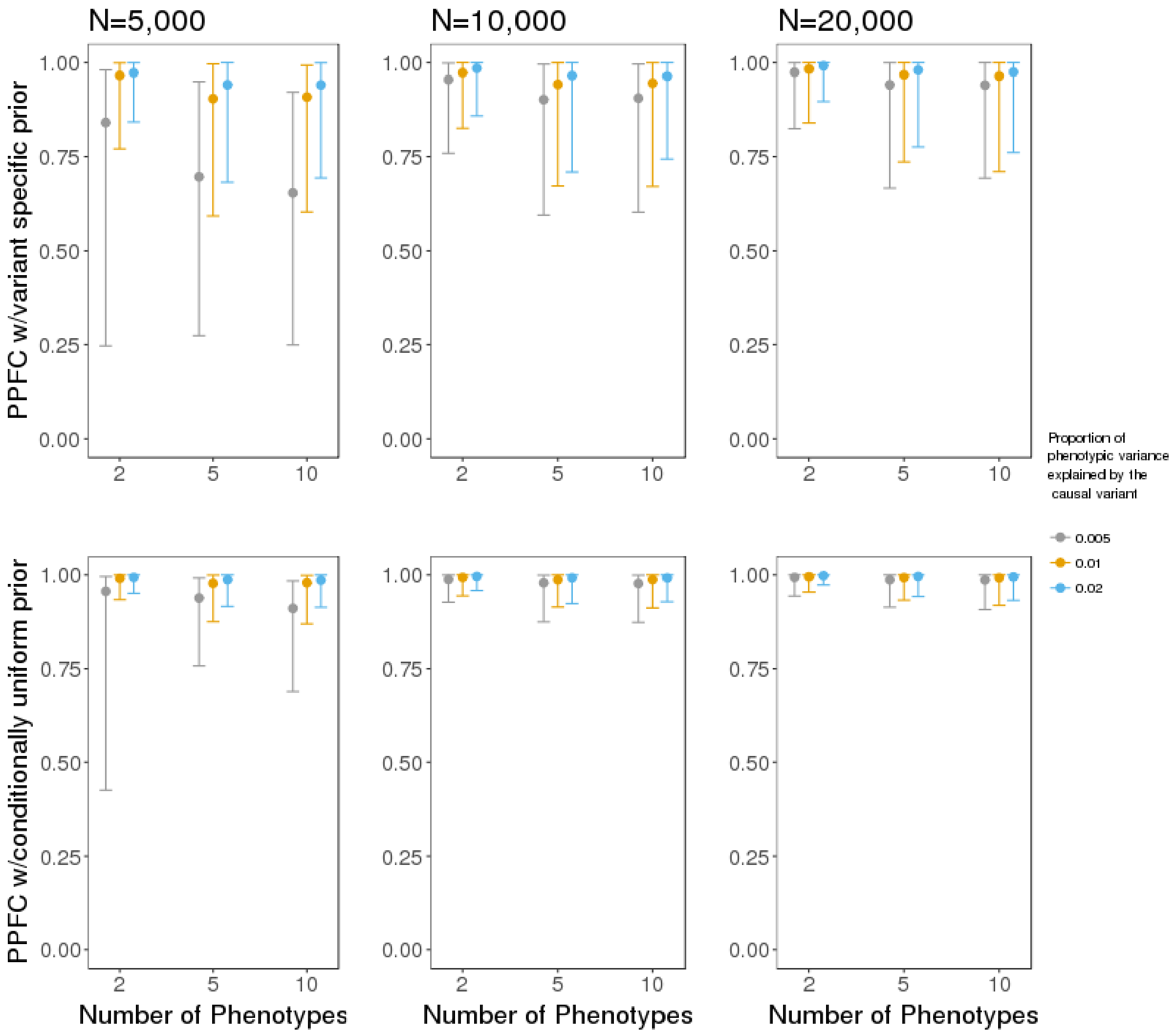

**Figure S3**: Comparison of the distribution of the posterior probability of colocalization when using the variant specific (top) and the conditionally uniform prior configuration probabilities. Plotted are the first, fifth (median), and ninth deciles for $m = 2, 5, 10$ traits, sample sizes of $N = 5{,}000, 10{,}000, 20{,}000$ and a single shared CV that explains either 0.5%, 1% and 2% of traits variance.

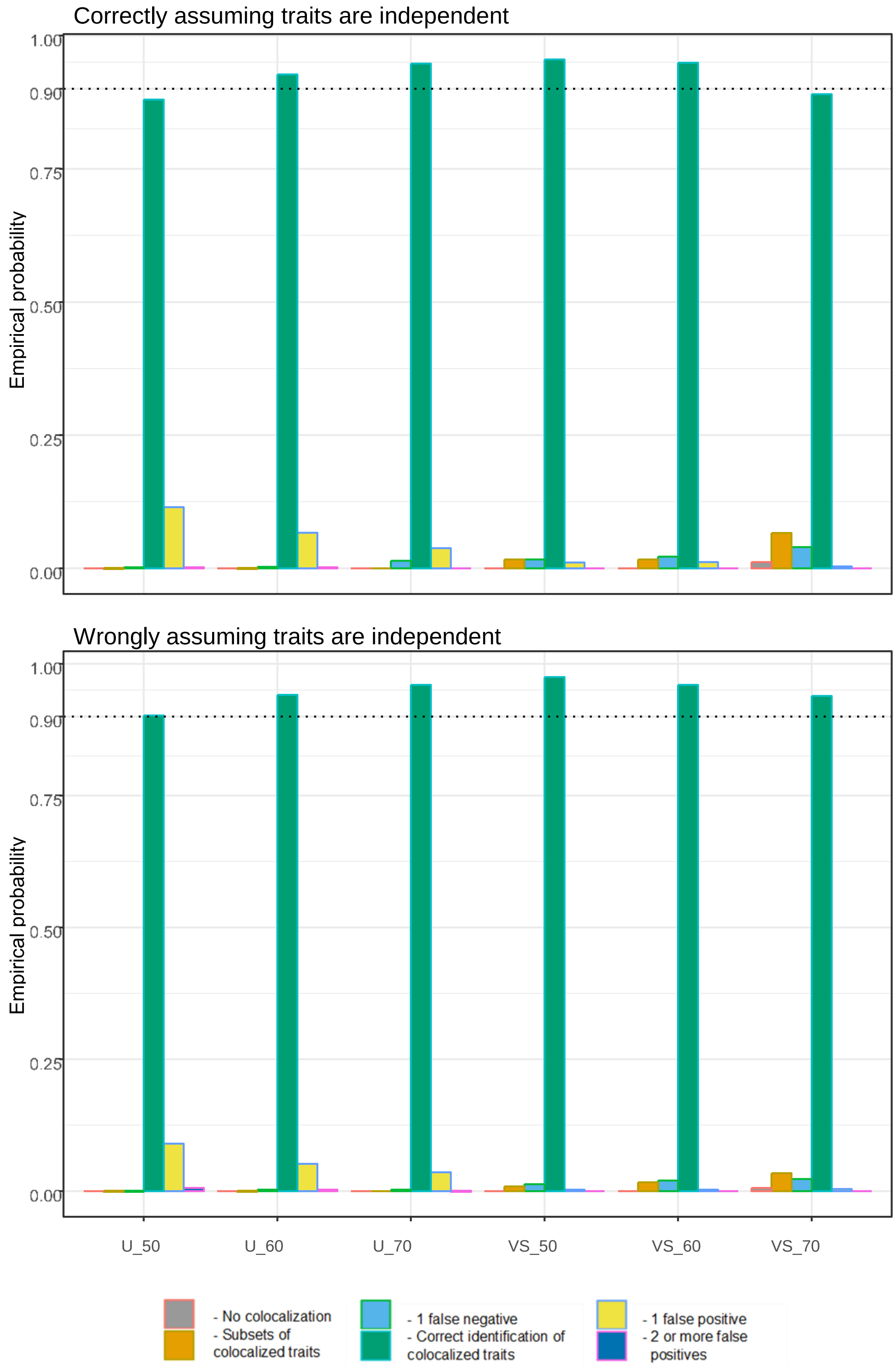

**Figure S4:** Comparison of algorithm performance when *correctly* or *incorrectly* assuming non-overlapping participants between studies (and *a priori* independence between traits). Presented is the probability of detecting clusters of colocalized traits for various values of the regional and alignment thresholds, *i.e.* $P_R^* = P_A^*$ and $P_R^*, P_A^* \in \{0.5, 0.6, 0.7\}$, under the variants specific (VS) and conditionally uniform (CU) prior configuration probabilities, using p=$10^{-4}$ and $\gamma = 0.98$. Note that, $U\_50$ denotes that the conditionally uniform priors were used with $P_R^* = P_A^* = 0.5$, and $VS\_50$ that the variant specific priors were used with $P_R^* = P_A^* = 0.5$, etc… Study sample were either correctly (**top**) or incorrectly (**bottom**) assumed to be non-overlapping and in all scenarios 7 of 10 traits share a single CV, the remaining traits did not have a CV.

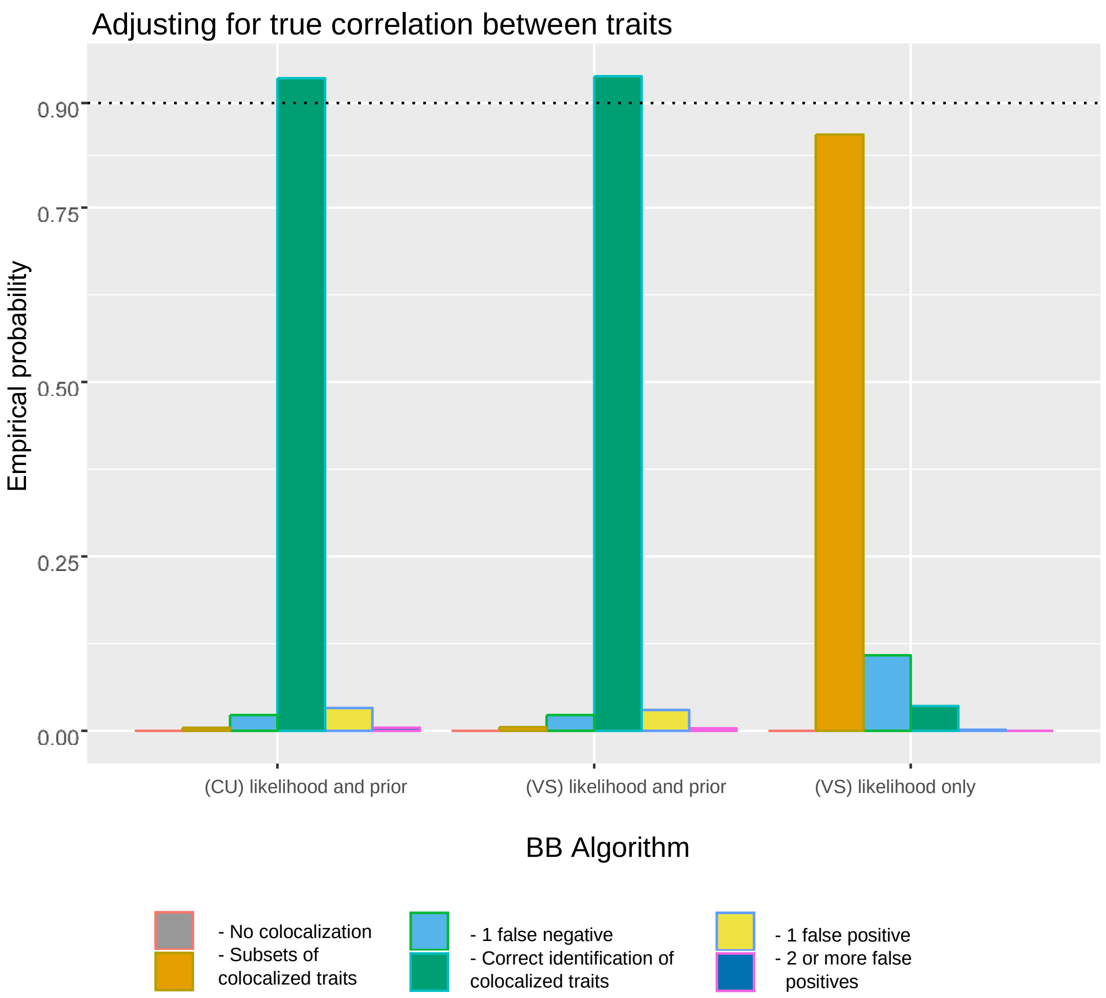

**Figure S5:** Accounting for overlapping participants between studies and *a priori* trait correlation in the BB algorithm. Presented is the probability of detecting colocalized traits when accounting for overlapping samples via the likelihood ( "likelihood only") or via the likelihood *and* prior configurations ("likelihood and prior") using the variant specific (VS) and conditionally uniform (CU)

prior configuration probabilities. In all scenarios the regional and alignment thresholds were set to $P_R^* = P_A^* = 0.5$ with p=$10^{-4}$ and $\gamma = 0.98$.

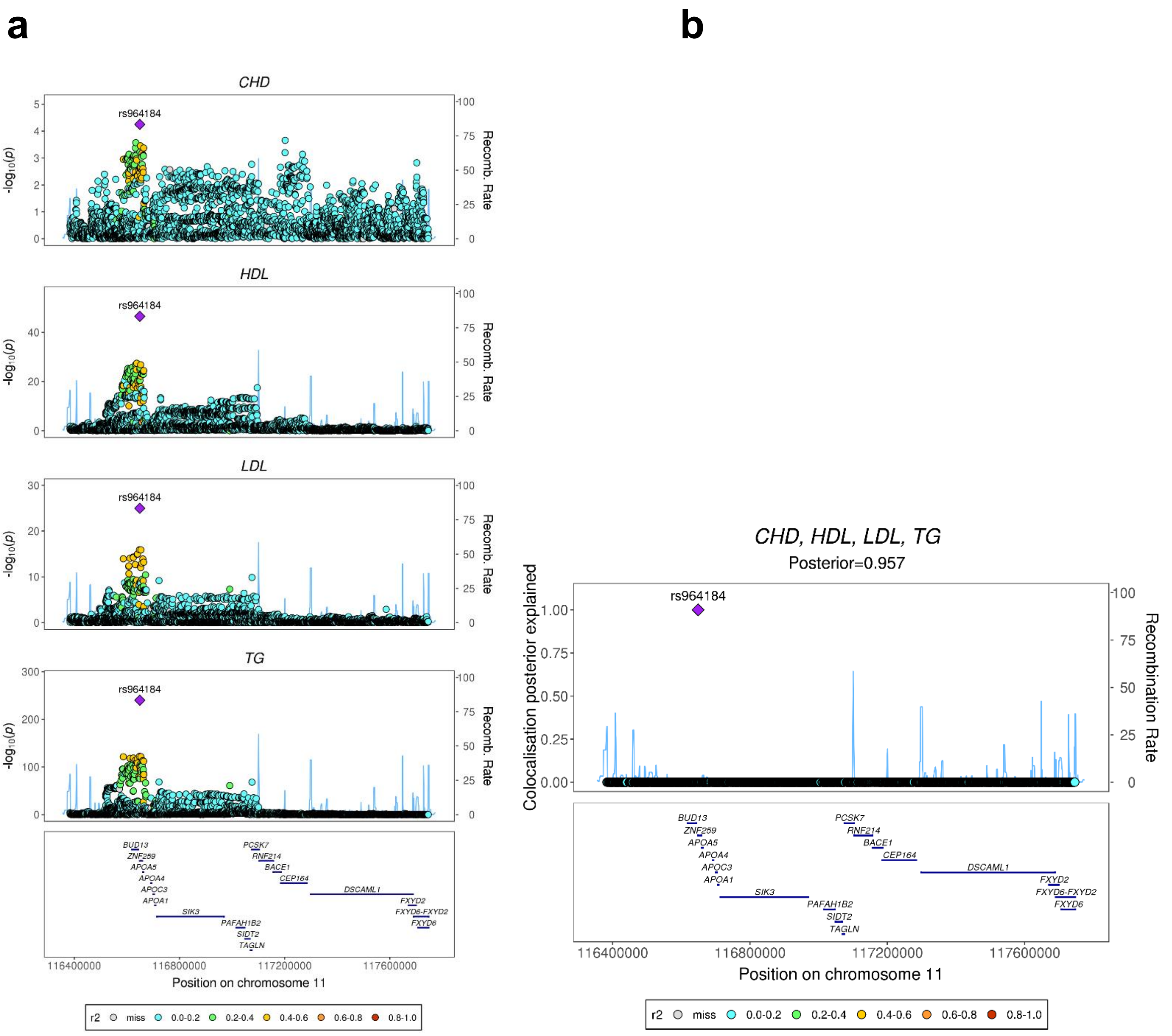

**Figure S6:** Visually striking example of borrowing strength between related traits. (**a**) Stacked association plots of CHD with high density lipoprotein (HDL), low density lipoprotein (LDL) and triglycerides (TG). The lead CHD variant rs964184 has nominal association strength in the CHD study[8]. However, on jointly analysing CHD and many related risk factors, HyPrColoc identifies a region which clearly highlights that borrowing information across related traits can boost power to detect colocalization and candidate causal variants. (**b**) The posterior probability of colocalization between CHD and the lipid fractions is 0.957 with the candidate causal variant rs964184 explaining almost 100% of the colocalization posterior probability.

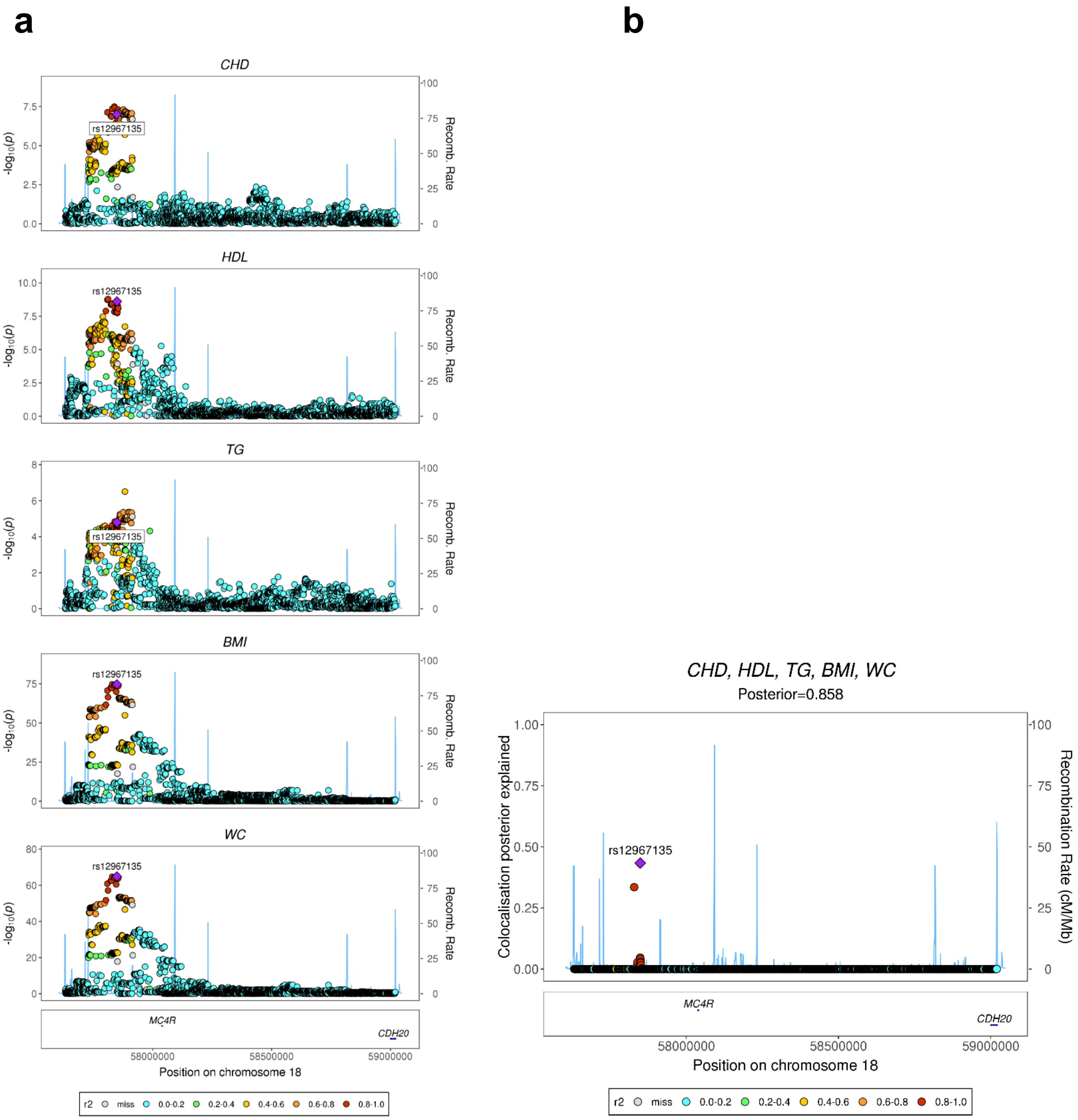

a
b
CHD
HDL
TG
BMI
WC
rs12967135
-log10(p)
Recomb. Rate
MC4R
CDH20
58000000
58500000
59000000
Position on chromosome 18
r2 miss 0.0-0.2 0.2-0.4 0.4-0.6 0.6-0.8 0.8-1.0
CHD, HDL, TG, BMI, WC
Posterior=0.858
Colocalisation posterior explained
Recombination Rate (cM/Mb)

**Figure S7**: A single shared associated region across traits with no definitive candidate causal variant. (**a**) Stacked association plots of CHD with high density lipoprotein (HDL), triglycerides (TG), body mass index (BMI) and waist circumference (WC). All traits share a single ‘peaked’ association region (regional probability $P_R$ =0.98) with a single causal variant colocalization mechanism appearing likely (posterior probability 0.86). (**b**) Despite the candidate variant rs12967135 explaining nearly 50% of the posterior probability of colocalization, an alternative candidate variant in strong LD with rs12967135 explains over 30% of the posterior. Thus, there is no clear evidence in favour of rs12967135 being the ‘true’ causal variant.

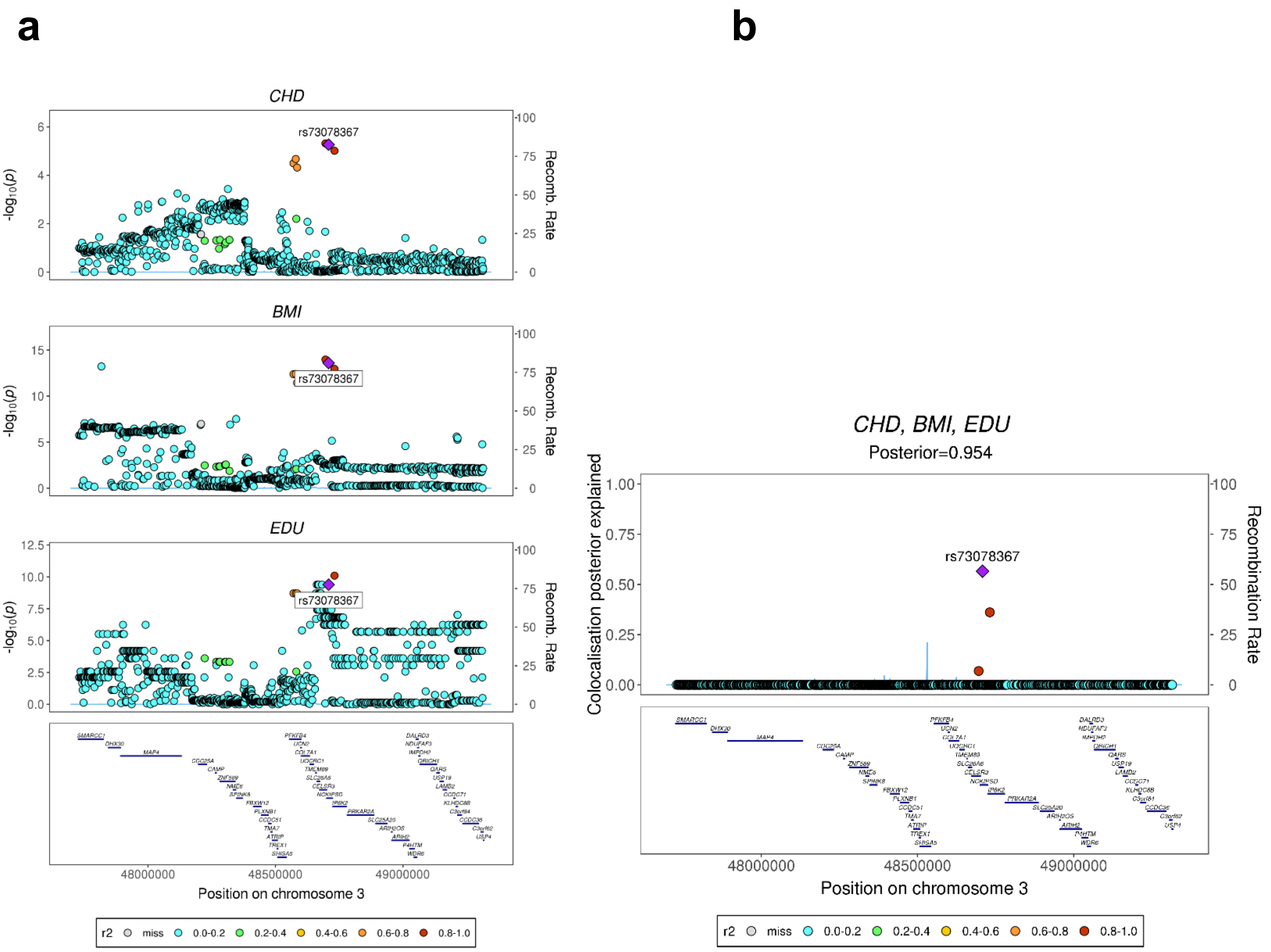

**Figure S8:** Evidence of a single shared causal variant between traits while multiple (within-trait) causal variants are likely. (**a**) Stacked association plots of CHD with body mass index (BMI) and education years (EDU). At least one trait, e.g. BMI, appears to have multiple causal variants in the region (noted by weak LD between the two strongest associated SNPs in BMI). Despite this, HyPrColoc identifies a single shared associated region and within this region implicates a single shared causal variant between the traits (posterior probability of colocalization 0.95). (**b**) The candidate causal variant rs73078367 cannot be clearly distinguished from other possible candidate variants however, i.e. rs73078367 explains around 60% of the posterior probability of colocalization with the next candidate explaining approximately 35%.
